# A global genetic interaction map of a human cell reveals conserved principles of genetic networks

**DOI:** 10.1101/2025.06.30.662193

**Authors:** Maximilian Billmann, Michael Costanzo, Xiang Zhang, Arshia Z. Hassan, Mahfuzur Rahman, Kevin R. Brown, Katherine S. Chan, Amy Hin Yan Tong, Carles Pons, Henry N. Ward, Catherine Ross, Jolanda van Leeuwen, Michael Aregger, Keith A. Lawson, Barbara Mair, Amy F. Roth, Nesli E. Sen, Duncan Forster, Guihong Tan, Patricia Mero, Sanna N. Masud, Yoonkyu Lee, Magali Aguilera-Uribe, Matej Usaj, Sylvia M.T. Almeida, Kamaldeep Aulakh, Urvi Bhojoo, Saba Birkadze, Nathaniel Budijono, Xunhui Cai, Joseph J. Caumanns, Megha Chandrashekhar, Daniel Chang, Ryan Climie, Kuheli Dasgupta, Adrian Drazic, Jose I. Rojas Echenique, Rafael Gacesa, Adrian Granda Farias, Andrea Habsid, Ira Horecka, Kristin Kantautas, Fenghu Ji, Dae-Kyum Kim, Seon Yong Lee, Wendy Liang, Julianne Lim, Kevin Lin, Xueibing Lu, Babak Nami, Allison Nixon, Nicholas Mikolajewicz, Lyudmila Nedyalkova, Thomas Rohde, Maria Sartori Rodrigues, Martin Soste, Eric Schultz, Wen Wang, Ashwin Seetharaman, Emira Shuteriqi, Olga Sizova, David Thomson Taylor, Maria Tereshchenko, David Tieu, Jacob Turowec, Tajinder Ubhi, Sylvia Varland, Kyle E. Wang, Zi Yang Wang, Jiarun Wei, Yu-Xi Xiao, Grant W. Brown, Benjamin F. Cravatt, Scott J. Dixon, Haley D.M. Wyatt, Hannes L. Röst, Frederick P. Roth, Tian Xia, Gary D. Bader, Robbie Loewith, Nicholas G. Davis, Brenda Andrews, Chad L. Myers, Jason Moffat, Charles Boone

## Abstract

We generated a genome-scale, genetic interaction network from the analysis of more than 4 million double mutants in the haploid human cell line, HAP1. The network maps ∼90,000 genetic interactions, including thousands of extreme synthetic lethal and genetic suppression interactions. Genetic interaction profiles enabled assembly of a hierarchical model of cell function, including modules corresponding to protein complexes, pathways, biological processes, and cellular compartments. Comparative analyses showed that general principles of genetic networks are conserved from yeast to human cells. A genetic interaction network mapped in a single genetic background complements the DepMap gene co-essentiality network, recapitulating many of the same biological connections and also capturing unique functional information to reveal roles of uncharacterized genes and molecular determinants of specific cancer cell line genetic dependencies.

## Main Text

Genetic interactions identify functional connections between genes and genetic modifiers that impact the genotype to phenotype relationship (*1*). Negative interactions, such as synthetic lethal or sick interactions, occur when a double mutant shows a fitness defect greater than the expected effect of the combined single mutants. Positive interactions, such as suppression interactions, are scored in double mutants that grow better than expected (*1*). Systematic analysis in the budding yeast, *Saccharomyces cerevisiae*, mapped a global network of ∼1,000,000 negative and positive interactions among its ∼5,000 nonessential and ∼1,000 essential genes (*2–5*). Yeast genetic interactions tend to occur among functionally related genes, and a genetic interaction profile similarity network connects gene pairs with similar sets of genetic interactions. The yeast genetic interaction profile similarity network revealed the functional architecture of a cell by clustering genes into groups of hierarchically organized modules of increasing size that correspond to protein complexes or pathways, biological processes and cellular compartments (*5*).

Genome-wide pooled CRISPR and transposon mutagenesis screens defined a core set of essential genes required for proliferation of most cell lines, as well as selectively essential genes required for growth of specific cancer cell lines to generate a gene-by-cell line dataset highlighting cancer cell-specific genetic vulnerabilities [e.g. The Cancer Dependency Map project (DepMap), depmap.org/portal](*6–14*). A gene-gene co-essentiality network connecting gene pairs that are essential in the same cancer cell lines also provides a view of gene function (*15–19*). Selectively essential genes are relevant to our understanding of genetic interactions because their essential roles may depend on synthetic lethality driven by variation specific to a particular genetic background (*20–22*). However, the underlying genetic mechanisms associated with these dependencies are mostly unknown and likely genetically complex, involving multiple variants and other cell line-specific factors (*1, 23–25*).

In addition to single gene perturbation analyses, combinatorial RNAi and CRISPR based on multiplexing shRNAs and gRNAs, respectively, have identified genetic interactions among subsets of human genes in a single cell line (*26–39*). Larger scale studies used CRISPR interference (CRISPRi) to map genetic interactions by modulating expression of thousands of gene pairs (*40, 41*), and similar methods promise to expand the scale and efficiency of genetic interaction screens in human cells (*33, 42*). Complementary approaches used genome-wide CRISPR or transposon mutagenesis approaches to introduce secondary mutations into engineered cell lines, each carrying a stable ‘query’ mutation of interest (*6, 43–46*). Collectively, these studies, along with the yeast genetic network, suggest that a genome-scale genetic network mapped in a single cell line should organize a large fraction of the human genome into functionally enriched gene modules and highlight the functional architecture of a human cell.

Here, we report analysis of ∼4,000,000 gene pairs to construct a functionally unbiased and genome-scale genetic network consisting of ∼90,000 genetic interactions in the human haploid cell line, HAP1. Like the global yeast genetic network, the HAP1 network is rich in functional information, organizing human genes into hierarchical structured sets of functional modules. The HAP1 genetic network complements the DepMap co-essentiality network, revealing similar roles of previously uncharacterized genes and molecular factors underlying specific cancer cell line genetic dependencies. We conclude that the general principles and topology of genetic networks are conserved from yeast to human cells.

## Results

### HAP1 essential and nonessential fitness genes

To systematically map a human cell genetic network, we chose HAP1 as a model cell line because it lacks aneuploidies and is amenable to loss-of-function (LOF) genetic screens (*6, 44, 45, 47–49*). Quantitative measurement of genetic interactions requires accurate single mutant fitness phenotypes, which we generated by performing 39 genome-wide, pooled CRISPR-Cas9 Knockout (KO) screens with the TKOv3 gRNA library in HAP1 wild-type (WT) cells (Files S1, S2)(*7, 50, 51*). Sequencing and *de novo* assembly confirmed that the HAP1 WT cell line genome largely reflects the reference human genome (hg38), validating use of the TKOv3 library (*7, 50, 51*). We quantified single gene mutant fitness by measuring TKOv3 gRNA abundance within infected HAP1 WT cell populations at regular time intervals, for up to 20 doublings in rich or minimal medium (fig. S1A-C)(*51, 52*). Individual perturbation of ∼22% (3,941/17,724) of library genes significantly altered HAP1 cell fitness (fig. S1C, File S2), and single mutant fitness measurements derived from both growth conditions were highly correlated (fig. S1D), with only a few genes showing significant condition-specific growth phenotypes (∼0.2%, 38 genes, File S2).

We applied a random forest model to our HAP1 WT screen data, trained on a set of cancer cell line core essential genes, to identify 1,524 genes (∼15% of expressed genes) that were essential for HAP1 cell proliferation (fig. S1B-C, Files S2, S3)(*10, 12, 51*), many of which (∼81%, 1231/1524) are also required for viability of most DepMap cancer cell lines. The remaining ∼19% (293/1524) of library genes were specifically essential in HAP1 cells but their perturbation often impacts growth of multiple cancer cell lines (fig. S2A-C)(*10, 51*). In general, HAP1 essential genes encode highly expressed, conserved proteins that span diverse bioprocesses and exhibit physiological and evolutionary properties commonly associated with essential genes identified in other cell lines and model organisms (fig. S2D-G)(*6, 7, 53–56*). We also identified 2,417 nonessential genes that resulted in a reproducible fitness phenotype when disrupted in HAP1 cells including 1,859 genes with a fitness defect and 558 genes with increased fitness relative to WT (fig. S1C, File S2).

### Genome-scale quantitative genetic interaction analysis

We developed a quantitative genetic interaction (qGI) score that compares the abundance of TKOv3 gRNAs derived from CRISPR-based screens in WT HAP1 cells vs. HAP1 query mutant cells carrying a stable mutation in a gene of interest (Fig. 1A)(*44, 45, 51, 52*). The abundance of gRNAs in a query mutant cell line provides an estimate of double mutant fitness (Fig. 1A). Negative interactions identify genes with gRNAs that show significantly decreased abundance in a query mutant relative to WT, whereas positive interactions reflect genes with increased gRNA abundance in a query mutant relative to WT (Fig. 1A-B). We constructed 222 query cell lines, most of which carried a LOF allele of a highly expressed, functionally diverse gene that showed a fitness defect in HAP1 cells. These query cell lines were used in 298 genome-wide screens to score genetic interactions among 3,934,506 unique gene pairs (fig. S1C, S2G, Files S1, S2, S4)(*51*). To determine false negative and positive rates for the qGI score, 7 different query genes were each screened 4-5 times and the results analyzed using a Markov Chain Monte Carlo (MCMC) estimation approach (File S1)(*51, 52*). This strategy generated genome-wide consensus profiles of genetic interactions for each query gene, which were used as a gold standard to estimate precision and recall rates and define optimal qGI score and significance thresholds (|qGI score| > 0.3, FDR < 0.1, fig. S3A, File S5). In total, we identified 88,933 genetic interactions, including 47,052 negative and 41,881 positive interactions (File S4).

**Fig. 1.**
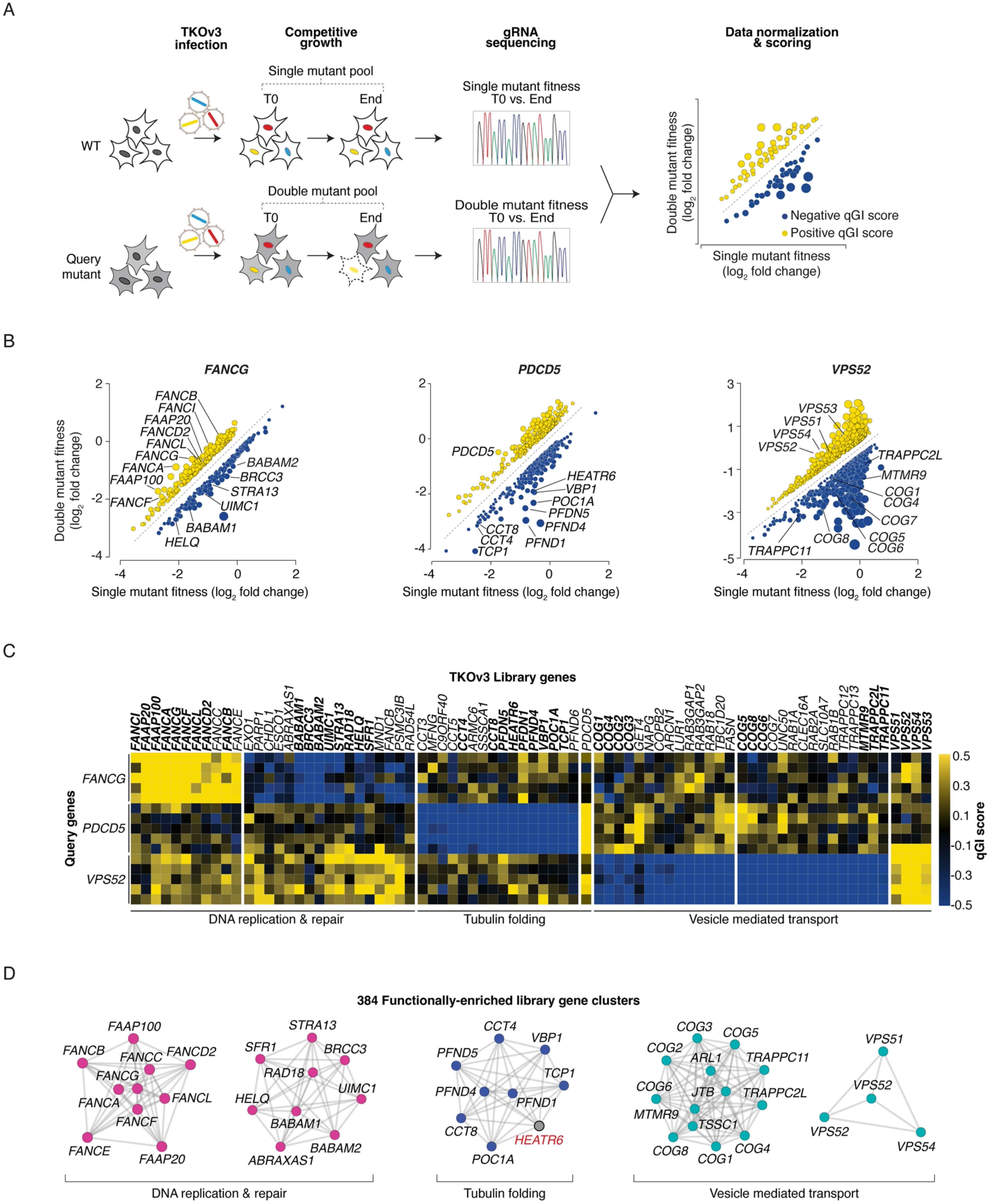
Genome-scale genetic interaction analysis in HAP1 cells. **(A)** Diagram of genetic interaction analysis pipeline in co-isogenic cell lines. The quantitative genetic interaction (qGI) score is based on the difference between log fold change measurements for a given library gene in the query mutant (i.e. double mutant) versus WT (i.e. single mutant) cell populations. **(B)** Scatterplots depicting genetic interactions for the indicated query genes. *FANCG*, *PDCD5*, *VPS52* screen identifiers correspond to GIN192, GIN189, and GIN 241, respectively. Negative (blue) and positive (yellow) genetic interactions that satisfied a standard genetic interaction threshold (|qGI| > 0.3, FDR < 0.1) are shown. Specific negative and positive interactions identified in each screen are indicated. **(C)** Heatmap of qGI values for selected reproducible genetic interactions (columns) from biological replicate screens (n=5) for the indicated query genes (rows). Negative qGI scores are shown in blue and positive qGI scores in yellow. Genes labeled in panel B are indicated in bold face. Functions enriched among specific groups of library genes are indicated. **(D)** Examples of functionally enriched gene modules derived from clustering of the entire genetic interaction dataset, as described (*39*). Node color represents shared general function and the poorly characterized HEATR6 gene is shown in red.

We performed several analyses to assess reproducibility of our genetic interaction measurements. First, replicate screens of an additional 43 query genes (n=2-5, File S1) revealed that double mutant fitness measurements were highly reproducible across replicate query screens (fig. S3B). Replicate correlation of qGI scores depended on the query gene and increased substantially when comparisons were restricted to interactions that satisfied defined score and significance thresholds (fig. S3A-C)(*57*). Second, we observed agreement among reciprocal gene pairs with significant genetic interactions (i.e. query A-library B vs. query B-library A, fig. S3D). Third, replicate genetic interactions measured in both rich and minimal medium were also highly correlated indicating that, like in yeast (*58*), HAP1 genetic interactions were largely robust to environmental differences (fig. S3E). Finally, interactions for 5 query genes were recapitulated using an independent gRNA library, demonstrating that qGI scores were not driven by gRNA-specific phenotypes (fig. S3F, File S6).

We benchmarked negative interactions for the *PTAR1* query gene to those previously identified in a HAP1 transposon-based gene trap screen (fig. S4A, File S7)(*6*). *PTAR1* encodes a geranylgeranyl transferase type III (GGTase III) that modifies and activates YKT6, an essential SNARE (soluble N-ethylmaleimide-sensitive factor activating protein receptor) protein involved in vesicle trafficking (*59*). We identified 316 negative interactions including ∼68% (40/59, *P* < 7×10^-59^, hypergeometric test) of interactions reported by gene trap analysis (fig. S4A, File S7)(*6*). *PTAR1* negative interactions uniquely identified by our study (∼87%, 276/310) were also enriched for vesicle traffic-related genes, suggesting that our CRISPR-knockout (KO) approach is precise and sensitive (fig. S4B, File S7).

### A genetic interaction profile similarity network for a human cell

The genetic interaction profile of a gene represents its unique signature of negative and positive interactions, which reflects its biological function (Fig. 1B). For example, profiles for replicate screens of *FANCG,* a query gene involved in DNA recombination (*60*), comprised genetic interactions with library genes involved DNA replication and repair (Fig. 1B-C). Consistent with its role as a regulator of tubulin polymerization (*61*), *PDCD5* query gene replicate profiles showed negative interactions with genes involved in tubulin function, including Prefoldin and cytosolic chaperonin CCT (Chaperone Containing TCP-1) complex genes, which control actin and tubulin folding (Fig. 1B-C)(*62, 63*). Replicate profiles for *VPS52*, which encodes a component of the GARP tethering complex involved in endosome sorting (*64, 65*), showed many interactions with vesicle trafficking genes (Fig. 1B-C).

Like in yeast (*3–5*), functionally related HAP1 genes belonging to the same pathway or bioprocess shared similar interaction profiles (figs. S5). Hierarchical clustering of the complete HAP1 genetic interaction dataset grouped genes together based on their interaction profile similarity, identifying sets of genes that function together in the same bioprocess, pathway or protein complex (fig. S6, File S8). In total, we identified 412 clusters involving ∼4,400 library genes, most (∼93%, 384/412) of which were enriched for specific Gene Ontology (GO) bioprocess terms that spanned diverse cellular functions (Fig. 1D, File S9)(*51*).

We constructed a genome-scale HAP1 genetic interaction profile similarity network (Fig. 2, File S10)(*51*). Nodes in this network represent library genes, and edges connect pairs of library genes that share similar interaction profiles (Fig. 2A). The distance between connected gene pairs reflects their profile similarity. Proximally located genes share more similar patterns of genetic interactions, while genes positioned farther apart in the network display more divergent profiles (Fig. 2A). The HAP1 network is relatively sparse because library gene profiles are based on genetic interactions with only 222 unique query genes. Nonetheless, ∼74% (2787/3784) of genes on the network belonged to large, discernible network clusters. By applying Spatial Analysis of Functional Enrichment (SAFE)(*51, 66*), with a GO bioprocess functional standard, we identified 17 densely connected network clusters, each enriched for related GO terms corresponding to a different bioprocess, such as DNA replication and repair or vesicle trafficking (Fig. 2B, File S11). Combining SAFE with a protein localization standard (*67*) highlighted seven larger network regions of neighboring bioprocess-enriched clusters that comprised proteins localized to the same subcellular compartment (Fig. 2C, File S11). Bioprocess-enriched network regions were also dissected into smaller subclusters corresponding to 71 protein complexes (Fig. 2D, File S11). Thus, the HAP1 genetic interaction profile similarity network shares a similar topology with the global yeast network (*4, 5*), where genes annotated to the same protein complex share similar patterns of genetic interactions and located next to one another (Fig. 2E-F).

**Fig. 2.**
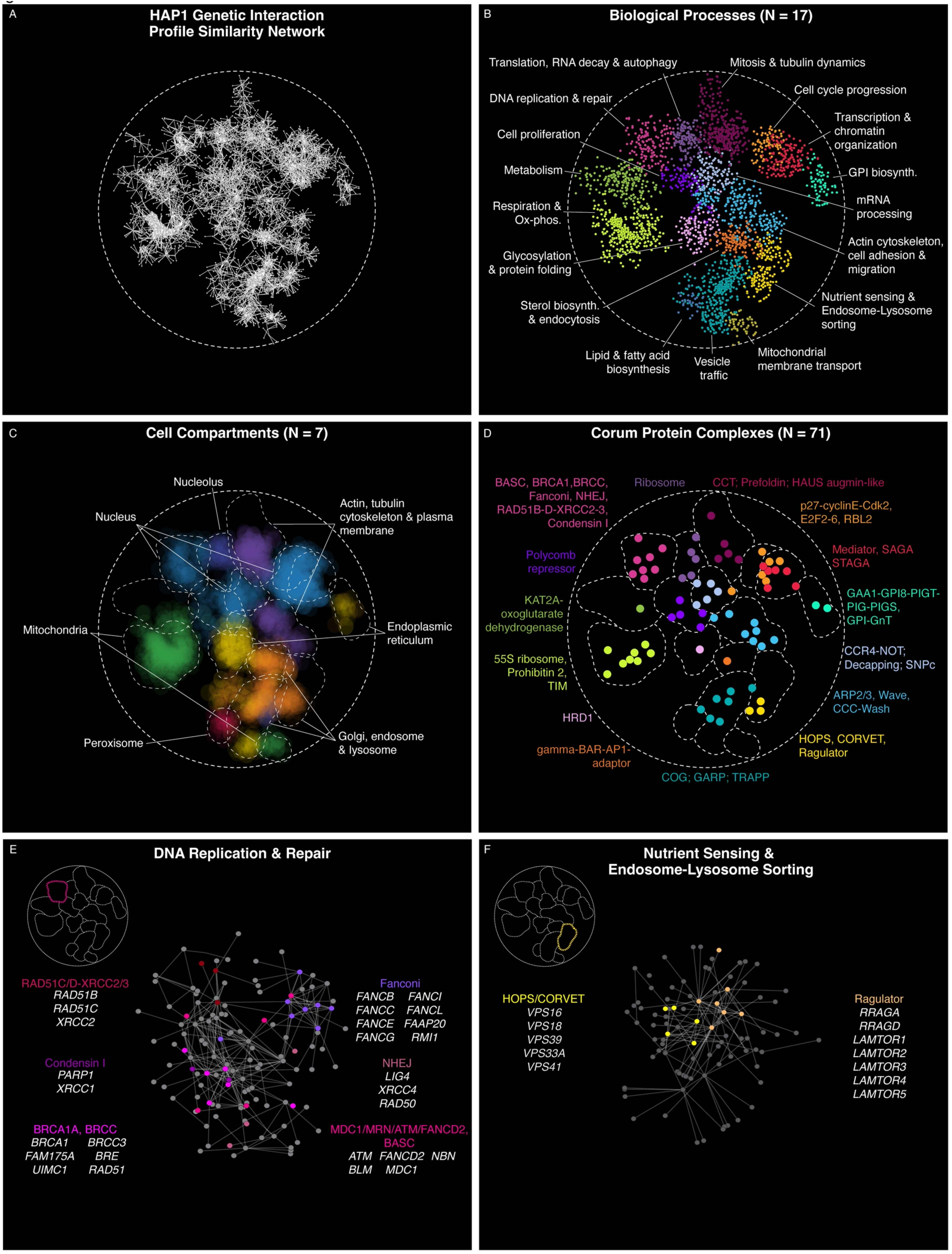
A genetic interaction profile similarity network for a human cell. (A) HAP1 genetic interaction network comprising 3784 human genes (nodes). Gene pairs were connected by profile similarity (PCC > 0.41) and graphed using a spring-embedded layout algorithm (edges)(*161*). Genes sharing similar genetic interaction profiles are positioned near each other, whereas genes with less similar genetic interaction profiles are farther apart. **(B)** HAP1 genetic interaction network annotated using SAFE (*66*) for bioprocess terms. **(C)** HAP1 genetic interaction network highlighting network regions that are enriched for proteins in the same cellular compartment. Node opacity reflects gene-level enrichment significance, with more enriched genes displayed more opaquely. Dashed lines represent network regions enriched for bioprocesses indicated in panel B. **(D)** HAP1 genetic interaction network annotated by highlighting CORUM protein complexes. Nonredundant protein complexes were identified by assigning each gene to the largest complex it belongs to with >2 unique members. The centroid of the network positions of the genes annotated to a given protein complex was used to create the protein complex node. Nodes are colored according to the biological process-enriched region of the network to which they localized. Dashed lines indicate the network regions enriched for bioprocesses indicated in panel B. **(E-F)** Genes belonging to the bioprocess– enriched network region highlighted in the inset were extracted from the HAP1 network and genes (nodes) in the subnetworks were colored according to their Corum protein complex annotation.

### Annotating gene function using the HAP1 profile similarity network

A genetic interaction profile similarity network provides a resource for annotating gene function (*5*). We linked 113 genes associated with relatively few citations and/or GO annotations to specific bioprocesses on the HAP1 profile similarity network (fig. S7A, File S12). For example, the *C1orf112/FIRRMM* profile suggested a role for this gene in DNA damage and repair, a prediction supported by recent studies (fig. S7A-B, File S12)(*46, 68*). Another poorly characterized library gene, *HEATR6*, shared interactions in common with members of the CCT chaperonin and Prefoldin complexes, suggesting that this gene may have a role in actin or tubulin folding (Fig. 1D, Files S9, S12).

Because chemical-genetic interactions often mimic genetic perturbations, the HAP1 profile similarity network provides insights into the mode-of-action of bioactive molecules (*4, 69, 70*). For example, genes that showed sensitivity or resistance to NGI-1, a small molecule inhibitor of the oligosaccharyltransferase (OST) complex (*71*), were enriched for roles in protein glycosylation and vesicle trafficking and localized to the corresponding functional domain region of the genetic profile similarity network (fig. S7C, File S13)(*51*). The HAP1 genetic profile network also highlighted functions shared among different subsets of genes associated with the same disease trait or phenotype (fig. S7D)(*51*). Thus, as first demonstrated in yeast, a network of genetic interaction profiles is rich in functional information, that can be used to predict gene function and discover mechanisms of sensitivity to bioactive compounds or other environmental perturbations.

### Genetic network connectivity

While the number of genetic interactions per library gene ranged between 0 and ∼70, the average library gene interacted with ∼2% of all query genes, exhibiting ∼2-3 negative and ∼2 positive interactions (Fig. 3A, fig. S8A, File S14). Some genes participated in many interactions representing genetic network hubs. The top 5% most connected genes exhibited ∼6-fold more interactions than the average gene (Fig. 3B, File S14). Although hub genes spanned different functions, genes with roles in mitochondrial-related functions were among the most highly connected and highly correlated genes in the HAP1 network, exhibiting numerous interactions, especially many positive interactions (Fig. 2B, figs. S5B, S8C-E, File S14). Gene pairs with strongly correlated co-essentiality profiles, required for fitness of the same set of DepMap cancer cell lines, also predominantly involve mitochondrial-related genes, especially those encoding the electron transport chain (ETC) or the 55S ribosome (*19*). ETC and 55S ribosome proteins are highly stable and detection of a growth phenotype resulting from disruption of these genes may not manifest until the WT protein is depleted. Experimental factors, such as sampling time and cell doubling rate, may impact fitness measurements and increase correlation between co-essentiality profiles of ETC and 55S ribosome genes (*19*). Given the potential for experimental factors to confound scoring of mitochondrial gene interaction profiles, we explored network properties using the complete dataset and a subset of data that excluded mitochondrial-related genes (*51*).

**Fig. 3.**
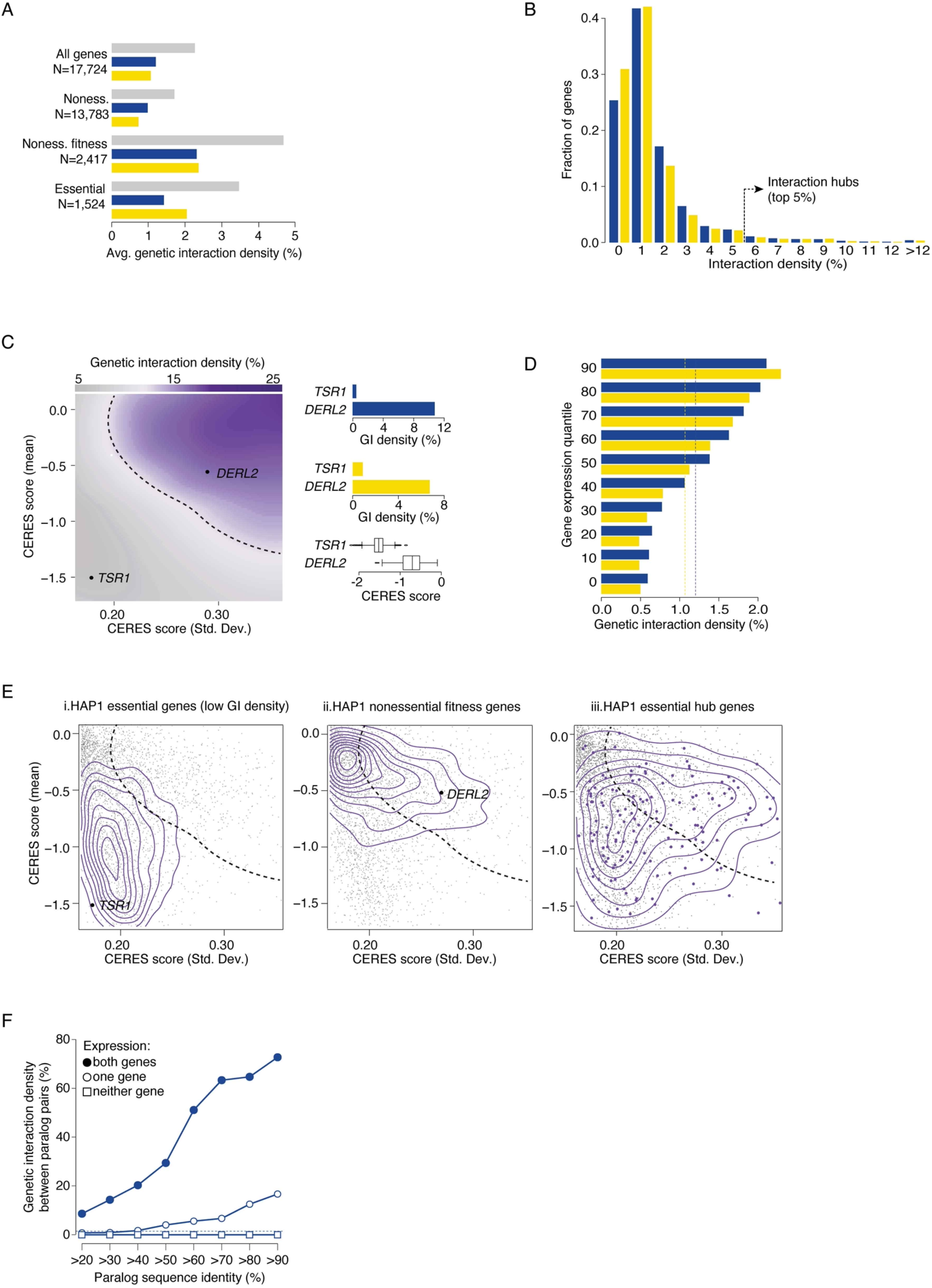
Genetic interaction density analysis. **(A)** Bar chart showing genetic interaction density (observed interactions/total gene pairs screened) for library genes by category (all genes, nonessential (noness), nonessential with fitness phenotypes (noness fitness), essential) at |qGI| >0.3, FDR < 0.1. Negative (blue), positive (yellow) and total (grey) interaction densities, along with the number of genes in each category, are indicated. **(B)** Density distribution of negative (blue) and positive (yellow) interactions, highlighting the top 5% of genes with the highest interaction density. **(C)** Genetic interaction density heatmap visualized as a function of a gene’s single mutant standard deviation in the DepMap dataset (x-axis, CERES score std. deviation) and the single mutant mean phenotype (y-axis, CERES score mean). Darker purple represents increased total genetic interaction density (positive and negative interactions) in the HAP1 GI network. Right bar plots show positive and negative density and CERES score mean for the genes *TSR1* and *DERL2*. The dotted line indicates the boundary between high and low genetic interaction density. **(D)** The average negative (blue) and positive (yellow) interaction density for library genes as a function of expression in HAP1 cells, with dotted lines indicating background interaction densities across all tested library genes. **(E)** The distribution of genes belonging to each gene set is plotted as a function of a gene’s single mutant standard deviation (x-axis, CERES score std. deviation) and mean phenotype (y-axis, CERES score mean) in the DepMap dataset. Plots show (i) HAP1 essential genes with lowest 50% GI density, (ii) HAP1 non-essential genes with significant fitness effects, and (iii) HAP1 essential genes with the top 20% total interaction density (right). The contour lines reflect the density of the corresponding gene sets in this two-dimensional space. The dotted line indicates the boundary between high and low genetic interaction density as defined in C. Grey nodes represent library genes with at least 1 genetic interaction (|qGI| > 0.3, FDR < 0.1). The purple nodes indicate HAP1 essential hub genes. **(F)** Negative genetic interaction density among pairs of duplicated genes with increasing sequence identity (i.e. paralogs).

Many physiological and evolutionary properties characteristic of yeast genetic network hub genes were also associated with high connectivity in the HAP1 genetic network (Fig. 3, figs. S8, S9)(*4, 5, 72*). For example, genetic interaction density (i.e. genetic interactions/gene pairs tested) was related to single mutant fitness in HAP1 cells (Fig. 3A, figs. S8B, S9). Nonessential genes with fitness defects interacted with ∼5% of query genes, or ∼2-fold more interactions relative to the average library gene (Fig. 3A, fig. S8A, File S14). Moreover, nonessential genes with increasingly severe fitness defects exhibited a greater number of negative and positive interactions (fig. S8B).

Genetic network connectivity was also related to single gene mutant fitness measured across the DepMap panel of cancer cell lines (Fig. 3C, fig. S9)(*51*). A mean CERES score reflects the average fitness associated with disruption of a particular gene while CERES score standard deviation indicates the variability of mutant fitness when assessed across multiple cell lines (*10*). Genes with extreme negative mean CERES scores and extremely low CERES score standard deviations, such as the ribosome maturation factor, *TSR1,* represent core essential genes that showed relatively few interactions in the HAP1 genetic network (Fig. 3C). This suggests that CRISPR-based inactivation of many essential genes results in rapid depletion of the corresponding mutant cells in a competitive growth assay, thereby reducing the potential to identify genetic interactions. Nonetheless, genes with less extreme yet increasingly negative CERES scores and higher standard deviations, such as *DERL2*, a gene involved in ER-associated protein degradation, exhibited higher than average genetic interaction density (Fig. 3C, fig. S9)(*73, 74*). Thus, DepMap fitness metrics are predictive of connectivity on the HAP1 genetic network.

Gene expression levels in HAP1 and other cancer cell lines, were also positively correlated with genetic network connectivity (Fig. 3D, fig. S9)(*51*). The most highly expressed HAP1 genes exhibited ∼2-fold more interactions compared to the average gene, while genes ranked in the bottom 40% for expression level in HAP1 cells had fewer interactions, as did genes with variable expression across different cell lines (Fig. 3D, fig. S9). Thus, HAP1 genetic network hub genes tend to be highly and stably expressed across many different cell types. HAP1 hub genes also tend to be evolutionarily conserved and less tolerant of mutations (Fig. 3, fig. S9)(*51*). In total, we identified over 30 different gene features related to genetic interaction density in HAP1 cells (fig. S9).

### Genetic interactions involving essential genes

While it can be difficult to measure genetic interactions for most essential genes, we were able to map interactions for a subset (Fig. 3A, fig. S8A, File S14)(*5*). Most HAP1 essential library genes (∼78%, 1188/1525) exhibited average or below average interaction density, but a fraction of essential genes (∼22%, 336/1524) exhibited above average interaction density and some (∼12%, 175/1524) were among the 5% most highly connected genes in the HAP1 genetic network (File S14). Essential hub genes tend to be expressed at higher levels and encode more abundant proteins compared to other essential genes (fig. S10B)(*51*). Thus, like ETC and 55S ribosome genes discussed above, fitness phenotypes associated with perturbation of essential hub genes may be delayed until levels of the residual WT protein are no longer sufficient to support cellular function. This phenotypic lag may lead to a measurable growth phenotype following gene disruption in the context of a pooled CRISPR assay allowing us to score interactions. Essential hub genes were also enriched for genes predicted to be haploinsufficient in humans and genes that more constrained because they are less tolerant of missense mutations than other essential genes (fig. S10B, File S14).

While most library genes had a similar number of negative and positive interactions, HAP1 essential library genes showed a modest bias towards positive interactions (Fig. 3A, figs. S8A, S10A). This bias may reflect instances of “masking” positive interactions where the expected fitness defect of a query gene mutation is not detected in the context of a more severe fitness defect of an essential library gene (*1, 75*). Alternatively, extreme positive interactions may represent examples of genetic suppression, where a query gene mutation bypasses the essential function of the library gene (*76–79*).

HAP1 essential gene interaction density also distinguished between DepMap core and selectively essential genes. HAP1 essential library genes that exhibited fewer interactions were more likely to be essential in most cancer cell lines (Fig. 3E (i), fig. S10B). Conversely, HAP1 essential library genes that participated in many interactions had an increased tendency to be selectively essential, often displaying a fitness defect that varied across different cancer line genetic backgrounds (Fig. 3E middle and right panel, fig. S10B).

### Genetic interactions involving duplicated genes

Consistent other studies in yeast (*80–84*) and human cells (*33, 85, 86*), paralog gene pairs, where both genes are expressed in HAP1 cells and share at least 20% sequence identity, were over 7-fold (*P* < 4.5×10^-30^, hypergeometric test) enriched for negative interactions, but not for positive interactions (Fig. 3F, fig. S11A). Negative interactions between HAP1 expressed paralogs sharing 90% identity were over 60-fold (*P* < 2.7×10^-16^, hypergeometric test). In contrast, paralog pairs belonging to larger gene families were connected by negative interactions less frequently and exhibited fewer interactions when surveyed across the entire genome (fig. S11B-D). We also observed an asymmetric pattern of interactions among paralogs, where one gene of each duplicate pair showed substantially more negative interactions, many of which showed biases over ∼5-fold (fig. S11E). Like yeast (*80, 81*), negative interaction asymmetry was significantly greater than expected (*P* < 0.01, Empirical P-value)(*51*), suggesting that the paralog with more negative interactions may be under stronger evolutionary constraint (*80, 81*).

### Relating genetic and physical interactions

Gene pairs that are co-expressed and/or whose products physically interact, either as individual protein-protein interactions (PPIs), as part of protein complexes (co-complex) or as part of biological pathways (co-pathway), overlapped significantly with genetic interactions (Fig. 4A)(*51*). For example, gene pairs whose products share a PPI were ∼3-fold enriched for negative interactions and ∼2-fold enriched for positive interactions (Fig. 4A, fig. S12A). While negative interactions involving either nonessential or essential genes overlapped PPIs to a similar extent, the overlap between PPIs and positive interactions depended on gene essentiality (Fig. 4A, fig. S12A). Similar to yeast (*5, 87*), positive interactions between nonessential genes overlapped significantly with PPIs reflecting that simultaneous perturbation of two genes encoding members of the same nonessential pathway or protein complex do not enhance the fitness defect associated with the corresponding single mutants (Fig. 4A, fig. S12A). For example, *VPS52* exhibited within-complex positive interactions with other GARP complex nonessential genes (Fig. 4B). Mitochondrial genes participated in many positive genetic interactions with each other and, while most functional trends were not impacted, an enrichment for positive genetic interactions connecting essential genes whose protein products physically interact was observed when mitochondrial genes were included in the analysis (fig. S12A) but not observed when excluded (Fig. 4A).

**Fig. 4.**
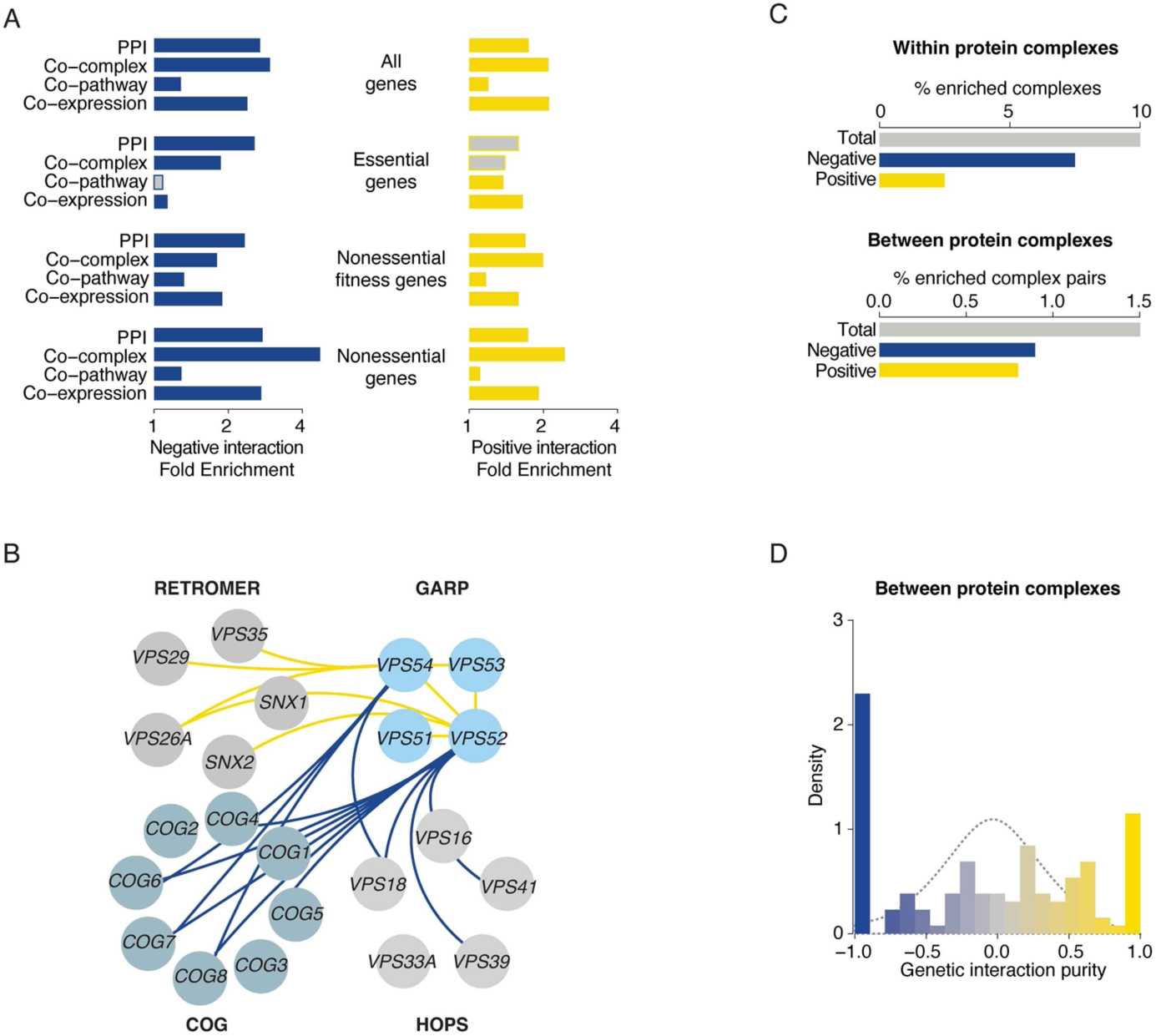
Relating genetic and physical interactions. **(A)** Bar charts indicating significant fold-enrichment (*P* < 0.05, hypergeometric test) for gene pairs encoding physically interacting proteins (PPI), proteins within the same protein complex (Co-complex), proteins within the same pathway (Co-pathway) or co-expressed gene pairs among negative (blue) and positive (yellow) genetic interactions. Enrichment was measured for all gene pairs, essential gene pairs, pairs of nonessential genes associated with a fitness phenotype, and nonessential genes lacking a fitness phenotype. Grey bars indicate non-significant enrichment. Genes with roles in mitochondrial-related functions were excluded from this analysis. **(B)** Network of coherent negative (blue) or positive (yellow) genetic interactions among genes of the RETROMER, GARP, HOPS and COG protein complexes. Node color indicates members of the same protein complex. **(C)** Bar charts depicting the percentage of CORUM protein complexes whose members were enriched for any type of genetic interaction (grey), negative (blue) or positive (yellow) interactions with each other within (top) or between (bottom) protein complexes. Only single complexes and complex-complex pairs with at least 5 tested gene pairs were included in the analysis. Genes with mitochondrial-related functions were excluded from this analysis. **(D)** Distribution of complex-complex pairs with respect to between-complex genetic interaction purity scores (*51*). A score of -1 indicates that genetic interactions occurring between a pair of protein complexes are exclusively comprised of negative interactions whereas a purity score of 1 indicates pairs of complexes connected strictly by positive interactions. The dotted grey line indicates the random expectation based on purity scores generated by sampling negative/positive interaction signs randomly according to a binomial distribution. Genes with mitochondrial-related functions were excluded from the analysis.

We examined negative and positive interactions among members of 40 protein complexes that were represented as both library and query genes (File S15)(*51, 88*). In total, ∼10% (4/40) of protein complexes were enriched for within-complex interactions (Fig. 4C, fig. S12B, File S15). We also identified 124 pairs of complexes (∼1.5%, 124/8250) that were enriched for between-complex interactions (Fig. 4C, fig. S12B, File S15)(*51*). Interactions within a single complex or between different complexes were strongly biased for a single type of genetic interaction, either negative or positive (Fig. 4B, 4D, fig. S12B, File S15)(*51*). For example, negative interactions connected the GARP complex genes*, VPS52* and *VPS54,* with components of the COG (Conserved Oligomeric Golgi) and HOPS (Homotypic Fusion and Protein Sorting) tethering complexes that mediate intra-Golgi and lysosome vesicle trafficking, respectively (Fig. 4B)(*65*). Positive interactions connected the GARP complex to components of the RETROMER complex, which may reflect a shared role in retrograde vesicle trafficking (Fig. 4B)(*89*). In total, we identified 45 protein complex pairs connected by purely negative or purely positive interactions, which mirrors coherent network topology observed in yeast (Fig. 4B, 4D, fig. S12B, File S15)(*5*).

### Functional distribution of negative interactions

Most HAP1 negative interactions occurred among related gene pairs annotated to the same GO bioprocess term, and the density of negative interactions increased with functional specificity of modules mapped in the HAP1 profile similarity network, defined in Fig. 2B-C (Fig. 5A, fig. S13A-C, File S16)(*51*). In total, ∼33% of genes pairs connected by negative interactions shared some degree of functional relatedness, connecting genes with roles in the same compartment, bioprocess, or pathway/complex (Fig. 5B, fig. S13C). Interaction strength provided a quantitative measure of functional relatedness between genes, as stronger negative interactions connected gene pairs with closer functional relationships. Gene pairs encoding members of the same pathway exhibited stronger negative interactions than genes in the same bioprocess, which often showed stronger negative interactions than genes whose products localize to the same compartment (Fig. 5C, fig. S13C)(*51*).

**Fig. 5.**
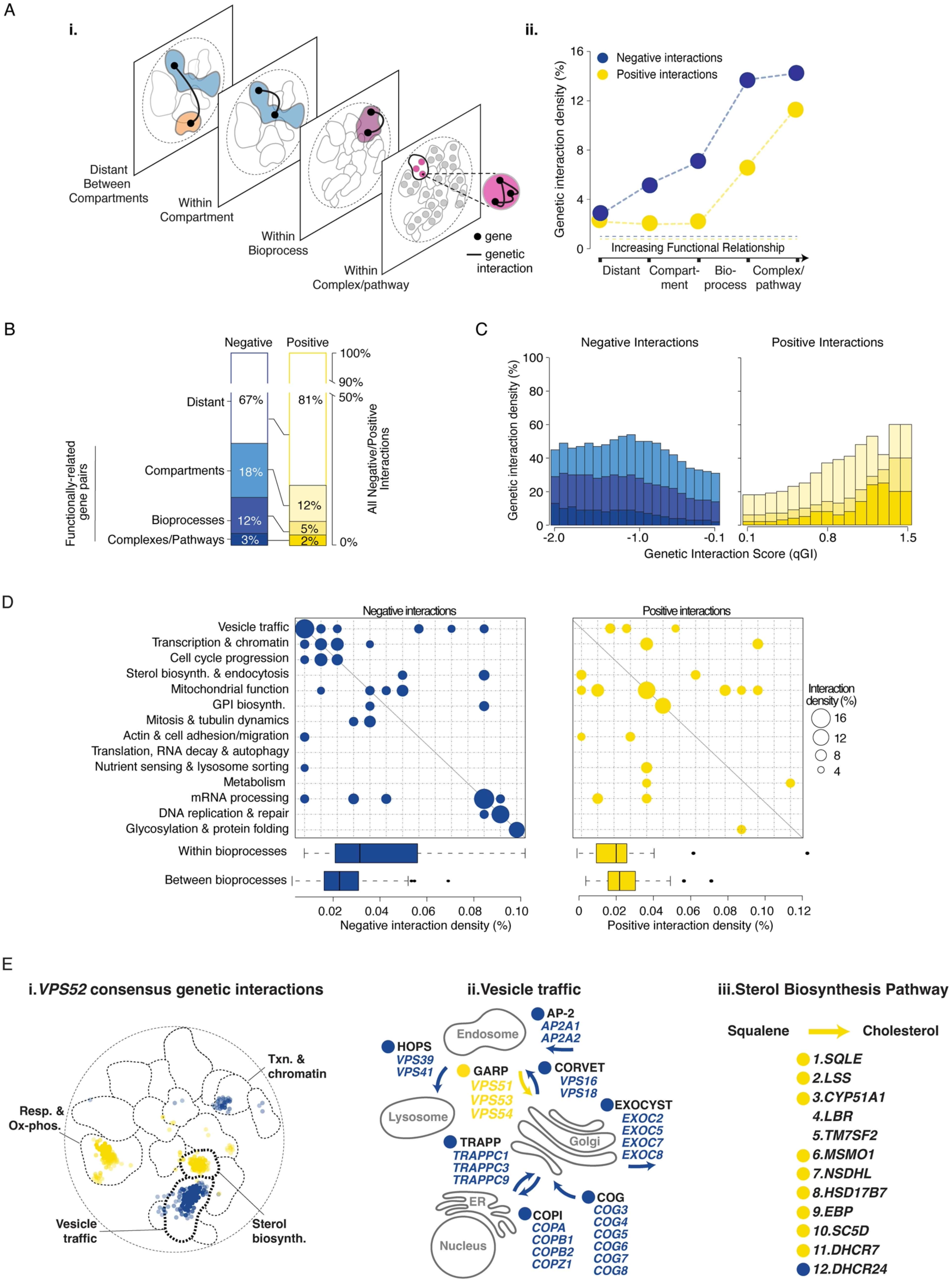
Functional distribution of genetic interactions. **(A)** (i) Schematic of genetic interactions within the functional hierarchy of the HAP1 genetic interaction profile similarity network, showing genetic interactions that occur within the same complex/pathway, biological process, or cellular compartment, and distant interactions between compartments (see Fig. 2). (ii) Line graph showing the observed genetic interaction density for genes within the same hierarchy level for negative (blue) and positive (yellow) genetic interactions (|qGI| > 0.3, FDR < 0.1). Pie charts indicate the total number of gene pairs examined at each level of the functional hierarchy. Horizontal dashed line show background density of negative and positive interactions. Analysis includes ∼1600 genes with high-confidence profiles but excludes mitochondrial-related genes (right). **(B)** Functional distribution of all negative (blue) and all positive (yellow) interactions (|qGI| > 0.3, FDR< 0.1) in the genetic network hierarchy. Genes with mitochondrial-related function are excluded from this analysis. **(C)** Fraction of negative (blue) and positive (yellow) interactions within specified qGI score ranges connecting genes within different functional levels. Different shades of blue and yellow correspond to levels of functional relatedness shown in B. Analysis includes ∼1600 genes with high-confidence profiles, excluding mitochondrial-related genes. **(D)** Network density of genetic interactions (|qGI| >0.3, FDR < 0.1) within and across biological processes for 14 enriched gene sets, as defined in Fig. 1B. Diagonal nodes represent interactions within bioprocesses, off-diagonal nodes represent interactions between bioprocesses. Node size reflects the fraction of interacting gene pairs. The average density of negative and positive interactions observed within and between bioprocesses is shown in the box plots. **(E)** (i) Network map showing regions of the HAP1 profile similarity network enriched for genes with negative (blue) or positive (yellow) consensus genetic interactions with a *VPS52* query gene (n=5 biological replicates). (ii) Genes encoding members of vesicle tethering complexes showing coherent genetic interactions with *VPS52*. (iii) Most genes with roles in the cholesterol biosynthesis pathway show positive interactions with the *VPS52* query gene.

We further examined connections within and between bioprocess modules (*51*). While negative interactions were enriched between genes spanning different bioprocess-enriched clusters (Fig. 5D off-diagonal, File S16), stronger enrichment was observed among genes within the same bioprocess (Fig. 5D on-diagonal, File S16). Although sparse, the connectivity of negative interactions observed within and between human cell bioprocesses resembles that observed in the global yeast network when restricted to an equivalent number of query genes (fig. S13D)(*5*). In general, the functional distribution of negative interactions observed in HAP1 cells mirrors those in the yeast network, highlighting the conservation of genetic network properties.

### Functional distribution of positive interactions

A subset of positive interactions involved “self” interactions where a library gene was targeted in a query cell line carrying a mutation in the same gene (fig. S13E)(*51*). These score as positive interactions because a cell line carrying a LOF allele in a query gene does not show an additional fitness defect when the same gene is targeted by the TKOv3 library. The strength of the positive self-interaction was inversely correlated with single mutant fitness, reflecting that fitness of the query gene mutant is usually not exacerbated by additional genetic perturbation. Thus, most HAP1 query genes (∼94%, 209/222) likely harbored complete LOF mutations. However, a few query genes (∼6%, 13/222) exhibited negative self-interactions, suggesting that these cell lines carried partial LOF query genes mutations (fig. S13E)(*90*).

As for negative interactions, positive interactions also connected functionally related gene pairs annotated to the same GO bioprocess, and the highest density and strongest positive interactions were observed among genes in the same pathway or complex (Fig. 5A, 5C, fig. S13A-C, File S16)(*51*). In particular, pairs of genes involved in mitochondrial- or GPI biosynthesis-related functions were enriched for positive interactions (Fig. 5D on diagonal, File S16). However, only ∼19% of all genes pairs connected by positive interactions shared some degree of functional relatedness as defined by the HAP1 genetic profile similarity network (Fig. 5B, fig. S13C, File S16). Like yeast (fig. S13D)(*5*), most HAP1 positive interactions connected pairs of genes that function in different bioprocesses (Fig. 5D off diagonal, File S16)(*51*).

These genetic interaction trends were exemplified by the *VPS52* genetic interaction profile. While *VPS52* negative interactions were enriched for related genes involved in vesicle trafficking (∼3X, hypergeometric test, Benjamini-Hochberg-corrected FDR < 5.3 x 10^-43^), *VPS52* positive interactions were enriched for genes annotated to different functions, such as sterol biosynthesis (∼3X, hypergeometric test, Benjamini-Hochberg-corrected FDR < 2.4×10^-18^, Fig. 5E). Studies in yeast, human cells, and mouse models show that GARP complex function is connected to cholesterol homeostasis (*91–93*). GARP sorts and localizes the Niemann-Pick Type C disease-associated protein, NPC2, to the lysosome where it exports low-density lipoprotein-derived cholesterol out of lysosomes (*94, 95*). NPC2 is missorted in GARP-deficient mutants, leading to lysosomal accumulation of cholesterol (*92*). Niemann-Pick Type C disease is an autosomal recessive neurodegenerative disorder characterized, in part, by cellular accumulation of cholesterol (*94, 95*). Consistent with these findings, *VPS52* exhibited strong positive interactions with most genes in the cholesterol biosynthetic pathway (Fig. 5E, fig. S14A). One exception was a negative interaction with *DHCR24*, suggesting that defects in *DHCR24* function results in accumulation of a toxic metabolite in *VPS52* mutant cells (Fig. 5E).

Most cholesterol genes were classified as suppression interactions because relevant double mutant fitness was greater than the fitness of the *VPS52* query mutant (Fig. 6A, fig. S14A, File S17). Similar suppression interactions were observed with *VPS54* (fig. S14A, File S17). This suggests that fitness defects associated with GARP complex disruption may result from toxic accumulation of cholesterol, which can be suppressed by reduced cholesterol biosynthesis. GARP mutant fitness defects were also suppressed by disruption of sphingolipid biosynthesis genes (Fig. 6A, fig. S14A-B, File S17). Mutations in GARP complex genes, *VPS53* and *VPS54*, have been linked to cerebello-cerebral atrophy type 2 and Amyotrophic Lateral Sclerosis, respectively (*96–98*). Both neurodegenerative disorders are characterized by cellular accumulation of sphingolipid intermediates and inhibition of sphingolipid biosynthesis rescues mutant phenotypes in relevant disease models (*91, 93*).

**Fig. 6.**
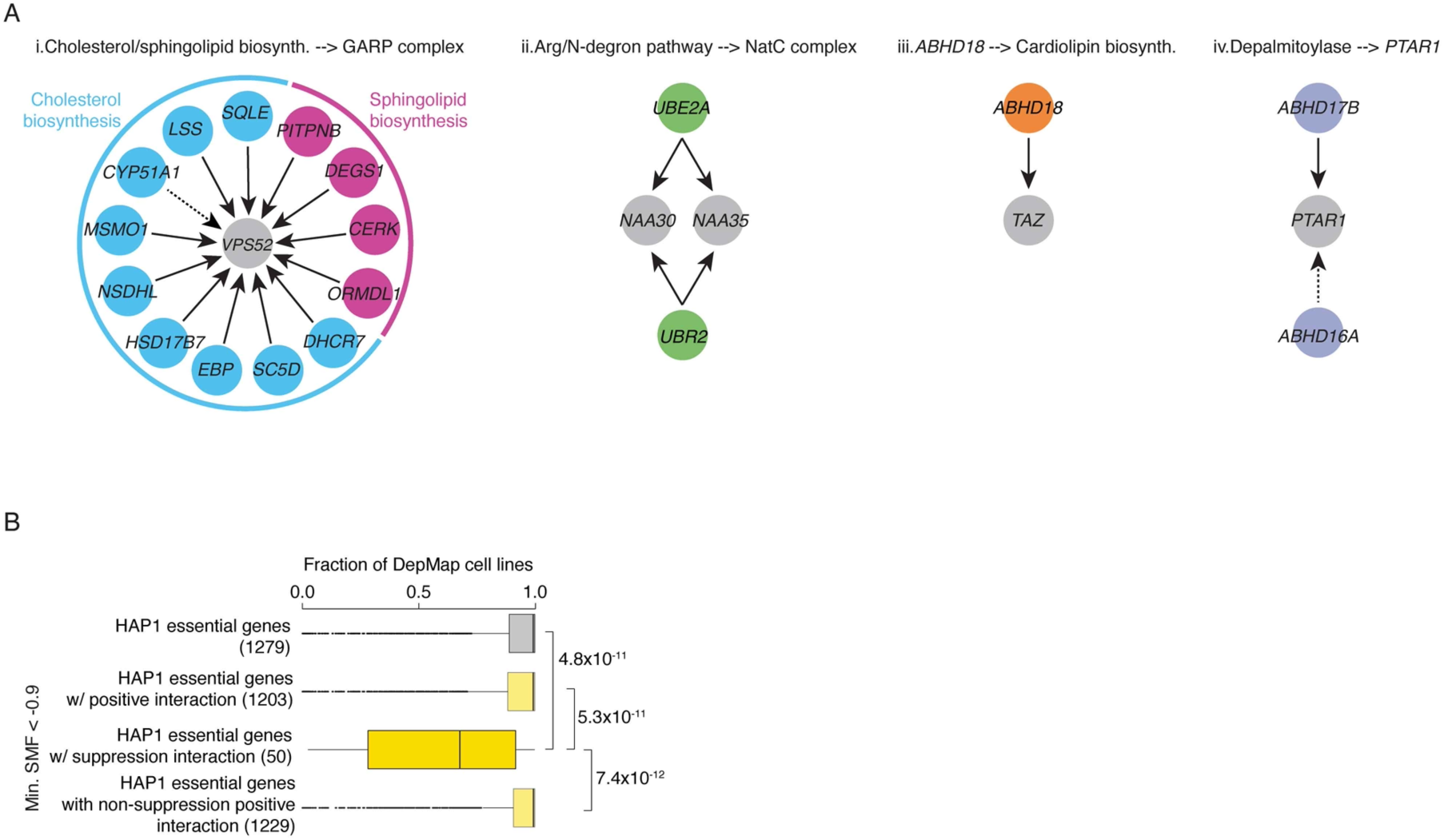
Genetic suppression interactions. **(A)** Specific examples of genetic suppression. Arrows indicate direction of suppression. Grey nodes indicate genes whose mutant fitness phenotype is suppressed and colored nodes represent suppressor genes. Dotted arrow indicates weak suppression interactions that did not satisfy a suppression score threshold (score >0.5). *VPS52* suppressors involved in cholesterol biosynthesis (blue) and sphingolipid biosynthesis (pink) are highlighted **(B)** Box plot showing the mean percentage of DepMap cell lines that depend on the indicated groups of HAP1 essential genes for viability. Numbers of essential genes tested in each group are indicated. Genes with mitochondrial-related functions were excluded from this analysis.

### Genetic suppression

Extreme positive suppression interactions often provide mechanistic insight into gene function (Fig. 6A, fig. S14A-C, File S17)(*76, 79*). In addition to the GARP suppression interactions described above, our HAP1 suppression interactions also revealed novel functional interplay between the NatC protein acetylation and the Arg/N-degron pathway (*44*) and identified a new component of cardiolipin biosynthesis, *ABHD18,* whose inactivation rescued Barth Syndrome disease gene (*TAFAZZIN*)-associated phenotypes (*99*).

To systematically examine the prevalence of suppression interactions, we compared double and single mutant fitness phenotypes for all gene pairs, including both essential and nonessential genes, connected by a positive interaction in the HAP1 genetic network (*51*). In total, ∼4% (1843/41881) of positive interactions represent potential suppression interactions where the double mutant fitness was greater than the fitness of the sickest single mutant (fig. S14A-C, File S17). Consistent with systematic suppression studies in yeast (*76, 79*) and reported suppression interactions among human genes (*78*), HAP1 suppression interactions were more functionally informative than positive interactions in general (fig. S14D). Potential extreme suppressor interactions were ∼2-fold more enriched for functionally related genes pairs annotated to the same GO biological process compared to positive interactions not classified as suppression (fig. S14D).

Previously, we showed that ∼17% of yeast essential genes can be rendered dispensable by spontaneous extragenic bypass suppressor mutations, and these dispensable essential genes were more likely to be nonessential in different yeast species compared to core essential genes (*76*). Despite only screening 222 query genes, ∼3% (48/1524) of HAP1 essential library genes exhibited at least one potential bypass suppression interaction (File S17)(*51*). Notably, this subset of HAP1 essential genes tend to be essential in fewer cancer cell lines and, thus, more likely to be classified as DepMap selective essential genes (Fig. 6B, fig. S14E)(*51*).

### Conservation of genetic network structure and topology

Genetic interactions can be conserved, especially at the level of network structure (*72, 100–104*). To more deeply explore conservation of network topology, we compared genetic interaction density within and between bioprocess functional modules from the yeast and HAP1 profile similarity networks (*51*). Remarkably, we found that the density of negative interactions both within individual bioprocess modules and between similar pairs of biological processes modules was significantly correlated, indicating that functional connectivity was conserved from yeast to human cells, a finding that is independent of sequence conservation among the gene sets (Fig. 7A, File S18).

**Fig. 7.**
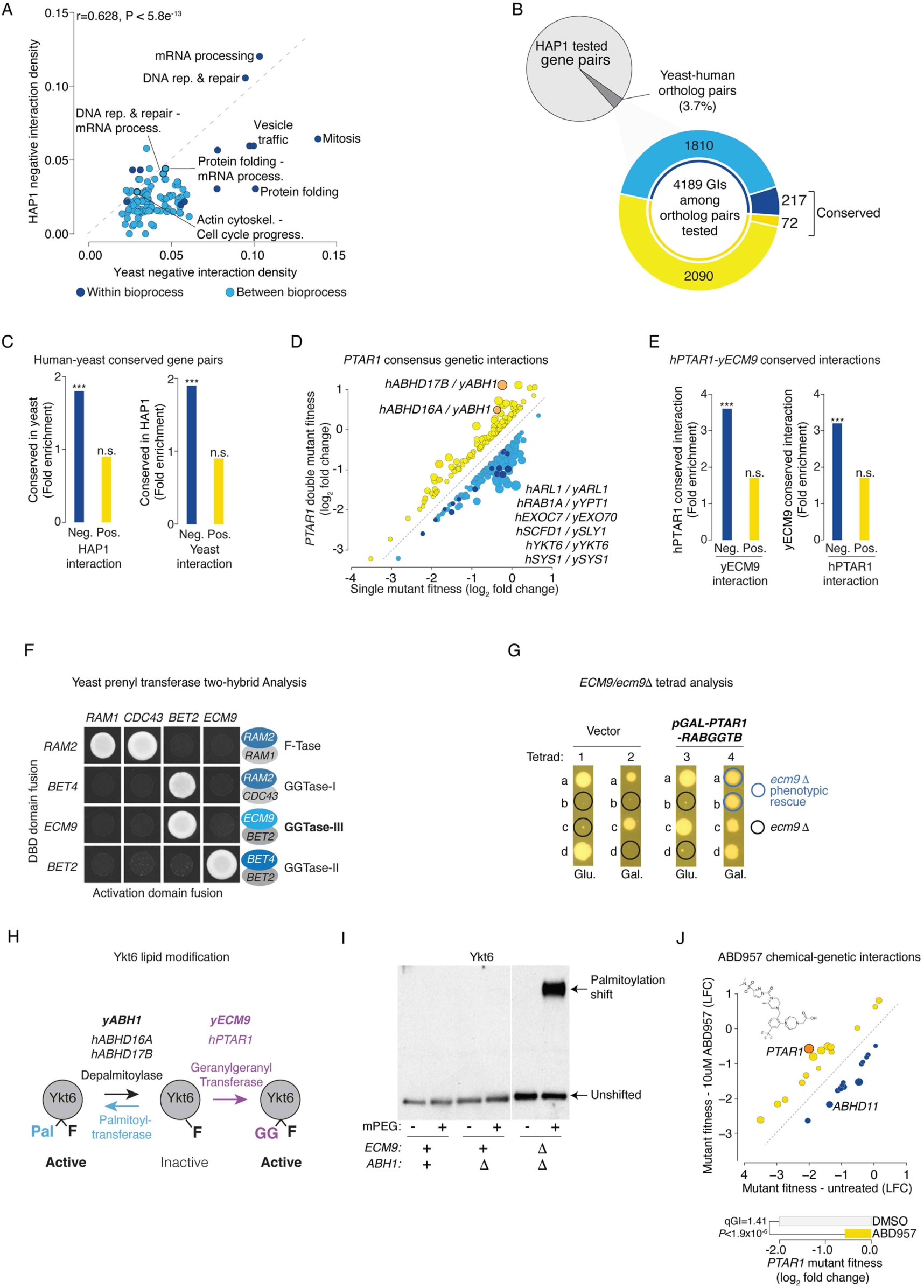
Genetic interaction conservation. **(A)** Scatter plot comparing human (qGI < -0.3, FDR <0.1) and yeast (SGA score < -0.08, *P* <0.05) negative interaction densities within bioprocesses (dark blue) and between pairs of bioprocesses (light blue). **(B)** Pie chart shows the fraction of conserved gene pairs tested in HAP1, while the donut plot summarizes negative and positive interactions in HAP1 among conserved gene pairs. Dark blue and dark yellow represent conserved negative and positive interactions, while light blue and light yellow indicate interactions found only in HAP1. **(C)** Bar graph illustrating enrichment for negative (blue) and positive (yellow) interactions in yeast among conserved gene pairs that showed a negative or positive genetic interaction in HAP1 cells (left). Bar graph illustrating enrichment for negative (blue) and positive (yellow) interactions in HAP1 cells among conserved gene pairs that showed a negative or positive genetic interaction in yeast (right). * indicates level of statistical significance (*** *P* < 10^-6^, Fisher’s exact test). **(D)** Consensus genetic interactions for *PTAR1*. Mean negative (blue) and positive (yellow) qGI scores (|qGI| > 0.3 and FDR < 0.1) based on genetic interactions from 4 independent *PTAR1* genome-wide screens are shown. Conserved negative (dark blue) and positive (orange) genetic interactions identified in HAP1 and yeast screens using human *PTAR1* and yeast *ECM9* orthologous query genes are shown and specific examples of conserved interactions are indicated. **(E)** Bar graphs illustrating enrichment for *PTAR1* negative (blue) and positive (yellow) genetic interactions in HAP1 cells among conserved gene pairs that showed a negative or positive genetic interaction with yeast *ECM9*, and vice versa. * indicates level of statistical significance (*** *P* < 10^-5^, Fisher’s exact test). **(F)** Yeast-two hybrid analysis illustrating the physical interaction between α and β subunits of the indicated prenyltransferases. **(G)** Tetrad analysis showing that co-expression of the human *PTAR1-RABGGTB* GGTaseIII complements essentiality of yeast *ECM9*. Yeast *ECM9/ecm9*11 heterozygous deletion strains carrying a vector control or a plasmid expressing human *PTAR1* and *RABGGTB* expressed from a bidirectional galactose-inducible promoter were sporulated. The meiotic progeny derived from four tetrads were dissected and tested for spore germination (denoted a-d) on either glucose (Glu.) medium, where the promoter is repressed or galactose medium (Gal.) where the promoter is induced. Black circles indicate spore progeny that are predcited to carry the *ecm*911 deletion. Blue circles indicate *ecm911* deletion mutants where the *ECM9* essential phenotype is rescued by galactose-inducible expression of the human *PTAR1-RABGGTB* GGTaseIII (bold). **(H)** Schematic model for dual lipid modification-dependent activation of Ykt6. **(I)** Immunoblot for Ykt6 palmitoylation assessed by mPEG replacement chemistry using protein extracts from the three indicated yeast strains (*51*). **(J)** Chemical-genetic interaction profile mapped for the depalmitoylase inhibitor ABD957. Negative (blue) and positive (yellow) chemical-genetic interactions that satisfied a standard confidence threshold are shown, with select genes highlighted. The chemical structure of ABD957 is shown. Bar graph shows *PTAR1* mutant fitness in ABD957 and DMSO conditions.

Positive interaction density was not correlated between yeast and HAP1 networks (fig. S15A, File S18). In yeast, most essential gene positive interactions do not share a direct functional relationship but rather capture more general regulatory connections related to mRNA degradation or protein turnover (*5*). Positive interactions observed for LOF alleles of HAP1 essential genes may also represent regulatory relationships, including those specific to human cell proliferation. Indeed, genetic interaction profiles associated with tumor suppressor genes were biased towards positive interactions (fig. S10A, File S19), which may reflect the ability of these genes to increase cell proliferation when disrupted in numerous query mutant cell lines. Genes involved in mTOR signaling were also positive interaction hubs, highlighting that disruption of master regulatory pathways can modulate fitness phenotypes associated with disruption of many genes (fig. S8E, File S14). Interestingly, HAP1 genetic interaction profiles identified query mutant cell lines that exhibited specific dependencies on either mTORC1 or mTORC2 signaling pathways, relationships we confirmed by monitoring mTORC1 and mTORC2 activity in specific query mutant cell lines (fig. S15B-E, see Supplementary text).

### Conservation of genetic interactions

Approximately 4% of HAP1 gene pairs tested for genetic interactions (146,664/3,934,728) have a corresponding pair of yeast orthologs (Fig. 7B, File S20)(*51*). Conserved gene pairs that showed a negative interaction in HAP1 cells were more likely to show a negative interaction in yeast and vice versa, indicating that negative interactions are significantly conserved (Fig. 7C, fig. S16A-B). In particular, gene pairs with roles in vesicle trafficking, mitosis, and DNA replication and repair were enriched for conserved negative interactions (fig. S16C). In contrast, orthology was not predictive of positive interactions (Fig. 7B, fig. S16A-B). Of the genetic interactions identified between conserved gene pairs in HAP1 cells, ∼7% (289/4189) were also observed in yeast indicating that a subset of genetic interactions is conserved over ∼1 billion years of evolution (Fig. 7A, File S20).

Orthologous gene pairs that exhibited similar interaction profiles in HAP1 cells also tend to have highly correlated interaction profiles in yeast (fig. S17A)(*51*). For example, *PTAR1* interactions mapped in HAP1 cells overlapped significantly with interactions of the yeast *YKT6,* conserved SNARE protein (fig. S17B). As mentioned above, PTAR1 binds RABGGTB to form a heterodimeric GGTase-III, which activates target proteins, such as YKT6 (*59, 105, 106*). Sequence alignment identified a previously uncharacterized yeast essential gene, *ECM9*, as a distant ortholog of human *PTAR1* (*107, 108*). We generated a temperature-sensitive allele of yeast *ECM9* and screened it for genetic interactions (File S20)(*51*). Yeast *ECM9* negative interactions were also enriched for vesicle trafficking genes, several of which are conserved in human cells, such that *ECM9* and *PTAR1* negative interaction profiles overlap significantly (Fig. 7D-E, fig. S17B-D, File S20). The yeast genetic interaction profile of the RABGGTB ortholog, *BET2,* also overlapped with the *PTAR1* interaction profile (fig. S17B). Two-hybrid analysis showed that yeast Ecm9 interacted specifically with yeast Bet2 and dual expression of human *PTAR1* and *RABGGTB* complemented the lethality of an *ecm9Δ* deletion allele (Fig. 7F-G)(*51*). Moreover, recent mass spectrometry analysis identified Bet2 in Ecm9 immunoprecipitates (*109*).

Although the overlap between yeast *ECM9* and human *PTAR1* positive interactions was not significant (Fig. 7E, fig. S17D), some positive interactions were conserved and biologically informative (Fig. 7D, File S20). In particular, the fitness defect of a *PTAR1* query mutant cell line was suppressed by LOF mutations in either *ABHD16A* or *ABHD17B*, which encode abhydrolase proteins that appear to function as depalmitoylation enzymes (Fig. 6D, Fig. 7D, fig. S14C, File S17)(*110*). Analogously, we previously showed that *ECM9* could be suppressed by disruption of yeast *ABH1* (*YNL320W*), which also encodes a potential deplamitoylation enzyme (*76, 107*). Expression of human *ABHD16A*, *ABHD17B*, or other members of this gene family, rescued the bypass suppression phenotype of an *ecm911 abh111* yeast double mutant, suggesting that several human abhdrolase genes are functional orthologs of yeast *ABH1* (fig. S17E)(*51*).

GGTase-III transfers a geranylgeranyl group to a mono-farnesylated form of yYkt6/hYKT6 to generate a dual prenylated and active SNARE protein (*59, 109, 111*). Our findings imply that in the absence of GGTase-III, farnesylated yYkt6/hYKT6 can be palmitoylated, generating an alternative dual lipid-modified and active SNARE. Mutations in depalmitoylase genes, such as yeast *ABH1*, human *ABHD16A* or *ABHD17B,* which negatively regulate palmitoylation, may promote yYkt6/hYKT6 activation (Fig. 7H). Indeed, yeast Ykt6 was palmitoylated in an *ecm911 abh111* double mutant but not in *abh1Δ* single mutant or WT cells, suggesting that palmitoylation activates Ykt6 in the absence of GGTase-III (Fig. 7I)(*51*). Moreover, a chemical-genetic screen in HAP1 cells using ABD957, a small molecule inhibitor of *ABHD17* depalmitoylases, identified a strong positive chemical-genetic interaction with *PTAR1,* suggesting that chemical inhibition of *ABHD17B* enzyme activity *in vivo* suppresses the fitness defect associated with a *PTAR1* mutant (Fig. 7J, File S13)(*51, 112*). ABD957 also showed a negative chemical-genetic interaction with *ABHD11*, which encodes a potential depalmitoylase that may be functionally related to *ABHD17* depalmitoylases (Fig. 7J, File S13, fig. S17E). Our analysis of *PTAR1*/*ECM9* genetic interactions illustrates the power of a comparative functional genomics approach to reveal new regulators of conserved bioprocesses.

### Deciphering molecular mechanisms underlying cancer gene dependencies

*PTAR1* LOF fitness phenotypes, measured across different DepMap cancer cell lines (depmap.org/portal), were related to the expression level of specific genes that showed genetic interactions with *PTAR1*. For example, cancer cell lines that are more dependent on *PTAR1* for growth tend to express *YKT6* at lower levels reflecting the *PTAR1-YKT6* negative interaction observed in HAP1 cells and supporting a role for PTAR1 as a YKT6 activator (Fig. 8A). Consistent with a *PTAR1-ABHD16A* positive interaction, *PTAR1*-dependent cell lines often expressed *ABHD16A* at higher levels, suggesting that ABHD16A antagonizes PTAR1 function (Fig. 8A). Indeed, *YKT6* and *ABHD16A* expression levels were the most predictive features of *PTAR1* cancer cell line dependency (depmap.org/portal).

**Fig. 8.**
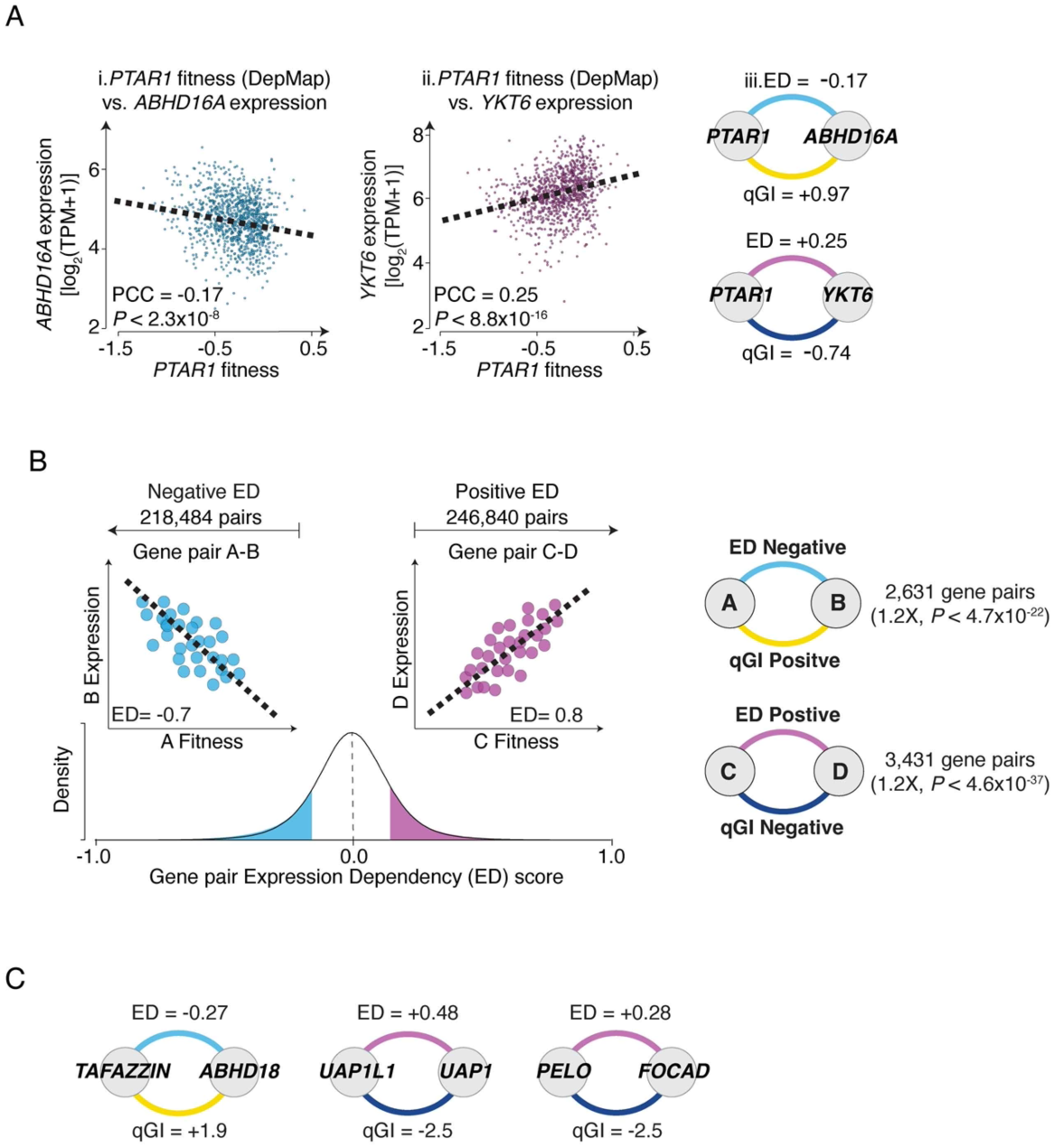
Relationship between DepMap cancer cell line expression dependency and HAP1 genetic interactions. **(A)** Scatter plots illustrating the relationship between *PTAR1* single mutant fitness and expression of either (i) *ABHD16A* or (ii) *YKT6* across DepMap cancer cell lines. (iii) The *PTAR1-ABHD18* gene pair shows a negative Expression-Dependency (ED) score and a positive genetic interaction score (qGI). The *PTAR1-YKT6* gene pair shows a positive ED score and a negative qGI score. **(B)** Schematic illustrating the distribution of ED scores for all gene pairs tested in this study and the overlap between ED and qGI scores. **(C)** Selected examples of specific gene pairs that exhibited significant ED-qGI combinations.

To systematically examine the relationship between cancer cell line genetic dependencies and HAP1 genetic interactions, we generated an Expression Dependency (ED) score, which represents the correlation between the LOF fitness phenotype and expression for gene pairs across a panel of DepMap cell lines (File S21)(*51*). A negative ED score (-ve ED) indicates that the fitness associated with LOF of Gene A is negatively correlated to the expression level of Gene B (Fig. 8B). A positive ED score (+ve ED) occurs when the fitness phenotype of one gene is positively correlated to the expression level of a second gene (Fig. 8B). In total, we computed ED scores for ∼4 million gene pairs tested for genetic interactions in our HAP1 network. After filtering for effect size and significance (|ED| > 0.1, *P* < 0.01)(*51*), we identified ∼218,000 gene pairs with a significant -ve ED and ∼247,000 gene pairs with a significant +ve ED scores (Fig. 8B, fig. S18A, File S21).

Approximately 1% (∼2,600/∼218,000) of gene pairs with extreme -ve ED scores were enriched for a HAP1 positive genetic interaction (1.2-fold, *P* < 4.7×10^-22^, hypergeometric test, Fig. 8B, fig. S18A, File S21), including *PTAR1*-*ABHD16A* and *TAFAZZIN-ABHD18,* which we classified and validated as suppression interactions (Figs. 6A, 8A, 8C, figs. S14C, S18B, Files S17, S21)(*99*). Many gene pairs involving the *TP53* tumor suppressor gene were also associated with a -ve ED and a positive genetic interaction (File S21). Notably, genes that a -ve ED and positive interaction with *TP53* were more frequently co-mutated with *TP53* in various cancers suggesting that disruption of these genes may enhance cancer cell phenotypes associated with *TP53* perturbation (fig. S18C)(*51*).

Approximately ∼1% (∼3,400/∼247,000) of gene pairs exhibiting an extreme +ve ED score were enriched for negative interactions in HAP1 cells (1.2-fold, *P* < 4.6×10^-37^, hypergeometric test, Fig. 8B, fig. S18A, File S21). Consistent with previous studies (*85, 113, 114*), duplicated genes often showed +ve ED scores highlighting the ability of paralogs to compensate for one another (fig. S18A). Gene pairs with both +ve ED and negative interaction scores were over 100-fold more enriched for paralogs compared to gene pairs with a +ve ED score alone (fig. S18A). For example, *UAP1* and *UAP1L1* which encode enzymes that catalyze the last step of uridine diphosphate-N-acetylglucosamine (UDP-GlcNAc) biosynthesis (*115, 116*), exhibited a strong +ve ED and extreme negative interaction scores in HAP1 indicating that these genes impinge on the same essential function (Fig. 8B-C, fig. S18B). The *PELO*-*FOCAD* gene pair also exhibited a strong +ve ED and HAP1 negative interaction, supporting recent studies that identified a synthetic lethal relationship between *PELO* and 9p21.3 deletions involving *FOCAD* (Fig. 8B-C, fig. S18B)(*117, 118*).

### Integrating genetic interaction and co-essentiality networks

Like genetic profile similarity networks, the DepMap co-essentiality network identifies genes that work together in functional modules (*4, 5, 15, 18*). We compared functional information captured by our HAP1 genetic profile similarity network and the DepMap co-essentiality network. To normalize for network size, we constructed 10 DepMap co-essentiality subnetworks, each consisting of 298 randomly sampled DepMap cell lines, which equals the number of genome-wide screens used to construct the HAP1 profile similarity network (*51*). We measured the overlap between pairwise combinations of DepMap co-essentiality subnetworks at varying similarity thresholds (Fig. 9A). At the most stringent thresholds (PCC > 0.4), we observed a relatively high overlap (Jaccard index of ∼0.5) indicating that each DepMap co-essentiality subnetwork contains similar functional information (Fig. 9A). In contrast, DepMap co-essentiality subnetworks showed substantially lower overlap (Jaccard index < 0.1) with the HAP1 profile similarity network, even at stringent similarity thresholds (Fig. 9A). Thus, DepMap coessentiality subnetworks and our HAP1 genetic profile similarity network contain a substantial amount of orthogonal functional information.

**Fig. 9.**
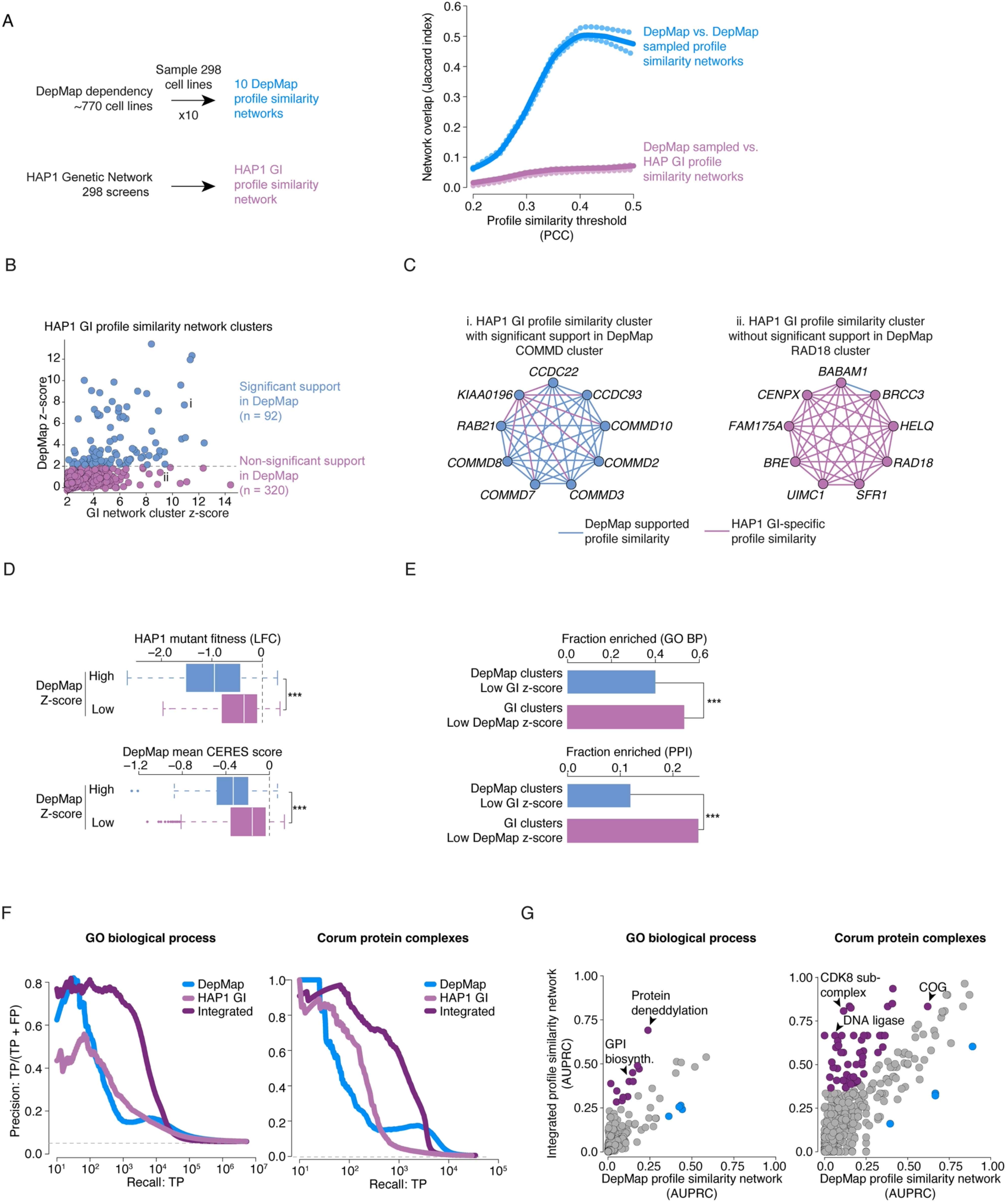
An integrated functional network based on genetic interaction and co-essentiality profiles. **(A)** Comparison of the overlap between correlated gene pairs in the complete HAP1 GI profile similarity and DepMap co-essentiality sampled networks. Co-essentiality networks were constructed by selecting non-overlapping random samples of 298 screens from the DepMap dataset (20Q2). This was repeated to generate 10 co-essentiality networks. Network overlap was assessed by computing Jaccard indices at increasing network similarity thresholds (Pearson’s correlation coefficient thresholds). The same procedure was used to measure similarity of each DepMap co-essentiality network to the HAP1 genetic interaction profile similarity network. Continuous lines represent the mean Jaccard index of the DepMap-DepMap network comparisons (blue) and the DepMap-GI network comparisons (purple). The dotted lines represent the quartiles of Jaccard indices. **(B)** Scatter plot of Z-scores for modules or gene clusters identified from the genetic interaction profile similarity network. Modules with significant similarity in the DepMap co-essentiality network (blue) and modules without significant similarity (purple) are plotted. The grey dashed line indicates Z-score threshold DepMap co-essentiality network similarity. **(C)** Examples of modules derived from the genetic interaction profile similarity network. **(D)** Box plots of mean fitness in HAP1 cells and fitness in DepMap cancer cell lines for genes in significant modules that share similar co-essentiality profiles (blue bars) or do not have strongly correlated co-essentiality profiles (purple bars). **(E)** Bar plot illustrating the fraction of genetic interaction profile similarity network modules (z-score > 2, File S22) enriched for the same GO-BP terms (hypergeometric test, Benjamini-Hochberg-corrected FDR < 0.2) or PPIs (hypergeometric test, Benjamini-Hochberg-corrected FDR < 0.05), including modules uniquely identified in the genetic interaction network profiles (GI module z-score > 2 and DepMap module z-score < 2, purple bars), or genetic network-derived modules that share highly similar DepMap co-essentiality profiles (GI module z-score > 2 and DepMap module z-score > 2, blue bars). **(F)** Precision-recall plots for genes exhibiting similar DepMap co-essentiality profiles (blue), genetic interaction profiles (light purple) or profiles from the integrated network. True positives (TP) involve gene pairs co-annotated to a gold standard set of GO-BP terms (left panel) or gene pairs encoding members of the same CORUM protein complex (right panel). Grey dashed line represents background co-annotation rates. Genes annotated with a mitochondrial-related function were excluded because profile similarity profiles tend to be dominated by mitochondrial genes (*19*). The same precision-recall analysis based on all genes, including mitochondrial genes, is shown in fig. S20F. **(G)** Comparison of individual GO-BPs or CORUM protein complexes captured by the DepMAP co-essentiality network and the integrated network. Nodes above (purple) or below (blue) the diagonal indicate better performance by the integrated or DepMap networks, respectively, based on AUPRC (Area Under a Precision Recall Curve) values, generated per GO-BP process or CORUM complex.

We identified 412 gene clusters or modules in our HAP1 network, each comprising a set of genes that share significantly similar genetic interaction profiles (Fig. 1D, Files S9)(*51*). A fraction of these modules (22%, 92/412) contained genes that were also significantly connected in the DepMap co-essentiality network, such as the COMMD complex involved in endosomal trafficking (Fig. 9B-C, File S22)(*62, 63, 119*). Modules identified in both networks were functionally informative and shared similar characteristics (fig. S19A-E, File S22). However, most HAP1 network modules (∼78%, 320/412) were not supported by strong evidence in the DepMap network and involved genes with more moderate single mutant fitness defects (Fig. 9B-D, File S22). A slightly larger fraction of HAP1 network-specific modules were enriched for functionally related genes and genes associated with PPIs compared to the fraction of enriched DepMap network-specific modules (Fig. 9E). This suggests that the HAP1 genetic network can capture functional relationships among nonessential genes with more subtle fitness defects, perhaps because they can be detected in the normalized context of a single genetic background. In a reciprocal analysis, we examined ∼1,100 functional modules from the DepMap co-essentiality profile similarity network (fig. S19C-E, File S22), most of which were uniquely identified in the DepMap coessentiality network (fig. S19C). DepMap coessentiality network-specific modules involved genes with lower HAP1 expression levels and more modest fitness defects when inactivated in HAP1 cells (fig. S19D).

Because the DepMap co-essentiality and HAP1 genetic profile similarity networks largely captured orthogonal functional relationships, we combined a novel method for processing CRISPR screen data, Onion normalization (*120*), with a deep learning-based approach for network integration, BIONIC (*121*), to generate an integrated network based on the two input networks (*51*). Functionally related gene pairs with roles in the same GO annotated bioprocess or whose products interact physically were connected more frequently in the integrated network compared to either input network (Fig. 9F, fig. S19F). Furthermore, modules derived from the integrated network corresponded to new functions and protein complexes that were not represented in either input network alone. Thus, integrating data derived from our HAP1 genetic interaction network with the DepMap co-essentiality network provides a more comprehensive view of human cell function (Fig. 9G, fig. S19G, File S22).

## Discussion

Applying a genome-scale CRISPR-KO screening approach in HAP1 cells, we tested 222 unique query genes for genetic interactions with ∼17,000 library genes to score ∼90,000 genetic interactions. A network based on genetic interaction profile similarity groups library genes into hierarchically organized modules corresponding to pathways or complexes, biological processes, and subcellular compartments. These modules are connected by coherent sets of negative or positive interactions, mapping a functional wiring diagram of a human cell. Our findings demonstrate that the general principles of genetic interaction networks are highly conserved from yeast to human cells.

A few genes participated in many genetic interactions and represented highly connected network hubs. Physiological and evolutionary properties of genetic network hub genes are also conserved from yeast to human cells, suggesting that gene specific properties can be leveraged to develop general models for predicting genetic network connectivity for genes in different cell types and organisms (*72*).

Yeast essential genes were previously assayed for genetic interactions using hypomorphic (partial function) alleles and shown to participate in ∼5-fold more interactions than nonessential genes. Essential gene interaction profiles are rich in functional information and form the central core of the yeast genetic profile similarity network (*5*). Our ability to score genetic interactions for HAP1 essential genes was primarily limited to disruption of a subset of highly expressed library genes and/or those encoding stable or abundant proteins, which likely decay more slowly over the course of a screen. Nonetheless, several essential genes served as hubs in the HAP1 genetic network. While human essential gene interactions have been previously examined using CRISPRi (*41*), perturbation systems based on transcriptional repression can be technically constrained, effectively repressing only the most highly expressed genes (*122*). Development of query cell lines that stably express hypomorphic alleles should yield informative genetic interaction profiles for human essential genes (*123*).

Genetic interactions highlight the potential for modifying the phenotypic consequences of genetic variation. We characterized ∼3,900 genes whose individual perturbation altered HAP1 cell fitness, including ∼1,500 essential genes and ∼2,400 nonessential fitness genes. We estimate that a comprehensive HAP1 genetic network may encompass ∼1.4 million gene-gene connections, including ∼85,000 synthetic lethal interactions, where a normally nonessential gene becomes essential for viability in a specific genetic background, and ∼45,000 suppressor interactions, where a fitness defect associated with LOF mutant is rescued by mutation of a second gene (*51*). The prevalence of extreme genetic interactions in HAP1 cells is relevant to human disease. Within a population of individuals, almost all phenotypes resemble quantitative traits, with the inherited component driven by the specific genetic background of the individual. Even Mendelian disease genes produce a wide range of phenotypic diversity, which may be traced to genetic modifiers that enhance or suppress the disease mutation. Thus, systematic genetic interaction analysis is critical for precision medicine (*124, 125*). We identified over 1,000 HAP1 extreme synthetic lethal interactions involving more than 350 nonessential disease genes annotated in the OMIM database (fig. S20)(*126*). These synthetic lethal interactions highlight potential disease gene modifiers and also opportunity to assess functional effects of disease genes by variant effect mapping (*127*). Moreover, ∼9% (∼287/3,330) of synthetic lethal interactions involved LOF mutations in known tumor suppressor genes (fig. S20)(*128*) that can be explored for potential targeted cancer therapy (*129–133*).

In rare cases, individuals can harbor mutations that normally cause severe Mendelian diseases but show no reported clinical manifestation of disease. This resilience may be associated with modifiers that suppress the effects of a disease gene (*134*). We found that a LOF mutation in *TAFAZZIN,* a gene linked to Barth syndrome, was suppressed by inactivation of *ABHD18*, which encodes an enzyme that metabolizes cardiolipin to monolysocardiolipin (*99*). Notably, an *ABHD18* chemical inhibitor phenocopied genetic perturbation of *ABHD18* and rescued Barth syndrome phenotypes in patient fibroblasts highlighting the utility of genetic interaction analysis in a model cell line to uncover genetic modifiers and potential targets for disease therapy (*99*). In total, we mapped ∼1,800 suppression interactions in HAP1 cells, including ∼600 suppression interactions involving an OMIM-associated disease gene (fig. S20).

Consistent with previous studies (*101, 102, 135–141*), we observed modest but significant evolutionary conservation of negative interactions. The extent of conservation from yeast to human cells is likely influenced by functionally redundant paralogs, many of which arose from whole genome duplication events (*142, 143*). The human genome contains a larger fraction of duplicated genes (up to ∼60%, ∼10,000/17,000)(*144*) compared to yeast (∼32%, ∼1,900/6,000)(*145*), which influences conservation of the HAP1 genetic network because functionally redundant paralogs tend to only interact with each other, and paralog pairs belonging to larger gene families are even less likely to be connected by negative interactions. Nevertheless, genetic profiles and roles of redundant paralogs can be revealed by mapping more complex trigenic interactions (*80, 81*).

The DepMap project catalogs genetic vulnerabilities in cancer cell lines (*7–11, 53*), and this gene-by-cell line analysis is relevant to the gene-by-gene analysis of our HAP1 network. We found that DepMap CERES score parameters are predictive of connectivity on the HAP1 genetic interaction network, suggesting that these data can be exploited for query gene or cell line selection to facilitate more efficient genetic interaction analysis. Restricting the subset of gene pairs with correlated fitness and expression levels in DepMap data to those that also show a genetic interaction in HAP1 identifies more functionally related gene pairs (fig. S18). For example, gene pairs with a negative ED score and a HAP1 positive genetic interaction can identify extreme suppressor interactions that can modify disease gene phenotypes. Indeed, ∼42% (263/1094) of library genes among the set of gene pairs with a negative ED and positive genetic are annotated to an OMIM disease term. Conversely, gene pairs with a positive ED score and negative HAP1 genetic interaction can identify cancer relevant synthetic lethal/sick gene pairs that can be exploited to develop new therapeutic strategies.

Complex diseases are characterized by detrimental variants in multiple genes, and it is possible that different combinations of variants can lead to the same disease (*146–155*). Indeed, mutations in different genes belonging to the same functional module can lead to the same phenotype (*156, 157*). The highly organized and conserved topology of genetic networks, where coherent sets of negative or positive interactions mediate connections between pairs of different functional module, provides a powerful framework to map the genetic architecture of inherited phenotypes. Functional modules provide prior knowledge that can be used to reduce the statistical burden required to detect genetic networks associated with human diseases (*156, 157*). Our HAP1 genetic network offers an unbiased approach to identify functional modules and the genetic wiring connecting them. Importantly, genetic networks mapped in diverse human cell lines should provide a critical resource for exploring genetic interactions in population-scale biobank datasets, which contain hundreds of thousands of human genome sequences coupled to numerous phenotypes, including diseases (*5, 158–160*).

## Acknowledgments

We thank Jasper Rine for interpreting the yeast *ECM9* genetic suppression interactions. We also thank The Centre for Applied Genomics at SickKids Hospital and the Donnelly Sequencing Centre for assistance with sequencing.

## Author contributions

Conceptualization: BA, CB, MC, JM, CLM; Methodology (Cell line construction and screening): KC, AHYT, JvL, MA, KL, BM, PM, SNM, SA, KA, MA, UB, MCh, RC, KD, A D, AGF, AH, KK, WL, JL, KM, LN, MS, MSR, AS, ES, OS, DTT, DT, JT, SV,KW, ZYW, YXX, SJD, MT, HDMW, MAU, SB, JJC, BN, NM, JW; Methodology (qGI score development): MB, MR, HNW, AZH, KRB, CLM; Investigation (Network analysis): MB, MC, XZ, AZH, MR, KRB, CP, HNW, CR, DF, YL, MU, NB, DC, IH, KL, XL, TR, ES, WW, HLR, GDB, NM, RG; Investigation (Follow-up biology): CB, AFR, NES, GT, JRE, RC, DKK, TU, GWB, BJC, FPR, RL, NGD, SYL, JM; Visualization: MB, MC, AZH, CR, MR, HNW, XZ; Funding acquisition: BA, CB, JM, CLM; Supervision: BA, CB, JM, CLM; Writing-original draft: MC, CB, CLM, JM; Writing-review & editing: All authors.

## Data and material availability

All data are available in the manuscript or supplementary material.

