## Supplemental material for "A global genetic interaction map of a human cell reveals conserved principles of genetic networks"

The PDF file includes:
Materials and Methods
Supplementary text
Figs. S1 to S20

### MATERIALS AND METHODS

#### General information about the datasets

***HAP1 essential gene and single mutant fitness datasets*.** We provide a list of HAP1 essential genes and a catalog of single mutant fitness measurements for all HAP1 genes targeted by the TKOv3 library {Hart, 2017 #4405} in both rich and minimal growth medium, as described below (HAP1 single mutant fitness effects and identification of essential genes). Essential library genes and library gene mutant fitness measurements are provided in Data File S2.

***HAP1 genetic interaction datasets.*** Genetic interaction analyses described in this study are based on either the [1] complete or [2] non-redundant HAP1 genetic interaction dataset as describe below and provided in Data File S4.

*[1] Complete genetic interaction dataset (298 query mutant cell lines x 17, 804 library genes).* The complete dataset comprises 298 genome-wide screens, corresponding to 222 mutant cells lines, each carrying a single loss-of-function (LOF) mutation in a unique query gene of interest. Thus, the complete dataset includes biological replicate screens for a set of 50 query mutant cell lines. The number of replicates/query mutant ranges from 2 to 8 replicates depending on the query mutant cell line. Each query mutant cell line was screened for negative and positive interactions with 17, 804 genes targeted by the TKOv3 library {Hart, 2017 #4405}. This includes a set of 80 library genes that exhibited highly variable single mutant fitness effects across replicate wild-type screens. A list of query mutant cell lines and genome-wide screens, including biological replicate screens performed as part of this study is provided in Data File S1. A list of all library genes targeted by TKOv3, including those with variable single mutant fitness, is provided in Data File S2. The complete genetic interaction dataset was used for all analyses based on genetic interaction profile similarity, unless otherwise indicated. Data File 4 contains single and double mutant fitness ($log_2$ fold change) measurements as well as genetic interactions (qGI scores) derived from the complete dataset, which includes all gene pairs tested in this study, including biological replicates of query mutant cell lines and library genes that showed variable single mutant fitness.

*[2] Non-redundant genetic interaction dataset (222 query mutant cell lines x 17, 724 library genes).* The non-redundant genetic interaction dataset consists of 222 mutant cell lines, each carrying a LOF mutation in a unique query gene. A representative screen was selected for each query gene mutant cell line that was repeated multiple times and replicate screens were excluded from this dataset. Gene pairs involving the set of 80 library genes that exhibited highly variable single mutant fitness in replicate wild type screens were also excluded from this dataset. Thus, the non-redundant dataset comprised negative and positive genetic interactions derived from 222 unique query genes screened against 17,724 library genes. Lists of query mutant cell lines and library genes that comprise the non-redundant dataset are provided in Data File S1 and Data File S2, respectively.

The non-redundant genetic interaction dataset was further filtered based on genetic interaction score (|qGI| > 0.3) and statistical significance (FDR < 0.1) thresholds, which we found to strike an optimal balance between false negative and false positives. The quality estimates of data produced at different thresholds was completed as described below (see section, "Estimating reproducibility of genetic interactions") and is provided in fig. S3 and Data File S5. In total, the non-redundant dataset contains fitness phenotypes associated with perturbation of 3,934,506 unique gene pairs and 88,933 high confidence genetic interactions (47,052 negative and 41,881 positive genetic interactions)(Data File S4). This dataset was used for all analyses based on direct negative and positive genetic interactions, unless otherwise indicated.

***Mitochondrial genes.*** Previous studies showed that gene pairs with strongly correlated co-essentiality profiles, required for fitness of the same set of DepMap cancer cell lines, predominantly involved mitochondrial-related genes, especially those encoding components of the electron transport chain (ETC) or the 55S ribosome {Rahman, 2021 #5666}. The HAP1 genetic interaction profiles for mitochondrial gene pairs were also among the most correlated in the HAP1 genetic network (Fig. 2B, fig. S5B). ETC and 55S ribosome proteins are highly stable and detection of a growth phenotype resulting from the disruption of these genes may not manifest until the wild-type protein is depleted. Indeed, experimental factors, such as sampling time and cell line doubling rate, were shown to impact fitness measurements and amplify correlation between DepMap co-essentiality profiles associated with ETC and 55S ribosome genes {Rahman, 2021 #5666}. Given the potential for various experimental factors to confound scoring and interpretation of mitochondrial-related genetic interactions and profiles, we explored network properties for all tested gene pairs as well as for the subset of gene pairs that excluded genetic interactions involving mitochondrial-related genes. To do so, we compiled a list of 1,102 genes targeted by gRNAs in the TKOv3 library that are also associated with mitochondrial function based on 4 sources: [1] Human MitoCarta 3.0 {Rath, 2021 #6366}, [2] the Kyoto Encyclopedia of Genes and Genomes (KEGG; Oxidative Phosphorylation term){Kanehisa, 2017 #6111}, [3] REACTOME database (Electron Transport pathway term){Gillespie, 2022 #6113}, [4] mitochondrial-related protein complexes annotated in the CORUM protein complex database {Tsitsiridis, 2022 #5807} as well as manually curated genes. The list of genes with mitochondrial-associated function is provided in Data File S2. To exclude mitochondrial signals, the indicated analyses were repeated by excluding gene pairs where both genes appeared on the mitochondrial-associated function gene list. Gene pairs that contained only one gene from the mitochondrial-associated function gene list were not excluded from analyses.

##### TKOv3 library, query mutant cell lines and genome-wide CRISPR screens

***HAP1 genome variant calling.*** To fully characterize the expected editing efficacy of sgRNAs in our TKOv3 library, we generated Illumina whole sequence sequencing for the HAP1 parental cell line to identify variants relative to the reference human genome. The sequencing data is summarized as follows:

| Type | Platform | Depth | # of reads | mean_length | mean_insert_size |
|---|---|---|---|---|---|
| Illumina | HiSeq | 145× | 1453105919 pair-end reads | 150bp | 291bp |

To call variants, we applied the GATK-recommended variant calling workflow {Poplin, 2018 #6377} with the ploidy parameter for HaplotypeCaller set to 1. Discovered variants were then filtered for exonic variants using ANNOVAR {Wang, 2010 #6378}. hg19 was used as the reference genome for all steps. This process resulted in a total of 14,941 exonic variants, which were then analyzed for overlap with TKOv3 guide library target sites (Data File S2). Approximately 0.07% (55 gRNAs) of all gRNAs in the TKOv3 library targeted regions where the HAP1 genome sequence varied from the reference genome (Data File S2). In each of these cases, at least two other gRNAs in the TKOv3 library targeted unaffected regions in the same gene.

***HAP1 query mutant cell line construction.*** Parental human HAP1 cells were obtained from Horizon Genomics (clone C631; sex, male with lost Y chromosome; RRID: CVCL_Y019). Screens were performed in either wild-type HAP1 cells, or HAP1 cells with a defined query gene knockout. HAP1 query mutant cell lines carrying a stable LOF mutation in a query gene of interest were obtained from Horizon Discovery (catalog numbers included in Data File S1) or generated in-house. In-house query mutant cell lines were generated by electroporation of single-guide RNAs (gRNA) targeting the gene of interest complexed with Cas9 protein (RNP complexes) into HAP1 parental cells using the Neon electroporation system (Thermo Scientific). For each query gene, a pool of three distinct gRNAs was used to maximize knockout efficiency (Synthego Gene Knockout Kit v2 guide pools or guides selected from TKOv3). RNP complex preparation and electroporation was carried out according to manufacturer's instructions. Genomic DNA extracted from knockout pools was sequenced by Sanger sequencing using primers proximal to the guide target sites and analyzed using TIDE to estimate editing efficiency. Single clones were selected by limiting dilution and mutations in clonal populations were confirmed using Sanger sequencing of the target region. A complete list of query mutant cell lines used in this study is provided (Data File S1).

***TKOv3 library virus production and multiplicity of infection determination.*** Genome-wide screens were carried out using the Toronto Knockout version 3 (TKOv3) CRISPR library (Addgene #90294), as previously described {Hart, 2017 #4405}. CRISPR library lentivirus production was conducted as previously described {Aregger, 2020 #4690}. To determine viral titers, 3 million HAP1 cells seeded in 15 cm plates were transduced with different dilutions of the TKOv3 lentiviral gRNA library in medium containing 8 µg / mL of polybrene, in a total of 20 mL medium. After 24 h, the virus-containing medium was replaced with 20 mL of fresh medium containing puromycin (1 µg / mL), and cells were incubated for an additional 48 h. Multiplicity of infection (MOI) of the titrated virus was determined 72 h post-infection by comparing percentage survival of puromycin-selected cells with that of cells that were infected but not selected with puromycin (puro-minus controls) after subtracting the percentage survival of uninfected cells in puromycin-containing medium.

***Pooled genome-scale CRISPR screens.*** Genome-wide genetic interaction screens were performed as previously described {Aregger, 2020 #4690}{Varland, 2023 #5760}. Briefly, 3 million HAP1 cells per plate were seeded in 15 cm plates in 20 mL of specified medium. A total of 90 million cells were transduced with lentivirus containing the TKOv3 library at a multiplicity of infection (MOI) of ~0.3 in the presence of polybrene (8ug/mL), such that each gRNA was represented in ~200-300 cells. Transduced cells were allowed to recover for 24 h, selected using puromycin (1ug / mL) for 48 hours, and then a pellet was collected to represent the T0 starting population. The remaining cells were split into triplicate 15 cm plates (3 million cells per plate, 15 million cells total per replicate, representing ≥200-fold library coverage), and cultured for 10-12 population doublings. Three pellets (one from each technical replicate) were collected to represent the endpoint populations. Screens were performed in either DMEM with 10mM glucose and 1mM glutamine ('minimal medium'), or IMDM with 25mM glucose and 1mM pyruvate ('rich medium'), as indicated in Data File S1. All media was supplemented with 10% FBS and 1% penicillin–streptomycin, and cultures were grown at 37˚C with 5% $CO_2$ and passaged every 3-4 days while maintaining a library coverage >200-fold throughout.

***Preparation of sequencing libraries and Illumina sequencing.*** Sequencing library preparation and Illumina sequencing was performed as described previously {Aregger, 2020 #4690}{Varland, 2023 #5760}. Genomic DNA was extracted using the Wizard Genomic DNA Purification Kit (Promega). The gDNA pellets were resuspended in TE buffer, and the concentration was estimated by Qubit using double-stranded DNA (dsDNA) Broad Range Assay reagents (Invitrogen). Sequencing libraries were prepared from 50 µg of the extracted gDNA in two PCR steps, the first to enrich gRNA regions from the genome, and the second to amplify gRNA and attach Illumina TruSeq adapters with i5 and i7 indices, as described previously, using staggered primers aligning in both orientations to the gRNA region {Aregger, 2020 #4690}{Varland, 2023 #5760}. Barcoded libraries were gel purified, and final concentrations were estimated by Qubit or qRT–PCR. Sequencing libraries were sequenced on an

Illumina HiSeq2500 using single-read sequencing and were completed with standard primers for dual indexing with HiSeq SBS Kit v4 reagents. The first 21 cycles of sequencing were dark cycles, or base additions without imaging. The actual 36-base read begins after the dark cycles and contains 2 index reads, in which i7 is read first, followed by the i5 sequences. The T0 and end time point samples were sequenced at 400- and 200-fold library coverage, respectively.

### HAP1 single mutant fitness effects and essential genes

We estimated robust single mutant fitness effects for all human genes targeted by our TKOv3 gRNA library in HAP1 cells. We also defined a robust set of HAP1essential genes.

***Library gene mutant fitness catalogue.*** Single mutant fitness (SMF) effects for 17,804 protein-coding genes targeted by the TKOv3 gRNA library were estimated from a collection of 39 independent wild-type control HAP1 screens. Those screens were completed in minimal (n = 21) and rich (n = 18) cell culture medium, and library gene SMF in each medium was estimated separately. First, SMF effects were estimated from each individual wild-type screen by subtracting the sequencing depth-normalized and $\log_2$-transformed read-counts representing the abundance of each gRNA in the library at the start of the experiment (T0) from those at the experimental endpoint. For each wild-type control screen, gRNA-level $\log_2$-fold changes (LFC) were then adjusted such that the median of all gRNA targeting a gold-standard, non-essential gene set {Hart, 2014 #4147}{Hart, 2016 #4406} is equal to zero, and mean-summarized across the three technical replicates per screen. Across all wild-type control screens in each media condition, gRNA LFC were then mean-summarized for all gRNAs not flagged by our empirical quality score (see Filtering TKOv3 gRNAs). Finally, library gene SMF was derived by mean-summarizing across the 2-4 independent gRNAs targeting each library gene. Library gene SMF measure in each media condition is provided in Data File S2 and shown in fig. S1.

***Random forest model to identify HAP1 essential genes***. In addition to library gene SMF, we also defined the subset of library genes that are essential for HAP1 growth in cell culture. The identification of the HAP1 essential gene set was formulated as a binary prediction problem. Features for the predictive model were derived from the complete collection of HAP1 wild-type control (described above) and query mutant genetic interactions screens (described below), and a gold standard set of “core” essential genes that are essential in at least 60% of human cell types examined in the 20Q2 release of Cancer Dependency Map project dataset (DepMap; depmap.org/portal){Meyers, 2017 #4545;Tsherniak, 2017 #4408;Dempster, 2021 #5782}. A gene was labeled as core essential if the gene met a CERES score cutoff of < -1 in more than 60% of DepMap cancer cell lines. A CERES score < -1 threshold corresponds to the median score of all essential genes in the DepMap data. We note that this is relatively stringent criteria as the goal was to identify a set of genes that is highly likely to be essential across most human cell lines as a basis for training our model. This set of high-confidence core essentials resulted in 733 essential genes amongst the 17,804 genes targeted by our library (Data File S2). We reasoned that these core essential genes could be used to learn specific characteristics of essential genes in our HAP1 screen data. The assumption is that the majority of “core” essential genes are also essential in HAP1, but that training a predictive model to recognize the features of those genes based on HAP1 data would identify additional HAP1 essential genes with high accuracy. This provides a more precise definition of HAP1 essential genes by leveraging the wealth of phenotype data collected as part of this study compared to more simple strategies such as applying a single threshold on the average LFC per gene derived from wild type control screens.

We trained a random forest supervised model to predict HAP1 essential genes. Three different types of features from our HAP1 screen data were used as input to this model, which can be grouped into three general categories: [1] *Central tendency (mean/median LFC) features*. These reflect average single mutant phenotypes across the 298 genome-wide genetic interaction screens completed as part of this study: [2] *Variance features*. These reflect variability of individual genes' phenotypes across the 298 screens. Specifically, the standard deviation and coefficient of variance of LFC values were used as inputs to our model. [3] *Screen dynamics features*. These features were derived from time-resolved wild-type control screens where we sampled gRNA abundance at different timepoints over the course of the screen, which helped to differentiate essential genes from genes that simply result in a slow growth phenotype in cell culture. Screen dynamics metrics were based on the pairwise slopes between gRNA abundance measured at different time points. The complete set of features, detailed descriptions, and the feature table used for this essential gene model are provided as Data File S3.

Input features along with the gold-standard core essential gene set was used to build and optimize parameters for a random forest classifier using a 5-fold cross-validation approach. For each gene, the predicted probability of gene essentiality was extracted from the model for which it was in the held-out fold, and a precision-recall curve was used to evaluate overall performance of the model. All genes that met a score threshold corresponding to a 0.95 recall or higher on the gold-standard essential gene set were defined as "HAP1 essential". This process was conducted separately for HAP1 screens performed in minimal and rich media to predict two essential gene sets, one in each media condition. The overlapping essential genes from the rich and minimal media conditions was used to define the complete HAP1-specific essential gene set, which consists of 1,524 genes. Genes that were not deemed essential by this process but exhibited a statistically significant SMF effect (LFC < 0 or LFC > 0) were deemed "non-essential fitness effect" genes, which included an additional 2,417 genes. The complete set of HAP1 essential library genes and nonessential genes associated with a fitness effect are listed in Data File S2 and shown in figs. S1-S2.

***Features associated with HAP1 essential genes.***

We analyzed the collection of HAP1 essential genes, described above, for differences in several physiological, evolutionary and functional gene features listed in Data File S14. For continuous-valued and discrete-valued features, we used Wilcoxon rank-sum tests to assess the difference in the HAP1 essential gene set distribution from the distribution of the non-essential gene set. For features that were statistically significant, we plotted the log-ratio of the mean of the essential gene group to the non-essential gene group for that feature (absolute value was first applied to negative means, fig. S2E). The essential gene set was also tested for enrichment across the Reactome pathways and GO biological process terms Data File S3). The results of these analyses are shown in fig. S2

### A quantitative genetic interaction (qGI) score

To derive quantitative genetic interactions, we modeled expected combinatorial gene perturbation effects and estimated the deviations from this null model as the quantitative genetic interaction (qGI) score. Each step is described in more detail below, but we provide a brief overview of the process here. We compare the fitness effects of each query mutation screen with a wild-type HAP1 control screen and fit a loess (locally estimated scatterplot smoothing) regression as the null model. This partially adjusts the growth effect distribution of all library genes to the background of a given query. Additional corrections to the data improve this null model. Since genetic interactions are estimated semi-independently for each query screen, those corrections are applied on a per genetic interaction screen basis followed by a network-wide set of corrections for systematic biases that cannot be estimated from individual screens. Each of these steps is described briefly below and in detail in an associated manuscript {Billmann, 2025 #6383}. All corrections were completed at the gRNA level to increase sensitivity unless stated otherwise.

***Filtering TKOv3 gRNAs.*** Based on screens completed as part of this study, we identified a small set of guides with clear efficacy issues and removed them from further analysis. Specifically, for every gene targeted in the TKOv3 library, we compared gRNA-level genetic interaction profiles (see qGI score described below) across all screens. For each within-gene gRNA pair, we computed a pairwise Pearson correlation coefficient and median-summarized this for each single gRNA (excluding self-correlation). This score quantified how similar genetic interaction profiles of gRNAs were to those of other gRNA targeting the same gene. Based on this metric, we removed the worst gRNA for 517 of 17,804 genes (2.9%) prior to computing the gene-level genetic interaction scores (qGI scores), the associated FDR, and the single mutant fitness (SMF). More details on the gRNA quality score are provided in {Billmann, 2025 #6383}.

***Individual screen corrections:***

*Hybrid T0 normalization.* First, we measured gRNA abundance by sequencing the library for each screen after the puromycin selection (T0) and balanced this abundance with an average of the T0 gRNA abundances derived from all other screens in the dataset. This prevented gRNA loss during puromycin selection, which can lead to spurious genetic interaction calls.

*MA transformation.* Next, for each wildtype control-query screen pair, we MA-transformed the fitness scores prior to fitting the loess model. This stabilized null model estimates for extreme positive or negative fitness scores.

*Effect size scaling.* Query mutant cell lines with extreme single mutant fitness phenotypes often exhibited strong, non-specific deviations from the null model. These effects were controlled by adjusting residual values for a screen by matching the variance of the central 80% of residual effects.

*Multiple WT-query pairs.* The null model is fit for all pairs of a query screen with all wildtype control screens performed in the same growth medium (21 minimal media screens; 18 rich media screens) and the residual effects are shrunk by providing the highest weight to the pair with the smallest sum of squared residuals.

*Local genetic interaction shift correction.* Finally, residual scores for gRNAs are arranged according to the genomic positions of their target genes, and smoothing is conducted to identify genome regions where a collection of gRNAs that target adjacent genomic regions deviate from the null model either positively or negatively. The median of residual scores of detected shifts was set to 0.

***Network-wide corrections:***

*Variable single mutant fitness correction.* To specifically isolate screen-to-screen variation due to the query mutation, we computed mock residual fitness scores by comparing wildtype control screens in each media condition to other wild-type control screens conducted in the same medium.

Mock residual fitness scores represent "pseudo-genetic interactions", or genetic interactions that are identified in the absence of a query mutation, and thus provide an estimate of expected single mutant fitness variability per library gene in actual query mutant cell line screens. Based on these estimates, we first corrected the per-gRNA variance across query screens to account for the observed variation across wildtype control screens. We also observed reproducible co-variance amongst gRNAs (and the corresponding genes) across multiple control screens, suggesting there are biological factors driving systematic effects in control screens (and query screens). To remove this covariation, we used Singular Value Decomposition (SVD) to isolate singular vectors describing the variation observed across control screens then projected the query screen data onto these control screen components and subtracted the resulting projections from the query residual matrix. This effectively removed signal from the query screen data that reflects patterns also observed in control screens. This analysis also identified library genes with highly variable single mutant fitness phenotypes in the control screens. A set of 80 library genes that exhibited the most variable fitness effects ("core" set of highly variable fitness genes) were excluded from all analyses related to genetic interaction degree and density in this study (Data File S2). An "expanded" set of library genes that exhibited more subtle variation in the control screens was also identified, which included 426 additional genes (Data File S2). The complete set of 507 "core" and "expanded" library genes with variable single mutant fitness were excluded from the construction and analysis of the HAP1 genetic interaction profile similarity network (see "Constructing a genetic interaction profile similarity network" section below) to ensure library genes susceptible to variation in controls did not affect our conclusions from these analyses. Information related to all query and library genes is provided in Data Files S1 and S2, respectively.

*Global systematic effect correction.* Minimal and rich medium SVD-corrected "pseudo-genetic interaction" matrices were merged into a global query gene-by-library gene residual matrix. Patterns explaining substantial variation in this matrix are not expected to reflect true genetic interactions (given that GIs are expected to be sparse), and thus, we applied a second SVD normalization to the merged matrix to removed contributions of the first four singular vectors.

*gRNA quality control and gene-level genetic interaction scores.* As described above (see Filtering TKOv3 gRNAs), based on an empirical gRNA quality score, we removed the lowest scoring gRNA for a total of 517 genes. Additional details on all of these scoring and normalization steps and evaluation of their specific impact on the resulting quantitative genetic interaction scores are available elsewhere {Billmann, 2025 #6383}. These correction steps produced corrected residual (gRNA-level genetic interaction) scores derived from analysis 324 query screens that comprised 69,474 high-quality gRNAs targeting 17,804 protein-encoding genes. All genes were targeted by at least two gRNAs each, and 90.4% of genes were targeted by four independent gRNAs.

To measure quantitative GI (qGI) scores for query-library gene pairs, the mean of the 2-4 residual values for a given query-library gene combination was computed. Each qGI score was assigned a statistical significance measure using the following procedure. gRNA-level residuals for each query-library gene pair were computed (i) prior to network-wide corrections to preserve experimental uncertainty of the measurements and (ii) resolved to show all contrasts between a given query and the multiple (min n = 21; rich n = 18) wildtype control screens. By considering those 18 or 21 gRNA-level values as technical replicates and the 2-4 independent gRNA sequences as biological replicates for the same gene pair, we computed *P*-values reflecting the deviation from 0 using a moderated t-test followed by multiple hypothesis correction using the Benjamini-Hochberg method to control FDR. As detailed below (see section Estimating reproducibility of genetic interactions), we performed extensive analysis of the reproducibility of genetic interactions at a range of effect size and FDR thresholds (Data File S5). Genetic interactions at a standard confidence threshold were identified using |qGI| > 0.3 and FDR < 10% thresholds, and strict genetic interactions were identified using |qGI| > 0.6 and FDR < 1% thresholds. Sensitivity and precision of the interactions identified at those thresholds are characterized in the "Estimating reproducibility of genetic interactions" section below.

#### Estimating reproducibility of genetic interactions

We applied a Markov Chain Monte Carlo (MCMC)-based approach to estimate the reproducibility of genetic interactions, as previously described {Costanzo, 2021 #6273}. Briefly, we performed 4-5 biological replicate screens for 7 representative query mutant cell lines (*FANCG, NGLY1, PDCD5, PELO, PTAR1* and *VPS52*)(see Data File S5). First, genetic interactions were scored independently for each biological replicate screen, generating qGI scores and an associated FDR for each library gene targeted in every query gene biological replicate screen. A "consensus profile" was then inferred for each query mutant starting from a thresholded version of the replicate interaction profiles, where a weak confidence threshold is applied (FDR < 50%) to define a set of positive or negative interactions for that screen. An MCMC approach is then used to jointly infer false negative rates (FNR) and false positive rates (FPR), as well as a binary consensus GI profile (separately for positive/negative GI). After this inference process converges, the resulting consensus profile is used to generate a data-derived interaction standard for evaluation of individual screen data (assuming pairs with posterior probability of interaction of > 0.5 are interactions). Finally, this consensus standard is used to evaluate per screen performance characteristics over a range of different qGI score effect size and statistical significance cutoffs (FDRs) on the per screen scores. At each evaluated qGI score effect size and statistical significance cutoff, we measured precision and recall metrics, which were then summarized across this collection of 7 replicated query screens. This evaluation led to the selection of two recommended cutoffs for our dataset. A "strict" cutoff (|qGI score| > 0.6 and FDR < 1%), which results in an estimated precision of per screen called genetic interactions of 0.91 and a recall of 0.12 and a "standard" cutoff (|qGI score| > 0.3 and FDR < 10%), which results in an estimated precision of per-screen called interactions of 0.61 and a recall of 0.37. For all analysis that depended on a thresholded set of genetic interactions, we applied the "standard" cutoff, unless otherwise noted. Data File S4 provides unfiltered qGI scores and FDR statistics for all tested gene pairs. Precision and recall estimates are provided over a range of different qGI score and FDR thresholds highlighting the relative balance between false positives and false negatives such that a user can define thresholds appropriate for future studies (Data File S5). The results of this analysis are presented in fig. S3A and in Data File S5.

#### Validating genetic interactions with an independent gRNA library

We constructed an independent gRNA library consisting of ~37,000 gRNAs targeting ~1,200 genes that showed significant negative or positive genetic interactions in at least one of five genetic interaction screens using *ARID1A*, *FANCG*, *FASN*, *NGLY1* or *PTAR1* query mutant cell lines (fig. S3F, Data File S6). Candidate gRNA sequences were identified from FASTA files of the hg38 chromosomes using the regular expression pattern '[ACGT]{20}.GG', then filtered to remove guides with polyT stretches and Esp3I restriction sites. Next, genomic coordinates for the filtered guide sequences were overlapped with exons from the desired target ~1,200 genes using bedTools 'intersectBed' to produce a list of candidate sgRNAs. Finally, to produce a library that widely covered the target genes, up to 40 sgRNAs per gene were selected randomly from the filtered candidate gRNAs. Of the genes targeted, 743 were targeted by 40 sgRNAs, while the remainder of genes were targeted by fewer. The vast majority (> 95%) of gRNAs in this library were not represented in the TKOv3 library. To calculate differential LFC scores, we performed the following steps: [1] Remove gRNAs with either NA or read count < 30 in any of the WT or query T0 screens; [2] Depth normalize read count data as follows: normalized reads per screen = [(raw reads/sum of reads) * 1,000,000] + 1; [3] Calculate LFC between each replicate and the reference timepoint; [4] Average across all gRNAs and technical replicates per library gene to obtain the final differential LFC for each query; [5] Calculate the differential LFC for each gene based on its LFC values measured in the query gene mutant cell line and the reference WT control cell line; [6] For each query gene, separate the 1,215 library genes into groups of "negative", "positive", and "no interaction" based on the qGI score and FDR associated with the corresponding gene pair derived from genome-wide TKOv3 screens. Raw read counts and differential LFC scores used to independently identify genetic interactions for 5 different query genes of the gRNA tiling library, are provided in Data File S6 and results from this analysis are shown in fig. S3F.

#### Hierarchical clustering of HAP1 genetic interaction profiles

We modified a previously described hierarchical clustering algorithm to cluster the complete HAP1 genetic interaction dataset and produce a set of uniformly sized gene clusters across layers of the dendrogram {Xiong, 2024 #6365}(https://doi.org/10.5281/zenodo.15320010). Briefly, for each dendrogram layer, an iterative process identified gene clusters based on their size and a signal-to-noise ratio (SNR), calculated as the average within-cluster Pearson correlation divided by its standard deviation {Xiong, 2024 #6365}. Gene clusters that met a minimum size requirement (at least three genes) and ranked in the top percentile of SNR were selected and removed from subsequent clustering iterations. Genes that did not belong to a cluster in the first iteration were re-clustered in subsequent iterations until only one or no genes remained or until remaining clusters did not meet the selection criteria. This process was then repeated at the next layer of the dendrogram. This iterative approach within each layer ensured balanced clustering by preventing a few highly coherent clusters from dominating the hierarchy. We applied this clustering algorithm to the complete genetic interaction dataset (298 query cell lines x 17, 804 library genes) to generate two layers of “Global” gene clusters (L2 parent clusters and L1 child clusters), which are provided in Data File S9. Results of this analysis are also shown in Fig. 1D and fig. S7A.

We also applied the algorithm to cluster a smaller subset of 3784 genes that comprise the high-confidence profile similarity network and used this subset of clusters to define the HAP1 functional hierarchy, as described below (see section: A genetic interaction profile similarity-derived functional hierarchy, Data File S16).

#### Constructing a genetic interaction profile similarity network

To construct the genetic interaction profile similarity network shown in Fig. 2, we performed two additional filtering/normalization steps before constructing the profile similarity network. The purpose of both of these additional steps was to ensure that the similarity network analysis was focused on the highest confidence set of genes and that any non-specific interaction signal was removed before computing similarity networks. First, excluded an expanded set of 507 library genes associated with variable single mutant fitness phenotypes from the complete HAP1 genetic interaction dataset. The list of library genes with variable mutant fitness is provided in Data File S2. Second, additional normalization procedures were applied to genetic interaction profiles in order to construct the profile similarity network. These normalization procedures are described in detail below.

***Genetic interaction profile normalization*****.** We identified two gene expression patterns related to several query mutant cell lines that were distinct from the HAP1 parental cell line gene expression profile based on genome-wide gene expression profiling of 60 selected HAP1 query mutant query cell lines, described below (see section Expression profiling of HAP1 query mutant cell lines). In both cases, the group of cell lines that exhibited a similar differential expression signature also exhibited similar patterns of genetic interactions (fig. S5C). For most cell lines, the observed differential expression patterns had no clear connection to the query mutation. Thus, we concluded that the observed expression signatures and genetic interaction profiles shared by query mutant cell lines comprising each group may reflect experimental and/or biological factors that were not entirely related to the specific query gene mutation. As a result, we applied additional normalization procedures to the genetic interaction dataset to minimize "non-specific" similarities between genetic interaction profiles prior to constructing the HAP1 genetic interaction profile similarity network.

First, we applied standard two-dimensional clustering analysis {Eisen, 1998 #1353} to the gene expression and the HAP1 genetic interaction data corresponding to the subset of 60 query mutant cell lines to identify two distinct groups of query genes that shared similar gene expression signatures and genetic interaction profiles (fig. S5C). The first group consisted of 5 query mutant screens that shared a similar gene expression and genetic interaction profiles (GSK3A_GIN247, GSK3B_GIN237, FANCA_GIN010, BCL2_GIN338, EMC6_GIN231)(Data File S1). The second group involved 5 query gene screens (CRIPT_GIN260, MPV17_GIN230, C5orf34_GIN261, TMEM126A_GIN309 and POLR2A_GIN281)(Data File S1) that also shared similar gene expression and genetic interaction profiles but whose profiles were distinct from the profiles associated query genes in the first group. For each group, we defined an "expression centroid" and a "qGI centroid" signature representing the average differential expression pattern and average genetic interaction pattern (based on qGI scores) for each group of query genes, respectively. We next examined the complete set of 298 genome-wide screens to identify additional query genes with genetic interaction profiles that resembled the genetic interaction profiles of query genes belonging to Centroid 1 or Centroid 2. An additional 4 query gene screens (GSK3A_GIN405, PC_GIN219, SLX1_GIN239, KAT7_GIN173) exhibited Centroid 1-related genetic interaction profiles and another 3 screens (POLR2A_GIN395, C5orf34_GIN357, TMEM126A_402) were associated with Centroid-2 related genetic interaction profiles (Data File S1). We then computed final qGI centroid vectors based these expanded query gene screen groups by computing the average qGI signature across the screens in each group, resulting in a 1x17,804 qGI vector for each centroid group. The complete genetic interaction dataset was first normalized for the Centroid 1 vector and then subsequently normalized for the Centroid 2 vector, using the following procedure:

[1] Each centroid was unit normalized by dividing by its $l^2$ norm:

$$U_{r\times 1} = C_{r\times 1}/\parallel C \parallel_2$$

[2] Each centroid unit vector was projected onto the complete qGI score matrix of 298 query screens:

$$M_{r\times c}^{projection} = U_{r\times 1} \times U_{r\times 1}^{T} \times M_{r\times c}$$

[3] The resulting centroid projection was subtracted as follows, creating a corrected matrix of qGI scores:

$$M_{r\times c}^{corrected} = M_{r\times c} - M_{r\times c}^{projection}$$

The resultant centroid-corrected genetic interaction dataset excludes 507 library genes with variable single mutant fitness (listed in Data File S2) is provided as a clustered matrix in Data File S8 and shown in fig. S6. The same dataset was used to construct the profile similarity network shown in Fig. 2 and described below.

***Expression profiling of HAP1 query mutant cell lines.*** Gene expression analysis was performed on 61 HAP1 cell lines (wild-type, and knockout queries) using RNA sequencing (RNAseq). Query cells were grown in the same conditions used for the screens, and pellets containing $2x10^7$ cells were collected when the cells reached 70-80% confluence. Total RNA was extracted from the pellets using Qiagen RNeasy kits. Poly(A)-enriched mRNA libraries were prepared using NEBNext Ultra II Directional RNA library prep kit using the manufacturer's protocol. Library quality was assessed on an Agilent 2100 Bioanalyzer, and all samples had an RNA integrity value (RIN) of 9.8 or higher. Samples were sequenced on an Illumina NovaSeq 6000 with an S1 flowcell with paired-end 151bp reads to an approximate depth of 25 million reads per sample. After checking the sequencing quality with FastQC (v.0.11.9), reads were aligned to human genome build hg38 with Gencode v32 gene annotations using the STAR short-read aligner (v.2.7.9a) using the following parameters: --outSAMtype BAM SortedByCoordinate --quantMode GeneCounts --sjdbGTFfile gencode.v25.annotation.gtf. Per-sample read count matrices were merged with ENSEMBL and Entrez Gene annotations in R. With the bulk RNA sequencing data covering 60 query screens across 16,546 TKOv3 library genes, we calculated their log2-fold-change (LFC) values against the single mutant (WT) expression data as differential expression (DE). Raw FASTQ files and read counts are available through the GEO database with accession GSE296341.

***Mapping the genetic interaction profile similarity network.*** We constructed a genetic interaction profile similarity network measuring similarity between library genes targeted in our genome-wide screens. First, Pearson correlation coefficients (PCC) were measured between all pairs of library genes using each gene's profile across 298 queries in the Centroid-normalized qGI score matrix described above (Data File S8). We then removed edges between gene pairs whose profile similarity did not satisfy a PCC > 0.41 threshold. Finally, we removed weak similarity edges for the most highly connected genes thus setting a maximum of 30 edges/connection per gene/node in the profile similarity network. This resulted in a "core" genetic interaction profile similarity network consisting of 1017 genes and 3483 edges. The core network was visualized using the "yFiles Organic" network layout in Cytoscape (Data File S10){Smoot, 2011 #1376}. A subset of 234 genes were disconnected from the core network, and these genes were manually placed on the periphery of core network closest to the region of the network enriched for functions related to the specific disconnected gene set.

Gene coordinates in the core network were then fixed, and the network was expanded to include additional genes using a K-nearest neighbors (KNN) approach. For each gene in the core network, we ranked all other genes based on their genetic interaction profile similarity (PCC) derived from the complete, centroid-corrected genetic interaction profile similarity matrix and then selected the top K neighbor genes, where K was set to 35, and all connected genes satisfied a minimum correlation threshold (PCC > 0.1). Each node in the core network could add a maximum of K new auxiliary nodes meeting this threshold. To prevent network oversaturation caused by highly connected nodes, only one edge connecting a new gene with the highest profile similarity to another gene in the core network was added to the expanded profile similarity network. This resulted in an expanded genetic interaction profile similarity network consists of 3,784 genes and 6,433 unique edges (Fig. 2, Data File S10). The core and expanded profile similarity networks are also provided as Data Files S8 and S10. The source code and files for this network construction process are also available from (https://doi.org/10.5281/zenodo.15320010).

***Biological process annotation of the genetic interaction profile similarity network.*** The expanded genetic profile similarity network was annotated using a modified version of Spatial Analysis of Functional Enrichment (SAFE){Baryshnikova, 2016 #4682}. The modified SAFE code is available from (Https://doi.org/10.5281/zenodo.15320010). To functionally annotate the genetic interaction profile similarity network shown in Fig. 2B, we applied the modified SAFE method using default parameters with the following exceptions:

node_distance_metric='shortpath_weighted_layout'
neighborhood_radius=0.12
attribute_distance_threshold=0.65
enrichment_threshold=0.2
multiple_testing=TRUE

To calculate node distance and define spatial network neighborhoods, we weighted profile similarity edges based on the ratio between the Euclidean distance ($d_{ij}$) measured between two genes/nodes in the network and the profile similarity (Pearson Correlation Coefficient, PCC) between the corresponding connected genes (i.e. edge weight = $d_{ij}$ / PCC). SAFE was combined with a reduced set of Gene Ontology (GO) Biological Process (BP) terms filtered for size (GO BP terms containing between 1-500 genes) to identify functionally enriched regions of the genetic interaction profile similarity network. SAFE analysis identified 42 functionally enriched network domains, which were reviewed and manually merged into 17 larger functional domains (Fig. 2B, Data File S11).

***Subcellular compartment annotation of the genetic interaction profile similarity network.*** To functionally annotate the genetic interaction profile similarity network shown in Fig. 2C, we applied the modified SAFE method using default parameters with the following exceptions:

node_distance_metric='shortpath_weighted_layout'
neighborhood_radius=0.15
enrichment_threshold=0.001
multiple_testing=TRUE

We used SAFE to identify regions of the profile similarity network enriched for genes that localize to the same subcellular compartment as defined by curated cell compartment annotations {Binder, 2014 #6288}

***Protein complex annotation of the genetic interaction profile similarity network***. We projected centroids of manually curated CORUM complexes {Tsitsiridis, 2022 #5807} based on the spatial positions of their member genes. Complexes were retained if at least two of their genes were present in the network, and each gene was assigned to only one complex, favoring the largest in cases of overlap. These criteria yielded a nonredundant set of 71 complexes. Centroids were computed from the Euclidean coordinates of genes in the embedded network layout, with minor jitter added to prevent visual overlap. Complexes were visualized and colored according to biological process as shown in Fig. 2D. A list of complexes visualized on the profile similarity network is provided (Data File S11).

### Using the HAP1 interaction profile similarity network to annotate function

To demonstrate the utility of the HAP1 genetic interaction profile similarity to functionally annotate genes and datasets, we applied SAFE or standard enrichment analysis to identify regions of the genetic interaction profile similarity network that were specifically enriched for genes sets derived from the datasets described below.

***Annotating individual query gene genetic interaction profiles.*** For a given query gene, we applied the standard genetic interaction confidence threshold ($|qGI| > 0.3$, $FDR < 0.1$) to generate a binarized profile of negative and positive genetic interactions for a given query gene. We then used the modified version of SAFE (Https://doi.org/10.5281/zenodo.15320010), described above, to identify regions of the HAP1 genetic interaction profile similarity network that were enriched for genes that showed negative or positive interactions with the query gene of interest. The modified SAFE method was applied using default settings with the following exceptions:

node_distance_metric='shortpath_weighted_layout'
neighborhood_radius=0.05
enrichment_threshold=0.001
multiple_testing=TRUE

This method was applied to identify regions of the genetic interaction profile similarity network that were enriched for negative and positive interactions identified with HAP1 query mutant cell lines carrying LOF mutations in either *C1orf112* as shown in fig. S7B or *VPS52* as shown in Fig. 5E.

***Annotating chemical-genetic interaction profiles.*** A chemical-genetic interaction refers to a mutant strain that exhibits sensitivity or resistance to a particular bioactive compound and the set of mutants that show differential compound sensitivity generates a chemical-genetic interaction profile reflecting the compound mode-of-action {Piotrowski, 2017 #4436}. We performed genome-wide chemical-genetic interaction screens to identify genes that showed differential sensitivity to different bioactive compounds (Data File S13). Chemical-genetic interaction screens were performed as previously described {Masud, 2022 #5659}. Briefly, HAP1 cells were grown in suspension format in 100-150mL IMDM medium supplemented with 1:700 dilution of anticlumping reagent (ThermoFisher, Cat# 0010057DG) in Erlenmeyer flasks and maintained in a shaking incubator (150 rpm, 37°C, 5% $CO_2$). At each timepoint, cells were collected by centrifugation and trypsinized with TrypLE.

One hundred million wildtype HAP1 cells stably expressing Cas9, were transduced with the lentiviral TKOv3 library at a low MOI (~0.3), such that each guide is represented in 200-300 cells. 24 hours post infection, cells were selected for viral integration with 1μg/mL puromycin for 48 hours.

Cells were then harvested and pooled and 30-50 million cells were collected for subsequent gDNA extraction and calculation of library representation at the initial timepoint, T0. At this stage, the pooled cells were divided into three technical replicates, each consisting of 15 million cells to maintain 200-fold library coverage. Selection screens were performed by subculturing cells after four days (T4), at which point each technical replicate was divided into treatment arms, including an untreated one (DMSO, vehicle control), and sub-cultured every three to four days for up to 15 days (~15 doublings). Compounds were added at predetermined concentrations with ABD957 used at 10uM, resulting in LD30 and NGI-1 was used at 9.5-15uM with an average LD65.

Genomic DNA from pelleted cells at the initial timepoint, T0 and the endpoint, T14 were isolated using the QIAamp Blood Maxi Kit (Qiagen) and DNA concentrations were determined using the Qubit dsDNA Broad Range Assay kit (Invitrogen). To prepare sequencing libraries, 50µg of genomic DNA was processed using a two-step nested PCR approach, to amplify the sgRNA and append Illumina TruSeq adapters with i5 and i7 indices. The barcoded libraries were then purified with the QIAquick PCR Purification and Gel Extraction kits (both from Qiagen). The resulting samples were then sequenced on an Illumina HiSeq2500, using single-read sequencing. Quantitative chemical-genetic interactions were measured used a modified version of the qGI scoring method described above and elsewhere {Lin, 2024 #6328}.

We applied a standard genetic interaction confidence threshold (|qGI| > 0.3, FDR < 0.1) to generate a binarized profile of negative and positive chemical-genetic interactions associated with the bioactive compound, NGI-1. We then used the modified version of SAFE (Https://doi.org/10.5281/zenodo.15320010), described above, to identify regions of the HAP1 genetic interaction profile similarity network that were enriched for genes that showed negative or positive chemical-genetic interactions with the NGI-1, as shown in fig. S7D and provided in Data File S13. The modified SAFE method was applied using the parameters described above (see section Annotating individual query gene genetic interaction profiles).

***Annotating OMIM disease genes*.** The OMIM disease set was downloaded from https://omim.org/ in July 2021 {Hamosh, 2005 #5684} and filtered to comprise a set of 208 diseases with at least 4 annotated genes. A hypergeometric test was conducted to assess which of the 17 biological process-enriched domains on the genetic interaction profile similarity network (Fig. 2B) were enriched for a particular disease gene set, assuming the background gene set of all genes present on the profile similarity network. Results pertaining to this analysis are shown in fig. S7C.

***Annotating GWAS-phenotype associated genes.*** The genome-wide association study (GWAS) trait set was downloaded from https://www.ebi.ac.uk/gwas/ {Sollis, 2023 #6368} in July 2021. This data was filtered to include a subset of 194 traits with at least 60 associated variants and linked genes that passed a GWAS significance threshold of $P < 5x10^{-7}$. A hypergeometric test was conducted to assess which (if any) of the 17 biological process-enriched domains on the genetic interaction profile similarity network (Fig. 2B) were enriched for genes associated with a particular trait, assuming the background gene set of all genes present on the profile similarity network. Results pertaining to this analysis are shown in fig. S7C.

### Genetic interaction degree and network density analysis

***Genetic interaction network hub genes.*** We defined library genes with a positive genetic interaction degree equal to or higher than the 95th percentile as “positive genetic interaction hub” genes. Similarly, genes were defined as “negative genetic interaction hub” genes if their negative GI degree was equal to or greater than the 95th percentile. Positive and negative GI degrees associated with all library genes are provided in Data File S14. Reactome pathway enrichment analysis was performed on the hub gene sets against the universe of all library genes, restricting the set of pathways tested to those with more than 10 total genes amongst the library gene universe and less than 300 member genes. The *P*-values were adjusted with the Benjamini-Hochberg correction and were filtered using an FDR < 0.2 threshold. The enrichment results are provided in Data File S14. Analysis of genetic interaction network hub genes are shown in Fig. 3 and fig. S8.

***Correlation analysis of genetic interaction degree.*** We assessed the correlation of the number of negative and positive genetic interactions associated with each library gene derived from the nonredundant dataset with several different physiological, evolutionary and functional gene features. Gene features are described in Data File S14 and include binary-, continuous-, and discrete-valued features. We used Wilcoxon rank-sum tests to determine if binary gene features partitioned genes into two groups such that one group had an average degree that was significantly higher or lower than the other group. Genes for which the value of the binary feature was unknown were excluded from the feature's test. For continuous- and discrete-valued gene features, we calculated the Pearson correlation coefficient (PCC) between genetic interaction degrees of genes and each feature. We considered the degree of genes targeted by the TKOv3 library, by counting the number of interacting query genes, meeting the standard confidence threshold (|qGI| > 0.3, FDR < 0.1) derived from the nonredundant dataset. The correlations for library genes are provided in Data Files S14 and shown in Fig. 3, and figs. S8-S9.

***Analysis of high- vs. low-genetic interaction degree essential genes.*** To understand properties associated with essential genes that exhibited high vs. low genetic interaction degree in our screens, we tested for differences between these two groups in terms of the same physiological, evolutionary and functional gene properties described above (see Section Features associated with HAP1 essential genes)(Data File S14). First, we formed a group of "high-density" essential genes (among the set of all library genes) by identifying the top 20% of HAP1 essential genes in terms of total genetic interaction degree. We formed a group of "low-density" essential genes by identifying the bottom 50% of HAP1 essential genes in terms of total genetic interaction degree. The continuous-valued and discrete-valued features listed in Data File S14 were then used to test for differences between these two groups with a Wilcoxon rank-sum test. For each feature, the mean of the high-density group and the low-density group was computed, and the $\log_2$ ratio of these means was plotted in fig. S10B, with and without the mitochondrial gene set (Data File S2).

### Genetic interactions involving paralogs

***Genetic interactions within pairs of paralog genes.*** A list of paralog gene pairs was obtained from the Ensembl database {Harrison, 2023 #6286}. In addition, a list of paralog genes classified as "Ohnolog" duplicated gene pairs was also obtained {Singh, 2020 #6282} . To evaluate the density of positive and negative genetic interactions observed between paralogs as a function of sequence identity, three subsets of Ensembl paralog pairs were selected based on the HAP1 expression level of each gene in a pair (Data File S2): [1] Paralog pairs in which both genes had expression scores of $\log_2$ (TPM+1) > 1, [2] Paralog pairs where exactly one gene met this expression criteria, and [3] Paralog pairs where neither gene met the expression criteria. Paralogs were further binned on the basis of their sequence identity to each other; for each sequence ID bin, the sets were filtered to include only pairs having a sequence identity greater than the bin and those pairs screened in our study for a genetic interaction. Finally, the percentage of positive and negative genetic interactions identified at a standard threshold (|qGI| > 0.3 and FDR < 10%) within the filtered set were calculated per bin for all three described subsets. To calculate the density of negative genetic interactions as a function of paralog family size, a subset of paralog pairs was selected for analysis by requiring expression in HAP1 [($log_2$ (TPM+1) > 1] for both paralogs and requiring that the paralogs' sequence identity was greater than the indicated cut-off (20%, 50% or Ohnolog set). The family size of each paralog pair was then recalculated with respect to only paralogs that satisfied the expression and sequence identity criteria. Finally, the density of negative genetic interactions (determined by standard cut-off of qGI < -0.3 and FDR < 10%) within the filtered set was calculated among the set of pairs screened in our study and reported for each family size bin. The results of this analysis are shown in Fig. 3F-G, fig. S11A-C.

***Overall genetic interaction density of paralog genes.*** To calculate the overall density of negative and positive genetic interaction for paralog library genes with all query genes as a function of paralog family size, a subset of paralog pairs was selected for analysis by requiring expression in HAP1 [$log_2$ (TPM +1) > 1] for both paralogs and requiring that the paralogs' sequence identity was greater than the indicated cut-off (20%, 50% or Ohnolog set). The family size of each paralog pair was then recalculated with respect to only paralogs that satisfied the expression and sequence identity criteria. Finally, the density of library paralog gene negative and positive genetic interactions (|qGI| > 0.3 and FDR < 0.1) with all possible query genes tested within the filtered set was calculated and reported for each family size bin. We also measured the negative and positive interaction density associated with duplicated query genes. Negative and positive interaction density was measured for query genes paralogs that shared greater than 20% sequence identity as well as Ohnolog genes that were screened as a query in this study. Query gene genetic interaction density reflects the number of negative or positive genetic interactions identified at a standard confidence threshold (|qGI| > 0.3, FDR < 0.1) per query gene divided by the total number of tested gene pairs involving the corresponding query gene. Paralog query gene interaction density was compared to negative and positive interaction density associated with non-duplicated query genes screened in this study. Results corresponding to this analysis are shown in fig. S11D-E.

***Paralog degree asymmetry analysis.*** We compared the negative genetic interaction degree associated with each gene of a paralog pair (i.e. the number of negative genetic interactions each paralog exhibited with all tested query genes). Gene pairs from the Ensembl paralog pair set were selected for this analysis where both genes were represented as library gene in the non-redundant genetic interaction dataset (Data File S4), both satisfied a HAP1 expression cutoff of $log_2$(TPM+1) > 1, and they represented paralog pair with sequence identity greater than 20% or 50%. Only paralog pairs with a negative degree (standard cut-offs of |qGI| > 0.3 and FDR < 10%) of greater than 4 were considered in this analysis. For each pair of library gene paralogs, we measured a negative interaction degree ratio statistic, computed by dividing the negative interaction degree associated with the paralog showing the higher interaction degree by the interaction degree of the sister paralog with fewer negative interactions. Ratios that were greater than 30 were assigned a value of 30. We used simulations to establish a null distribution for this ratio statistic that assumed genetic interactions would accumulate with equal probability for each gene within a paralog pair (i.e. symmetric negative interaction degree). Statistics for random paralog pairs under this null model were created by randomly generating negative interaction degrees for each gene pair from a binomial distribution with probability of 0.5 and the maximum limit of total observed degree for each real paralog pair. One thousand different random scenarios were generated, with ratio statistics measured for all random paralog pairs in each scenario. An empirical *P*-value was generated using the R ‘qvalue’ package (empPvals function) by comparing the observed average degree ratio and the average degree ratios from 1000 random models. The same analysis was repeated using the Ohnolog paralog list. Results from this analysis are shown in fig. S11F.

### Overlap of genetic interactions with other genomic datasets

We characterized the functional relationships between pairs of genes connected by negative or positive genetic interactions by evaluating their overlap with other types of molecular and functional interactions, described below. The results of this analysis are show in Fig. 4A and fig. S12A.

***Protein-protein interactions.*** Protein-Protein interactions (PPI) were collected from three sources and combined to generate a unified PPI standard that was compared to the HAP1 genetic interaction dataset. These included HuRI {Luck, 2017 #6018}, BioPlex {Huttlin, 2021 #5655}, and huMAP {Drew, 2017 #5910}. All possible pairs of proteins within this set that did not demonstrate a PPI in any of the datasets were assumed to be non-interacting for the purposes of this analysis.

**Gene Ontology (GO) biological process co-annotated gene pairs.** GO term annotations were downloaded in June of 2021 {Harris, 2004 #4357} and a functional standard for co-annotation to GO Biological Process (GO BP) terms was built using the FLEX software {Rahman, 2021 #5666}. Briefly, a set of GO BP terms was curated to include only those terms that contained between 2 and 300 genes. Terms of this size were considered to have enough specificity to be functionally informative. Two genes were considered functionally related if they were co-annotated to one or more of these GO BP terms. Two genes were considered unrelated if both genes were annotated to at least one GO term but the lowest common ancestor to which both genes were annotated was not part of the functionally specific set of GO bioprocess terms defined above. All other gene pairs (e.g., those with one or both unannotated genes), were ignored for the purposes of co-annotation analysis.

***Co-expressed gene pairs.*** We measured gene co-expression using the Cancer Cell Line Encyclopedia (CCLE) dataset {Ghandi, 2019 #4805}. To generate a set of co-expressed gene pairs, expression profile similarities were calculated using Pearson correlation (PCC) for all possible gene pairs. The top 0.1% of all the gene pairs (according to PCC values) were considered as co-expressed and the rest were not considered to be co-expressed.

***Co-localized protein pairs.*** Gene product pairs were considered co-localized if they shared one or more cellular compartment annotation(s) based on a standard we derived from the COMPARTMENTS Subcellular localization database retrieved in July 2021 {Binder, 2014 #6288}. Specifically, we applied a high-confidence cutoff of > 4.5/5 and focused on the following subset of 9 terms: Nucleus, Nucleolus, Golgi apparatus, Mitochondrion, Peroxisome, Endoplasmic reticulum, Lysosome, Endosome, Cytoskeleton & Plasma membrane.

***Co-complex annotated protein pairs.*** Co-complex pairs were derived from the CORUM protein complex standard (release 3.0){Tsitsiridis, 2022 #5807}. Any pair of genes whose products were members of the same complex was treated as a positive example while any gene pair for which each of the gene products were annotated to at least one complex, but not the same complex, was treated as a negative example.

***<u>Co-pathway annotated gene pairs.</u>*** Co-pathway pairs were derived from the Molecular Signatures Database (MSIGDB){Liberzon, 2011 #4481}. Any pair of genes whose products were members of the same MSIGDB pathway was treated as a positive example while any gene pair for which each of the gene products were annotated to at least one pathway, but not the same pathway, was treated as a negative example.

### Precision-recall analysis

All precision-recall analyses were performed using the FLEX R package {Rahman, 2021 #5666} along with the corresponding GO BP co-annotation (GO BP) or CORUM co-complex functional standards, described above. Results of precision-recall analyses are shown in Fig. 9F and figs. S5B, S13A and S20F.

### Genetic interactions within and between protein complexes

***<u>Genetic interaction enrichment within and between protein complexes</u>***. We performed tests for enrichment of genetic interactions that occur within and between protein complexes annotated to the CORUM protein complex database {Tsitsiridis, 2022 #5807}.

*<u>Within-complex enrichment analysis</u>*: tests if gene pairs connecting genes that encode members of the same protein complex are enriched for negative and/or positive genetic interactions.

*<u>Between-complex enrichment analysis:</u>* tests if gene pairs encoding members of two different complexes (i.e. interaction between a gene annotated to complex 1 and a second gene annotated to complex 2) are enriched for negative and or positive genetic interactions.

For these analyses, we considered only those CORUM protein complex where at least three genes were represented in our library gene set and at least one gene was represented in the query gene set. A standard threshold (|qGI| > 0.3 and FDR < 0.1) was applied to identify significant negative and positive genetic interactions. When computing enrichment statistics, the library gene universe was reduced to only those genes that overlap with the CORUM complex standard genes, and all query genes in the non-redundant genetic interaction dataset (Data File S1, and S4) were included in the analysis. Enrichments were calculated independently for positive, negative and total interactions (combined positive and negative) genetic interactions. *P*-values were derived from one-tailed hypergeometric tests, focusing on increased genetic interaction density relative to the background genetic interaction density. *P*-values were corrected using the Benjamini-Hochberg method applied to the within-complex and between-complex sets separately. After correction, any enrichment driven by a single interaction was marked insignificant by setting the *P*-value to 1. Genetic interaction enrichment observed within and between CORUM-annotated protein complexes is provided in Data File S15. Results pertaining to genetic interaction enrichment within and between protein complexes is shown in Fig. 4 and fig. S12.

***<u>Genetic interaction purity score.</u>*** The genetic interaction purity score refers to the fraction of positive or negative genetic interactions relative to the total number of observed genetic interactions within the same complex or between a pair of different complexes {Baryshnikova, 2010 #2405}{Costanzo, 2016 #4354}{Segrè, 2004 #5876}. The genetic interaction purity score was normalized to the range of [-1,1], where -1 indicates that all genetic interactions within a specific complex or between a particular pair of complexes are exclusively negative. A purity score of 1 indicates all genetic interactions observed with a complex or between a pair of complexes are all positive.

The genetic interaction purity score distribution plots, shown Fig. 4D and fig. S12B summarize the genetic interaction purity for protein complexes consisting of more than 5 screened pairs and that are enriched for genetic interactions (FDR < 0.10, hypergeometric test, Benjamini-Hochberg-corrected). To generate a null (background) purity score distribution, a random positive genetic interaction count was generated for each complex from a binomial distribution given the number of total genetic interactions in the complex and the background probability of observing a positive interaction (fraction of positive genetic interactions measured across all complexes). This analysis was completed both within and without the mitochondrial gene set. In fig.S12, a set of smaller complexes that were subsets of larger complexes were excluded from the analysis results. This reduced complex set was formed based on overlap index scores computed for all pairs of complexes in the CORUM standard. The overlap index between two complexes is the number of shared genes divided by the size of the smaller complex. An overlap-index cut-off of 0.3 was applied here such that no two complexes with an overlap index of ≥ 0.3 were both included in the non-redundant set. To observe the contribution of mitochondria-related complexes to this result, all analyses were repeated after removing a set of complexes for which more than 50% member genes are mitochondrial. Genetic interaction purity scores are provided in Data File S15. Results pertaining to genetic interaction purity scores within and between protein complexes are shown in Fig. 4 and fig. S12.

### A genetic interaction profile similarity-derived functional hierarchy

***Constructing a functional hierarchy.*** As described above (Hierarchical clustering of HAP1 genetic interaction profiles), we modified a previously described hierarchical clustering algorithm to the complete genetic interaction dataset to produce a set of uniformly sized gene clusters across layers of the dendrogram (Data File S9){Xiong, 2024 #6365}(https://doi.org/10.5281/zenodo.15320010). The same algorithm was also used to cluster the subset of 3784 genes derived from the same centroid-corrected, genetic interaction profile similarity network shown in Fig. 2 and described above (Constructing a genetic interaction profile similarity network) to build a genetic interaction profile-based hierarchy of gene function (Data File S16). Initial clustering organized these genes into 4 levels of hierarchically organized modules. Gene modules identified in level 4 of the hierarchy represented parent clusters in the hierarchy, which were separated into smaller child clusters in lower hierarchical levels, with level 1 of the hierarchy consisting of the smallest child gene clusters. Genes that did not belong to a functionally enriched cluster identified at level 2 of the hierarchy, were removed from the analysis. Functionally enrichment was defined as level 2 clusters whose gene members were statistically enriched (FDR < 0.1, hypergeometric test, Benjamini-Hochberg-corrected) for one or more GO bioprocess terms {Yu, 2012 #6367}. The resultant functional hierarchy consisted of 1,863 genes associated with functionally rich genetic interaction profiles. These genes were organized into 6 large parent clusters at level 4 of the hierarchy, 18 child clusters of level 3 of the hierarchy, 181 level 2 clusters and finally, 593 level 1 clusters (Data File S16). Importantly all

1,863 genes were represented at each level of the functional hierarchy and a set of 98 genes that did not belong to a cluster at each of the 4 hierarchical levels were excluded from this analysis. This was a critical filtering step because it ensured that the same set of genes was examined at all levels of hierarchical study and that any differences observed (e.g. in genetic interaction density) were due to hierarchical structure at different levels and not differing sets of gene involved in the hierarchy at each level. As a result of these procedures: [1] all genes were members of only one cluster at each level of the hierarchy; [2] all clusters were strict subsets of their parent clusters one level above, and [3] each cluster was comprised of the union of its children clusters one level below. In addition, at the fourth layer, we identified one cluster including 292 genes that are associated with mitochondrial functions, which was later excluded for some analyses (where noted in Data File 16).

To functionally characterize each level of the hierarchy, we performed enrichment analysis for co-annotated genes within the clusters at each layer of the hierarchy using the CORUM {Tsitsiridis, 2022 #5807}, GO BP {Ashburner, 2000 #2281}, and cellular compartment {Binder, 2014 #6288} functional standards. At each layer, we calculated the fraction of gene pairs within each cluster that are co-annotated to the same term in a functional standard relative to the population of all possible gene pairs (i.e. 1,863 choose 2 = 1,734,453) and then performed a hypergeometric test to assess enrichment of genes within the same cluster to be co-annotated to the same functional term. Note that to distinguish functional signals that were specific and unique to each level of the hierarchy, the hypergeometric test was performed using only the additional gene pairs added by each hierarchical layer compared to the preceding layer. Specifically, this includes those gene pairs that are newly co-clustered at a given level of the hierarchy that were not previously co-clustered based on the descendant clusters. These sets of gene pairs are referred to as the "shell" at each level of the hierarchy, and enrichment fold-change and *P*-values are reported for these sets. This analysis showed that parent clusters at the fourth level of the hierarchy were specifically enriched for distinct subcellular compartments. Smaller level 3 clusters were enriched for more specific but distinct GO bioprocess terms while clusters identified at levels 1 and 2 of the hierarchy were uniquely enriched for distinct biological pathways and protein complexes (fig. S13B, Data File S16). This analysis was also repeated after excluding 292 mitochondrial genes Data File S16. Analyses pertaining to the genetic interaction profile-based functional hierarchy are shown in Fig. 5 and fig. S13.

***Measuring interaction density and magnitude at different levels of the functional hierarchy.*** The functional hierarchy was used to assess the density of genetic interactions at varying degrees of functional relatedness, where relatedness was defined by the hierarchy level described above. For each screened pair of genes, we noted whether or not it shared a genetic interaction and a level of functional relatedness of that pair, as defined by the most specific level of the hierarchy in which they shared membership to a common cluster. We used this measure for calculating interaction density at different levels of functional relatedness and assigned each pair to the level of their most specific common cluster. This prevented the density estimate at each level from being dominated by the trends observed at lower levels. Density was defined as the number of significant interactions meeting our standard significance threshold (|qGI| > 0.3; FDR < 0.1), divided by the number of screened pairs. Similarly, after binning all pairs of genes by their qGI score magnitude, we measured the fraction of pairs in each bin belonging to each hierarchy level using the most specific level available for each pair. To alleviate the potential bias introduced by mitochondrial genes, we also performed the same measurement excluding 292 mitochondrial genes identified in one of the 6 large clusters at the fourth layer. The results from this analysis are shown in Fig. 5 and fig. S13.

***Genetic interaction enrichment within and between biological processes.*** We performed tests for enrichment of genetic interactions that occur within and between different biological process-level gene sets. Gene sets representing functionally coherent groups of genes for this analysis were based on the domains described in the section above "*Biological process annotation of the genetic interaction profile similarity network*". For each domain, we compiled the list of library genes and query genes associated with that domain. For these analyses, we considered only those domains where at least three genes were represented in our library gene set and at least one gene was represented in the query gene set, which resulted in a total of 15 domains (Data File S16). We tested each domain for enrichment of genetic interactions (within-domain analysis), and we also tested each pair of domains for enrichment of genetic interactions (between-domain analysis).

For each within-domain or between-domain gene pair set, the enrichment test was completed as follows. First, a standard threshold (|qGI| > 0.3 and FDR < 0.1) was applied to identify significant negative and positive genetic interactions. When computing enrichment statistics, the library gene universe was reduced to only those genes that overlapped the domains of interest, and the query gene universe was also restricted to only those genes that overlapped the domains of interest. Enrichments were calculated independently for positive, negative and total interactions (combined positive and negative) genetic interactions. *P*-values were derived from one-tailed hypergeometric tests, focusing on increased genetic interaction density relative to the background genetic interaction density. *P*-values were corrected using the Benjamini-Hochberg method applied to the within-domain and between-domain sets separately. After correction, any enrichment driven by a single interaction was marked insignificant by setting the p-value to 1. Genetic interaction enrichment observed within and between these bioprocess-level domains is provided in Data File S16, and the related figure is plotted in Fig. 5D.

### Genetic Suppression Analysis

***Suppression score.*** To define the subset of positive genetic interactions that may represent instances of genetic suppression, we formulated a suppression score for all positive interactions that satisfied the standard genetic interaction significance threshold (qGI > 0.3,FDR < 0.1).

$$S_{ij} = \frac{DMF_{ij} - SMF_{min}}{|SMF_{min}|}, \quad if\ SMF_{min} < (-0.9)\ \&\ SMF_{min} < DMF_{ij}$$
$$S_{ij} = 0,\ otherwise$$
$$SMF_{min} = minimum(L_i, Q_j)$$
$$DMF_{ij} = L_i + Q_j + qGI_{ij}$$

Where $L$ is the library gene single mutant fitness (in $\log_2$ fold-change) and $Q$ is the query gene single mutant fitness (in $\log_2$ fold-change). Suppression scores for all positive interacting gene pairs are provided in Data File S17. Results related to genetic suppression analysis are shown in Fig. 6A and fig. S14A-C.

***Functional enrichment analysis of genetic suppression interactions.*** To assess if genetic suppression gene pairs were more functionally related compared to gene pairs connected by positive genetic interactions in general, positive genetic interactions were sorted based suppression scores. FLEX{Rahman, 2021 #5666} was then used to compute the precision of identifying gene pairs annotated to the same GO bioprocess term based on suppression scores at two different thresholds (suppression score > 0.5 and > 0.8) and for all positive genetic interactions (qGI > 0.3, FDR <0.1 and suppression score ≥ 0). Fold-enrichment for co-annotated gene pairs was calculated by dividing precision determined at the suppression score thresholds described above by the background precision (size of positive class divided by total number of gene pairs). The *P*-values comparing each set relative to background precision were determined by Fisher-exact tests. This analysis was also repeated after excluding mitochondrial genes. These results of this analysis are shown in fig. S14D.

***HAP1 essential gene suppression interactions.*** We examined if HAP1 essential genes involved in genetic suppression interactions where more likely to be considered core or context-dependent essential genes based on the DepMap 20Q2 dataset (DepMap; depmap.org/portal). Positive interactions with a suppression score greater than 0.5 were defined as genetic suppression interactions. HAP1 essential genes were grouped into the following categories: [1] all HAP1 essential genes, [2] HAP1 essential genes involved in at least one suppression interaction, [3] HAP1 essential genes with positive genetic interactions, and [4] HAP1 essential genes with no suppression interactions. We then determined the percentage of cancer cell lines in the DepMap 20Q2 dataset that also depended on HAP1 essential genes from each category for viability. Cancer cell line essential genes were defined as genes that met a CERES score threshold of < -1, which corresponds to the median score of all essential genes in the DepMap data. Statistical significance was determined by Wilcoxon-rank sum tests between each pair of HAP1 essential gene groups. The same analysis was repeated excluding the mitochondria gene set. The results of this analysis are shown in Fig. 6B and fig. S14E.

### Genetic interaction conservation

The following methods were used to evaluate conservation of genetic interactions between human HAP1 cells and the budding yeast, *S. cerevisiae*. For gene pairs screened more than once, we considered a genetic interaction if at least half of the screened pairs resulted in a significant genetic interaction score (|qGI| > 0.3, FDR < 0.1).

***Conservation of genetic interaction density between biological processes.*** We classified human and *S. cerevisiae* genes into 15 distinct bioprocess functional groups (Data File S18). For each pair of bioprocess groups, we calculated the negative and positive genetic interaction density between the corresponding gene sets in the HAP1 genetic network described in this study and the *S. cerevisiae* genetic interaction network {Costanzo, 2016 #4354}. Genes annotated to more than one bioprocess group were excluded from the analysis, and we only considered pairs of bioprocesses for which at least 100 gene pairs were tested for genetic interactions. Finally, we calculated the Pearson's correlation coefficient between the interaction density obtained in the HAP1 and *S. cerevisiae* genetic networks. The results of this analysis are provided in Data File S18 and shown in Figure 7A and figure S15A.

***Conservation of genetic interactions between human-yeast ortholog gene pairs.*** We used DIOPT (https://fgr.hms.harvard.edu/diopt){Hu, 2011 #5815} to retrieve orthology mappings between human and *S. cerevisiae* genes and selected genes with 1:1 orthologs between both species or gene with more than one ortholog in human (i.e. N:1 orthology relationships, such that 1 gene in *S. cerevisiae* has N orthologs in human). Of all the gene pairs tested in HAP1 cells, 3.7% involved such selected genes. Next, we selected the gene pairs that were tested for genetic interactions in both HAP1 cells and *S. cerevisiae* and counted the number of genetic interactions identified only in HAP1 cells (i.e. HAP1-specific genetic interactions) and in both species (i.e. conserved genetic interactions). The results of this analysis are provided in Data File S20 and shown in figure S16A.

***Conservation of genetic interaction profile similarity human-yeast orthologs.*** We measured the genetic interaction profile similarity for human gene pairs as explained above (see section, Mapping the genetic interaction profile similarity network). Genetic interaction profile similarity values for *S. cerevisiae* gene pairs were downloaded from thecellmap.org {Usaj, 2017 #4404}{Costanzo, 2016 #4354}. For yeast gene pairs associated with multiple temperature-sensitive mutant alleles, the correlation values were averaged. We only considered gene pairs involving genes with a 1:1 or N:1 human to *S. cerevisiae* orthology relationship for which profile similarity data was available in both species. Next, we retrieved gene pairs within the top 5% of HAP1 profile similarity, and determined the fraction of those pairs whose orthologs were within the top 5% similarity values in the *S. cerevisiae* genetic interaction profile similarity network. We calculated the same fraction for the remaining 95% of gene pairs in the HAP1 profile similarity network. Statistical significance was calculated using Fisher's exact test. The results of this analysis are show in figure S17A-B.

***Genetic interaction enrichment analysis among yeast-human conserved genes.*** We first selected gene pairs tested for genetic interactions in HAP1 cells whose ortholog pairs (with 1:1 or N:1 orthology relationships, see above) were also tested in *S. cerevisiae* {Costanzo, 2016 #4354}. For the sets of gene pairs with and without a genetic interaction in HAP1 cells, we calculated the fraction that showed a corresponding genetic interaction in *S. cerevisiae*. We derived the fold enrichment in the conservation of genetic interactions in yeast by calculating the ratio between both fractions and calculated the statistical significance using Fisher's exact test.

To calculate the conservation of genetic interaction in HAP1 cells, we used the same approach but starting from gene pair sets with and without a genetic interaction in *S. cerevisiae* and calculated the fraction of pairs which had a genetic interaction in HAP1 cells. We repeated the same calculations using: [1] 100 randomized genetic interaction networks respecting the network topology and compared the obtained fold enrichments to the result in the real genetic interaction network to obtain an empirical *P*-value; [2] Process-specific genetic interaction networks involving only pairs with at least a gene annotated to the corresponding biological process.

We followed the same approach to examine the conservation between *hPTAR1* and *yECM9* genetic interactions but restricted the gene pairs in HAP1 cells to those involving *PTAR1*, and the gene pairs in *S. cerevisiae* to those involving *ECM9*. *ECM9* genetic interactions were identified using SGA analysis described in the next section. Only reproducible *ECM9* genetic interactions identified in independent biological replicate screens were considered in this analysis. The results of these analyses are provided in Data File S20 shown in Figures 7B-D and figure S16B-C.

### Follow-up Experiments

***mTOR activity assays.*** For figure S15C, wild-type and query mutant cell lines (*ANGPTL4*, *EMC6*, *FANCA*, *GSK3A*, *SLX1*, *C1orf112*, *PDCD5*, *TAPT1*, *VPS52*) were cultured in IMDM (Gibco), 10% FBS (Gibco) and 1X penicillin and streptomycin (Gibco) at 37°C with 5% $CO_2$. For immunoblots, 250k cells seeded in 12-well plates were lysed at 70-80% confluency directly in well using RIPA buffer [50 mM Tris-HCl (pH8.0), 150 mM NaCl, 2 mM EDTA, 1% NP-40, 0.5% sodium deoxycholate, 0.1% SDS with 1X Halt Protease and Phosphatase Inhibitor Cocktail (Thermo Fisher)]. 10 µg total protein was mixed with 6X loading buffer [375 mM Tris-HCl (pH7.4), 50% Glycerol, 6% SDS, 30% 2-Mercaptoethanol, 0.04% Bromophenol blue], and incubated at 95°C for 5 min. Samples were run on pre-cast gels (Bio-rad) and transferred to Nitrocellulose membranes using iBlot 2 Gel Transfer Device (Thermo Fisher). Membranes were blocked in 5% BSA/TBS-T and incubated overnight at 4°C with primary antibody against phospho-S6 Ribosomal Protein Ser240/244 (CST, #2215), S6 Ribosomal Protein (CST, #2317), phospho-Akt Ser473 (CST, #4060), and Akt (CST, #2920). Secondary antibody incubation was done for 1 hour at room temperature in dark using donkey anti-mouse IRDye® 680RD (Li-Cor, 926-68072) and donkey anti-rabbit IRDye® 800CW (Li-Cor, 926-32213). Membranes were visualized with Li-Cor Odyssey Classic instrument and ImageStudio was used for signal quantification. GraphPad Prism v.10 was used for statistical analyses.

For figures S15D-E, HAP1 wild-type and query mutant cell lines (*GSK3A*, *CDKN2B*, *ANGPTL4*) were seeded at a density of 500,000 cells/well in a 6-well plate and cultured for 24 hours. The following day, cells were treated with 250 nM rapamycin (Selleckchem, Cat.#S1039) for 24 hours. On day 2, cells were subjected to amino acid starvation for 50 min using amino acid-free DMEM (Wisent, Cat.#319-004-CL), followed by amino acid refeeding for 10 min. Cells were then washed twice with ice-cold PBS and lysed on ice using RIPA lysis and extraction buffer (Thermo Fisher Scientific, Cat.# 89901) supplemented with Halt Protease and Phosphatase Inhibitor Cocktail (Thermo Fisher Scientific, Cat.#23227). Protein concentrations were quantified using the Pierce BCA Protein Assay Kit (Thermo Fisher Scientific, Cat.#23227). Cell lysates were mixed with NuPAGE LDS sample buffer (Invitrogen, Cat.# NP0007) and denatured at 95°C for 5 min. Proteins were separated using NuPAGE 4-12% Bis-Tris Mini Protein Gels (Invitrogen, Cat.#NP0323BOX) and transferred for immunoblotting. Primary and secondary antibodies and dilutions were as follows: Akt(pan)(40D4) mouse mAb at 1:1000 (Cell Signaling, Cat.#2920; RRID: AB_1147620); phospho-Akt1(Ser473)(D7F10) XP rabbit mAb at 1:1000 (Cell Signaling, Cat.#9018; RRID: AB2629283); p70 S6 kinase (49D7) rabbit mAb at 1:1000 (Cell Signaling, Cat.# 2708; RRID: AB_390722); phospho-p70 S6 kinase (Thr389) (108D2) rabbit mAb at 1:1000 (Cell Signaling, Cat.#9234, RRID: AB_2269803); β-actin (8H10D10) mouse mAb at 1:10,000 (Cell Signaling, Cat.#3700; RRID:AB_2242334); goat anti-mouse IgG (H+L) cross-adsorbed secondary Ab, HRP (Invitrogen, Cat.#A16072; RRID: AB_2534745); and goat anti-rabbit (H+L)-HRP conjugate (Bio-Rad, Cat.#170-6515; RRID: AB_11125142). Chemiluminescence was detected using the SuperSignal West Pico PLUS Chemiluminescence Substrate (Thermo Fisher Scientific, Cat.#34580) and images were acquired using the iBright FL1500 Imaging System (Invitrogen) in chemiluminescent mode.

***yECM9/hPTAR1*** **functional studies.** The following sections describe methods to validate *ECM9* as a yeast palmitoyltransferase important for prenylation and activation of Ykt6. The results from these experiments are shown in Fig. 7E-I and fig. S17E.

Yeast strains. All strains used for SGA, yeast two hybrid analysis and complementation assays were derivatives of BY4741 or Y7092, the construction of which was described previously {Tong, 2006 #1101}. A temperature-sensitive allele of *ECM9*, TSQ3052 (*MATα ecm9-5002::natMX can1Δ::STE2pr-Sp_his5; lyp1Δ; his3Δ1 leu2Δ0 ura3Δ0 LYS2*) was used as a query mutant for SGA analysis. Yeast complementation analysis was performed using Y16258 (*MAT* ***a****/α ECM9/ecm9Δ::natMX CAN1/can1Δ::STE2pr-Sp_his5; LYP1/lyp1Δ; HIS3/his3Δ1 LEU2/leu2Δ0 URA3/ura3Δ0*) or Y15767 (*MATα ecm9Δ::natMX abh1Δ::URA3 can1Δ::STE2pr-Sp_his5; lyp1Δ; his3Δ1 leu2Δ0 ura3Δ0 LYS2*) as indicated below.

Yeast Two Hybrid Analysis. Genes encoding the alpha and beta subunits of farnesyltransferase, and geranylgeranyl transferase type I-III were amplified by PCR from *S. cerevisiae* genomic DNA, using sequence specific primers fused to common sequences used for homologous recombination cloning {Tonikian, 2009 #2493}. Genes were cloned into DNA binding domain- and activation domain-fusion plasmids and two hybrid analysis was performed as described elsewhere {Tonikian, 2009 #2493}.

Synthetic Genetic Array (SGA) analysis to map genetic interactions for yeast *ECM9*. We constructed a hypomorphic, temperature-sensitive (TS) mutant of the yeast essential gene, *ECM9* (TSQ3052), in the SGA query strain background and screened it for genetic interactions against ordered arrays of 3,827 nonessential yeast deletion mutants and 786 TS alleles representing 560 yeast essential genes, as previously described {Costanzo, 2016 #4354}{Kuzmin, 2014 #4234}. All SGA selection steps were conducted at permissive temperature (22°C) except for the final selection of haploid double mutants, which were incubated at a semi-permissive temperature (26°C) prior to imaging. Genetic interactions were identified and measured using the quantitative SGA genetic interaction score, as described elsewhere {Baryshnikova, 2010 #2405}{Costanzo, 2016 #4354}. *ECM9* genetic interactions are provided in Data File S20.

*PTAR1-RABGGTB complementation assay.* An *ECM9* heterozygous deletion strain (Y16258) harboring a low-copy vector control plasmid with a *URA3* selection marker (plasmid P13744) or the same plasmid expressing human *PTAR1* and *RABGGTB* (plasmid P13719) genes from the galactose-inducible *GAL1* promoter were sporulated in liquid sporulation medium for 1 week, as described elsewhere {Kuzmin, 2014 #4234}. Asci were digested with Zymolyase and tetrads were dissected onto YEPD and YEPGalactose and grown 4 days at 30°C. Plates were photographed and replica plated onto selective YEPD or YEPGalactose medium supplemented with neurothrecin as well as synthetic defined medium lacking uracil to follow the segregation patterns of knockout alleles relative to the fitness phenotype, as described elsewhere {Dowell, 2010 #1627}.

*ABHD16A and ABHD17B complementation assay.* Overnight cultures of an *ecm9Δabh1Δ* double deletion mutant (Y15767) harboring a low-copy vector control plasmid with a *URA3* selection marker or the same plasmid expressing human *ABHD16A* (plasmid P13708), human *ABHD17B* (plasmid P13712) or yeast *ABH1* (P13713) genes from the galactose-inducible *GAL1* promoter, were serially diluted and spotted onto selective agar media lacking uracil supplemented with 2% glucose or 2% galactose as indicated. Strains were grown for 2 days at 30°C and imaged.

*Palmitoylation Detection by mPEG replacement chemistry.* Denatured, whole-cell yeast protein extracts were subjected to three chemical steps to replace thioester-linked palmitoyl-modifications with the methoxypolyethylene glycol maleimide (mPEG) (Sigma-Aldrich 712469): [1] blockade of free thiols with N-ethylmaleimide (NEM), [2] treatment with neutral pH hydroxylamine to release thioester-linker modifications (e.g. palmitoylation), restoring thus the cysteinyl thiol and [3] modifying the newly exposed thiols with the 10 kDa thiol-reactive mPEG reagent, resulting in a size-shift for proteins having acylated cysteines. In practice, Steps 2 and 3 were combined, i.e. hydroxylamine treatment and mPEG reaction were done concurrently. The treated extracts were subjected to Tricine SDS-PAGE and then Western blotting, using a rabbit polyclonal anti-Ykt6 antiserum and aHRP-conjugated goat anti-rabbit secondary antibody. Final visualization was via enhanced chemiluminescence (ECL) and x-ray film exposure.

The early steps of the acyl-mPEG exchange protocol, closely track the previously described small-scale acyl-biotinyl exchange (ABE) protocol {Roth, 2006 #6363}. 1 x $10^8$ cells were harvested by centrifugation from log-phase yeast cultures growing in YPD media (1% yeast extract, 2% peptone, 2% glucose) with the cell pellet resuspended into ice-cold 200 ul Lysis Buffer (LB: 150 mM NaCl, 50 mM Tris/Cl, 5 mM EDTA pH 7.4), supplemented with 10 mM NEM (Sigma) and 2xPI (1xPI:

1mM PMSF, and 0.25 mg/ml each of antipain, leupeptin, pepstatin, and chymostatin). A 200 ul volume of glass beads (Sigma) was added and lysis was effected with five 45 sec intervals of vigorous vortex mixing, interspersed with 2 min rest periods on ice. The lysate was decanted away and combined with a 300 ul LB/10 mM NEM/1xPI quick wash of the beads. To solubilize cellular membranes, Triton X-100 (Anatrace) was added to 1.7% and the lysates were subjected to gentle mixing at 4°C. Protein from a 150 µl portion of the lysate was collected by chloroform-methanol (CM) precipitation {Wan, 2007 #6355}. The scaled-down version of the CM precipitation used here involves 8-fold reduced volumes relative to the previously detailed CM precipitation (Wan et al., 2007), allowing the use of 1.5 ml screw-cap centrifuge tubes and a microfuge. The resulting protein pellet was solubilized with 30 ul SDS Buffer (SB: 4% SDS, 50 mM Tris/Cl, 5 mM EDTA, pH 7.4) with 10 mM NEM and incubated at 37°C for 10 min to denature protein. The sample was then diluted with 120 ul LB supplemented with 1 mM NEM, 0.2% Triton X-100, and 1xPI and further incubated for 60 min at 4°C. Three sequential CM precipitations were then used to fully remove residual NEM. After the first two precipitations, protein pellets were dissolved into 30 ul SB and then diluted with 120 ul LB with 0.2% Triton X-100. Following the third CM precipitation, protein pellets were dissolved in 70 ul SB, which was used for the two experimental conditions, + and –mPEG. For the +mPEG condition, a 15 ul of the sample was diluted into 60 ul 0.8 M hydroxylamine (Sigma-Aldrich 467804), 150 mM NaCl, 2 mM mPEG, 0.2% Triton X-100, 1xPI. For the –mPEG control condition, 15 ul was diluted into 60 ul 0.8 M hydroxylamine, 150 mM NaCl, 0.2% Triton X-100, 1xPI. Samples were incubated at room temperature with gentle rotation for 1 hour. Finally, 75 ul LB was added and samples were subjected to a final CM precipitation with the protein pellet being dissolved into sample buffer for gel loading.

*HAP1 chemical-genetic interaction profiling of the small molecule, ABD957.* We performed a genome-wide chemical-genetic interaction screens with ABD957, a small molecule inhibitor of the *ABHD17* subset of depalmitoylases. Chemical-genetic interaction screens were performed as described above (see section Annotating functions using the HAP1 interaction profile similarity network) and elsewhere {Masud, 2022 #5659}. ABD957 negative and positive chemical-genetic interactions were scored as previously described {Lin, 2024 #6328}.

#### Analysis of genetic interactions underlying cancer gene dependencies

***Expression dependency (ED) score analysis.*** For each unique gene pair tested in the HAP1 nonredundant genetic interaction dataset (Data File S4), we extracted CRISPR gene effect fitness scores (Chronos score, DepMap 22Q4) and gene expression [$\log_2$(TPM+1)] derived from a panel of cancer cell lines examined in the DepMap 22Q4 dataset (depmap.org/portal). We then measured the correlation (PCC) between cancer cell line fitness scores associated with one gene and expression of the second gene across the same panel of cancer cell lines. In doing so, we generated two PCC and *P*-values using scipy.stats.pearsonr {Virtanen, 2020 #6375} for each pair of genes tested based on Gene A fitness-Gene B expression and Gene A_expression-Gene B fitness. Gene pairs were filtered based on correlation and statistical thresholds (|PCC|>0.1 and *P*-value < 0.01)(Data File S21). Reciprocal gene pairs that satisfied these thresholds but exhibited PCC values of opposite signs were also removed. We referred to the correlation between gene pair fitness and expression measured across a panel of cancer cell lines as a gene pair Expression Dependency (ED) score.

***Comparing expression dependency (ED) and quantitative genetic interaction (qGI) scores.*** We performed an overlap analysis to compare gene pairs associated with a significant negative or positive ED score, described above, to the set of 41,773 HAP1 gene pairs that showed a positive genetic interaction and 47,052 gene pairs that exhibited a negative genetic interaction at the standard confidence threshold (|qGI| > 0.3, FDR < 0.1). A small subset of HAP1 query genes (*GFPT1, CDKN2B, ITGAV, SGF29, TAPT1, VPS52, SP1, BCL2, NDUFA2*) each accounted for a disproportionate fraction (~2%) of gene pairs that exhibited significant qGI and ED scores and gene pairs involving this subset of query genes were excluded from this analysis. Among all tested gene pairs (~4 million) in the HAP1 non-redundant genetic interaction dataset (Data File S4), we identified a total of 246,840 gene pairs with significant positive ED score and 218, 484 gene pairs with a significant negative ED score (Data File S21).
A hypergeometric test was used to evaluate the significance of overlap for all possible ED and qGI score combinations (positive ED-negative qGI, positive ED-positive qGI, negative ED-positive qGI, positive ED-positive qGI). In addition, functional co-annotation enrichment analysis was performed using GO biological process, co-complex, and PPI functional standards, and paralog gene lists, described above. The statistical test universe included the set of gene pairs tested for genetic interactions and represented in each functional standard. ED scores for all tested gene pairs are provide in Data File S21 and results pertaining to these analyses are shown in Fig. 8 and fig. S18A-B.

***TCGA Pan-cancer analysis of co-occurring mutations with TP53.*** The cBioPortal for Cancer Genomics repository was used to investigate genetic alterations within the Cancer Genome Atlas (TCGA). A total of 32 distinct cancer types, comprising 10,967 samples (corresponding to 10,953 patients), were subjected to analysis. Our analysis focused on TP53 and 20 genes identified in positive genetic interactions with TP53. Mutation and copy number variant data for those 21 genes were extracted from the cBioPortal database. The co-occurrence of mutations or copy number changes in genes in the positive GI set with TP53 mutations were tested 2 different ways: (1) tracking only co-occurrence of mutations, and (2) tracking co-occurrence of either mutations or copy number. For both approaches, co-occurrence of mutation or copy number deletion in TP53 and each of the candidate genes with positive genetic interactions in HAP1 was tested for enrichment using Fisher's exact test. The Benjamini-Hochberg method was used to correct for the multiple hypothesis testing and calculate false discovery rate (FDR). Results pertaining to these analyses are shown in fig. S18C.

### Comparing genetic interaction and co-essentiality networks

***Jaccard network overlap analysis.*** To compare functional information captured by the HAP1 genetic interaction profile similarity and DepMap co-essentiality networks, we first subsampled the DepMap co-essentiality network (20Q2 release) by randomly sampling two groups of cancer cell lines, with each group consisting of 298 non-overlapping cell lines. We then measured Pearson's correlation coefficients (PCC) based on CERES score essentiality scores for all possible gene pairs in each cell line group. The resultant PCC matrices were binarized at different thresholds (PCC ranging from 0.2 to 0.5 at intervals of 0.01). At each threshold, the Jaccard index between connected genes in the two DepMap networks was computed as follows:

$$J(N1, N2, t) = \frac{E(N1, t) \cup E(N2, t)}{E(N1, t) \cap E(N2, t)}$$

Here, J is the Jaccard index between networks N1 and N2 at threshold t and E is the set of edges of a network N at threshold t.

This process was repeated 10 times using different random resampling of DepMap cancer cell lines. We also computed the Jaccard index between each randomly sampled DepMap co-essentiality network and the HAP1 genetic interaction profile similarity network. The 10 DepMap-DepMap network Jaccard indices and the 20 DepMap-HAP1 genetic profile network Jaccard indices were plotted, where the mean at each threshold is indicated by the solid line and the minimum and maximum at each threshold are indicated with the dotted line. The results of this analysis are shown in Fig. 9A.

***Genetic interaction profile- vs. DepMap co-essentiality-derived network clusters.*** We evaluated clusters derived from the HAP1 genetic interaction profile similarity network for support in the DepMap-derived co-essentiality network. To accomplish this, we applied the hierarchical clustering algorithm described in the section "Hierarchical clustering of HAP1 genetic interaction profiles" to the complete genetic interaction dataset (298 query cell lines x 17, 804 library genes) to generate two layers of "Global" gene clusters (L2 parent clusters and L1 child clusters), which are provided in Data File S9. For each L1 child cluster from this analysis, we computed a cluster statistic reflecting the normalized within-cluster pairwise similarity. This statistic was computed by applying Fisher's z-transformation to the pairwise Pearson correlation coefficient for each within-cluster gene pair. The resulting z-transformed similarity statistics were then averaged per cluster to derive a mean correlation z-score for each cluster. This process was completed for each HAP1 network-derived cluster. Analogous statistics were computed in the DepMap co-essentiality network each of the HAP1 network-derived cluster to derive a z-score reflecting the support for that cluster in the DepMap co-essentiality network. Clusters were then grouped based on the combination of their HAP1 z-score and their DepMap z-score for downstream analyses. Data associated with this analysis is provided in Data File S22 and shown in Fig. 9B-E and fig. S20A-B.

The process described above starts with cluster definitions derived from the HAP1 profile similarity network. We also completed a reciprocal process starting from clusters derived from the same clustering algorithm applied to the DepMap co-essentiality network. The resulting clusters were then evaluated for support in the HAP1 profile similarity network. The figures resulting from this analysis are presented in fig. S20C-E and are also included in Data File S22.

Several other statistics were also computed for each of the clusters derived from the process above. These features include the average SMF for genes in each cluster, the number of publications citing genes in the cluster, degree of annotation for genes in each cluster, enrichment for GO biological processes for each cluster, enrichment for PPI within the cluster, average gene expression level and variance within the cluster, and the DepMap average single mutant phenotype and standard deviation. All cluster statistics related to this analysis are reported in Data File S22. Gene features used for this are also further described in Data File S14. The results of this analysis are provided in Data File S22 and shown in Fig. 9B-E and fig. S20A-E.

***An integrated functional network.*** The complete DepMap co-essentiality profile similarity network was generated by computing Pearson Correlation Coefficient (PCC) between all pairs of gene essentiality profiles derived from analysis of all cell lines tested in the DepMap 20Q2 dataset (depmap.org/portal). NA values were replaced with gene-wise average CERES scores prior to network creation. The HAP1 genetic interaction profile similarity network was constructed from the centroid-corrected genetic interaction dataset, as described above (see section Constructing a genetic interaction profile similarity network).

To construct an integrated network, we normalized the HAP1 GI network using the RPCO method with SNF hyperparameters k=5, $\alpha$=3. Similarly, we applied RPCO with SNF hyperparameters k=5, $\alpha$=5 to normalize the complete DepMap co-essentiality profile network. Normalized networks were then filtered at a similarity threshold corresponding to the 99.85th percentile such that gene pairs with similarity values above this threshold were retained as edges, while gene values below this similarity threshold were removed. Finally, RPCO-normalized and filtered HAP1 genetic interaction profile similarity and DepMap co-essentiality profile similarity networks were integrated using the BIONIC network integration method {Forster, 2022 #5693}. Briefly, BIONIC was run unsupervised for 10,000 epochs with a batch size of 2048, learning rate .0001, and integrated gene embedding size of 4096. The graph attention network (GAT) encoders were set to have an internal embedding size of 256 with 10 attention heads. One GAT layer was used for each input network encoder. All other hyperparameters were unchanged from the previously described default settings {Forster, 2022 #5693}. The resulting matrix contains 19274 genes with embedding dimension 4096.

***Functional evaluation of integrated vs. individual networks.*** The resulting integrated network was functionally evaluated using the FLEX tool, as previously described {Rahman, 2021 #5666}. Briefly, FLEX was used to produce global precision-recall curves based on CORUM complex and GO biological process (BP) functional standards for the integrated network as well as the individual HAP1 genetic interaction profile and DepMap co-essentiality profile networks. Also, FLEX was used to produce per-functional module (defined by specific terms in the GO (BP) and CORUM complex standards) area under the PR curve (AUPRC) scores for each standard. Specifically, for each module in the protein complex or GO BP functional standard, gene pairs with both genes annotated in that module are considered positive examples and gene pairs with only one gene annotated in that module are negative examples in calculating a per-module AUPRC score. A higher per-module score indicates higher pair-wise gene similarities within the module relative to between-module pairs. The results of this analysis are provided in Data File S22 and shown in Fig. 9F-G and fig. S20F-G.

### Estimating size of a complete HAP1 genetic interaction network

***Total number of genetic interactions.*** To estimate the size of a complete genetic interaction network, we considered only the subset of ~11,000 genes that are expressed in in HAP1 cells [log2(TPM+1) > 1](Data File S2). The number of total genetic interactions (i.e. negative + positive interactions) was estimated by multiplying the total number of possible expressed gene pairs (11,000 choose 2 = ~60 500 000 possible gene pairs) by the average genetic interaction density per library gene (File S14). Thus, the estimated size of a complete HAP1 genetic interaction network is ~1.4 million genetic interactions (60 500 000 gene pairs x 0.0226 average interaction density).

***Extreme synthetic lethal and suppressor interactions.*** The current HAP1 genetic interaction network consists of 3, 341 extreme negative, synthetic lethal interactions. Considering the total number of expressed gene pairs tested for genetic interactions (222 query genes x ~11,000 expressed library genes = ~2 442 000 gene pairs), this translates to a synthetic lethal interaction density of ~0.14% (3,341 / ~2 442 000 HAP1 gene pairs x 100%). Thus, we estimate that a complete

HAP1 genetic interaction network will consist of ~85 000 extreme synthetic lethal interactions (i.e. ~60 500 000 expressed gene pairs x 0.0014 interaction density).

The current HAP1 genetic interaction network consists of 1,843 extreme positive, suppression interactions (suppression score > 0.5) where the double mutant fitness is at least 50% greater than the fitness associated with the sickest single mutant of a given gene pair. Considering the total number of expressed gene pairs tested for genetic interactions (222 query genes x ~11,000 expressed library genes = ~2 442 000 gene pairs), this translates to a suppression interaction density of ~0.075% (1,843 / ~2 442 000 HAP1 gene pairs x 100%). Thus, we estimate that a complete HAP1 genetic interaction network will consist of ~45 000 extreme suppression interactions (i.e. 60 500 000 expressed gene pairs x 0.00075 interaction density).

### SUPPLEMENTARY TEXT

#### Genetic interaction profiles reveal mTOR signaling dependencies

As described, genes involved in mTOR signaling were major positive interaction hubs in the HAP1 genetic network (fig. S8E, File S14). Clustering analysis identified two inverse patterns of genetic interactions involving library genes with roles in mTORC1 and mTORC2 signaling (fig. S15B). The first pattern, mTOR Cluster I, comprised 17 query mutant cell lines that showed strong negative interactions with mTORC1 signaling genes and many strong positive interactions with mTORC2 signaling genes (fig. S15B). Many of these query genes (11/17, ~65%), such as *GSK3A* and *GSK3B*, which encode glycogen synthase kinase 3, have established regulatory roles in mTOR signaling (File S14){Hermida, 2017 #6336} or other functional connections to mTOR signaling pathways. mTORC1 activity appeared normal in these cell lines because, in both standard growth conditions and in response to either amino acid starvation or rapamycin treatment, wild type levels of a phosphorylated mTORC1 substrate, p-S6, were observed for most mTOR Cluster I query mutants (fig. S15C-E). Conversely, mTORC2 signaling was severely compromised in these mutants, as indicated by reduced levels of a phosphorylated mTORC2 substrate protein, S473-phosphorylated AKT. Thus, mTOR cluster I positive interactions may indicate that additional perturbation of mTORC2 signaling pathway genes does not result in a more severe fitness defect in query mutant cell lines that already lack mTORC2 activity. However, a subset of mTOR Cluster I positive interactions could be classified as genetic suppression (~19% ,26/139, $P < 4.1x10^{-10}$, hypergeometric)(File S17), identifying genes whose LOF may improve the fitness of cells with reduced mTORC2 activity. Nonetheless, these findings suggest that the fitness of mTOR Cluster I query mutants are more dependent on mTORC1 signaling relative to mTORC2 signaling.

A second cluster, mTOR Cluster II, highlighted 68 different query mutant cell lines that had negative interactions with mTORC2 signaling genes and positive interactions with mTORC1 signaling genes, suggesting that the fitness of these query mutants depends on mTORC2 signaling. While some of these query genes (23/68, ~34%), like the *LATS1* gene that encodes a core component of the HIPPO signaling pathway, are known to impact mTOR signaling {Gan, 2020 #6337}, most mTOR Cluster II query genes did not appear to have a direct role in mTOR signaling.

An inverse correlation between the relative levels of mTORC1 and mTORC2 activity has been observed previously across different human cell lines, suggesting that the balancing of mTORC1 and mTORC2 activity is not specific to HAP1 cells {Sarbassov, 2004 #5810}. Indeed, differences in mTORC1 and mTORC2 dependencies were also reflected in DepMap LOF genetic screens. Like HAP1 query gene mutants in mTOR cluster I, 40 DepMap cancer cell lines showed a strong fitness dependency on mTORC1 signaling genes while perturbation of mTORC2 signaling genes led to improved fitness in the same cell lines. Like HAP1 query gene mutants in mTOR cluster II, we identified 15 cancer cell lines that depended on mTORC2 signaling genes for normal fitness but showed increased fitness following inactivation of mTORC1 genes (File S14).

### SUPPLEMENTARY FIGURES

figure S1

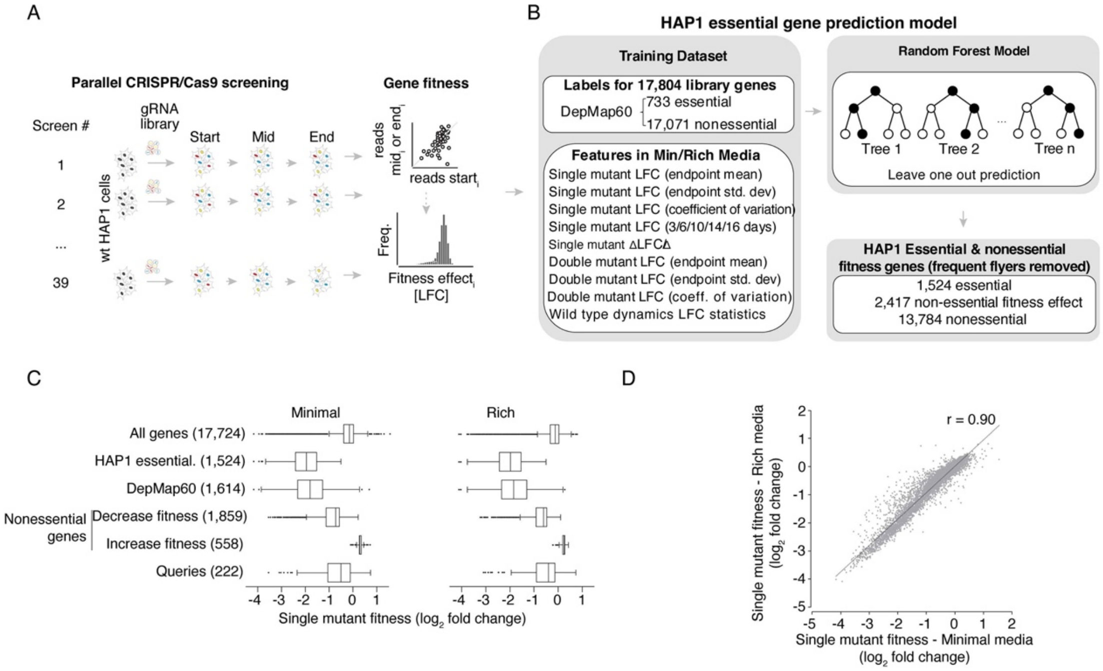

**Fig. S1. Identifying essential genes and measuring single mutant fitness in HAP1 cells. (A)** Experimental setup for performing parallel, genome-scale pooled CRISPR screens in parental HAP1 cells with the TKOv3 library to measure single mutant fitness (LFC = log fold change). **(B)** Workflow for generating a random forest model to identify essential genes based on HAP1 screen data. **(C)** Box plots showing the distribution of single mutant fitness effects for HAP1 genes grouped into the indicated fitness categories. DepMap60 includes genes that were classified as essential (CERES < -0.5) in at least 60% of cell lines tested in the DepMap 20Q2 dataset. The number of genes in each category is indicated in brackets. **(D)** Scatter plot of single mutant fitness effects (mean Log Fold Changes) derived from parental HAP1 screens in rich (n=18) or minimal (n=21) media.

figure S2

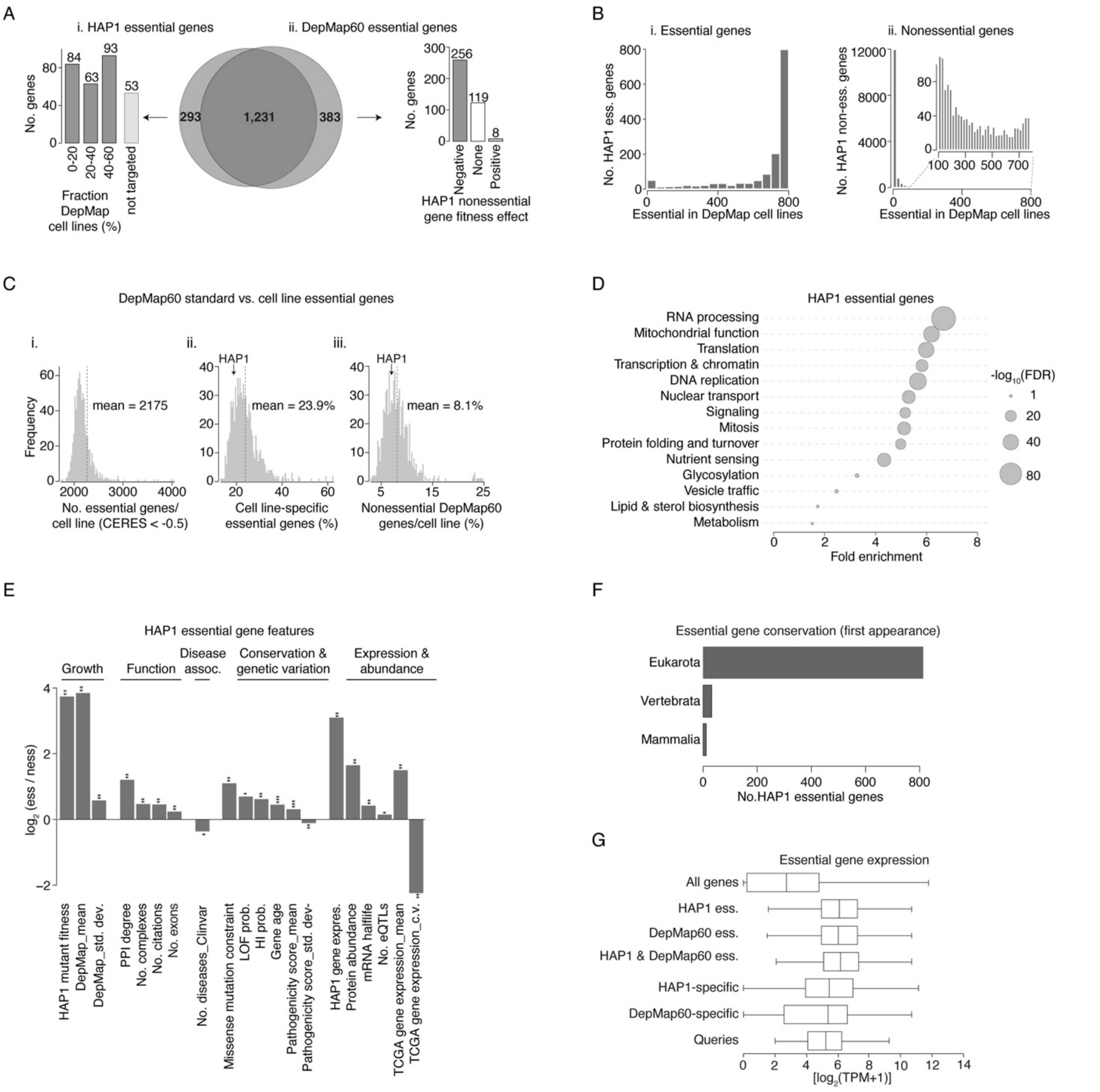

**Fig. S2. Characterization of HAP1 essential genes. (A)** Venn diagram showing the overlap between HAP1 essential genes and DepMap60 genes. (i) Bar plot reporting the number of HAP1-specific essential genes (293 genes) that show DepMap essential phenotypes (mean CERES < -0.5) in the indicated fraction of DepMap cell lines. Genes not targeted by DepMap CRISPR libraries are indicated. (ii) Bar plot reporting the number of DepMap60-specific essential genes (383 genes) exhibiting fitness effects in HAP1 cells. **(B)** Distributions showing the number of HAP1 essential genes (i) and non-essential genes (ii) that exhibited a similar phenotype in DepMap cell lines (number of DepMap cell lines for which CERES < -0.5). **(C)** (i) Distribution illustrating the percentage of essential genes (CERES <-0.5) per DepMap cell line. Dotted line indicates the average number of essential genes per cancer cell line in DepMap. (ii) Distribution showing the percent of DepMap cell line-specific essential genes per cell line. Dotted vertical line indicates the average percentage of cell line-specific essential genes identified in a given DepMap cancer cell line. The percentage of HAP1-specific essentials is indicated. (iii) Distribution illustrates the percentage of DepMap60 genes that are nonessential (CERES > -0.5) in a particular cancer cell line. The average percentage of DepMap60 genes that are non-essential for viability in a specific cancer cell line and the percentage of DepMap60 genes nonessential in HAP1 cells are indicated (dotted lines). **(D)** Reactome Pathway terms, ranging in size from 10 to 300 genes {Griss, 2020 #6281}, statistically enriched (hypergeometric test, Benjamini-Hochberg-corrected FDR < 0.2) among HAP1 essential genes are summarized according to the functional descriptions shown. **(E)** Sequence, functional and evolutionary properties significantly associated with HAP1 essential genes relative to HAP1 nonessential genes (ess/noness). * indicates level of statistical significance (* $P < 10^{-3}$, ** $P < 10^{-10}$, Wilcoxon rank-sum). **(F)** Box plot showing the conservation of HAP1 essential genes across the indicated evolutionary classes. **(G)** Box plot indicating average gene expression values ($log_2$[TPM+1]) for the indicated gene sets broken down by fitness categories.

figure S3

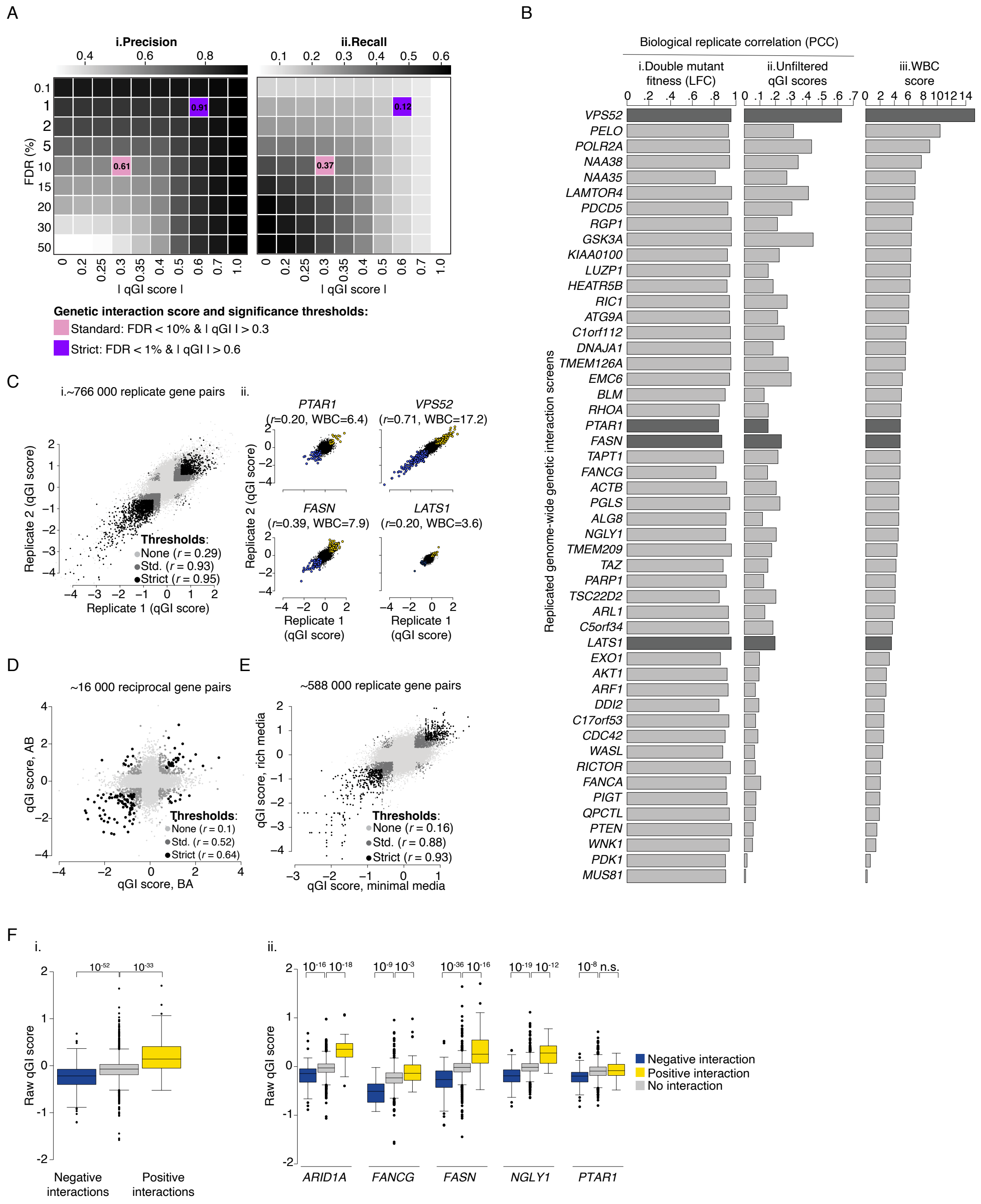

**Fig. S3. Data quality and reproducibility. (A)** Heatmap illustrating precision and recall estimates derived using an MCMC approach based on a set of screens with at least 4 replicates each. Estimated (i) precision and (ii) recall are plotted at varying effect size (qGI score) and FDR cutoffs, as described {material, #6370}. Standard (pink) and strict (purple) qGI score and FDR thresholds used to filter the raw genetic interaction dataset are shown. **(B)** Biological reproducibility analysis from screens performed for the indicated query genes (range n = 2-5). Bar plots showing the average PCC for (i) LFCs/double mutant fitness, (ii) unfiltered qGI scores, and (iii) Within-Between Correlation or

WBC scores {Billmann, 2023 #5721}. Dark grey bars indicated query genes that are highlighted in Panel C. **(C)** (i) Scatter plot of qGI scores for all replicated gene pairs tested in this study. PCCs were computed after applying the indicated confidence thresholds. (ii) Scatter plots of qGI scores derived from replicated screens with the indicated query gene mutants. **(D)** Scatter plot of qGI scores between reciprocally tested gene pairs. PCCs were computed after applying the indicated confidence thresholds. **(E)** Scatter plot of qGI scores for the same gene pairs measured in different media conditions. PCCs were computed after applying the indicated confidence thresholds. **(F)** Box plots showing results of re-screening 5 query genes from our genome-wide dataset using an independent CRISPR-KO library that targeted ~1,200 genes with ~30 gRNAs/gene (i.e. ~37,000 gRNAs in total)(File S6){material, #6370}. These gRNAs were not present in the TKOv3 library but targeted genes that showed significant genetic interactions with at least one of the 5 selected query genes in the TKOv3 library. (i) Box plot shows the distribution of all unfiltered qGI genetic interaction scores derived from screens using an independent gRNA library {material, #6370}, for groups of gene pairs that showed a negative (blue), positive (yellow) or no interaction (grey) in screens using the TKOv3 gRNA library. (ii) Box plot showing the distribution of qGI genetic interaction scores derived from screens using an independent gRNA library and involving specific query genes that showed a negative (blue), positive (yellow) or no interaction (grey) in screens using the TKOv3 gRNA library (right). This analysis recapitulated both negative and positive interactions, indicating that the genetic interactions we identified were not driven by gRNA-specific phenotypes.

figure S4

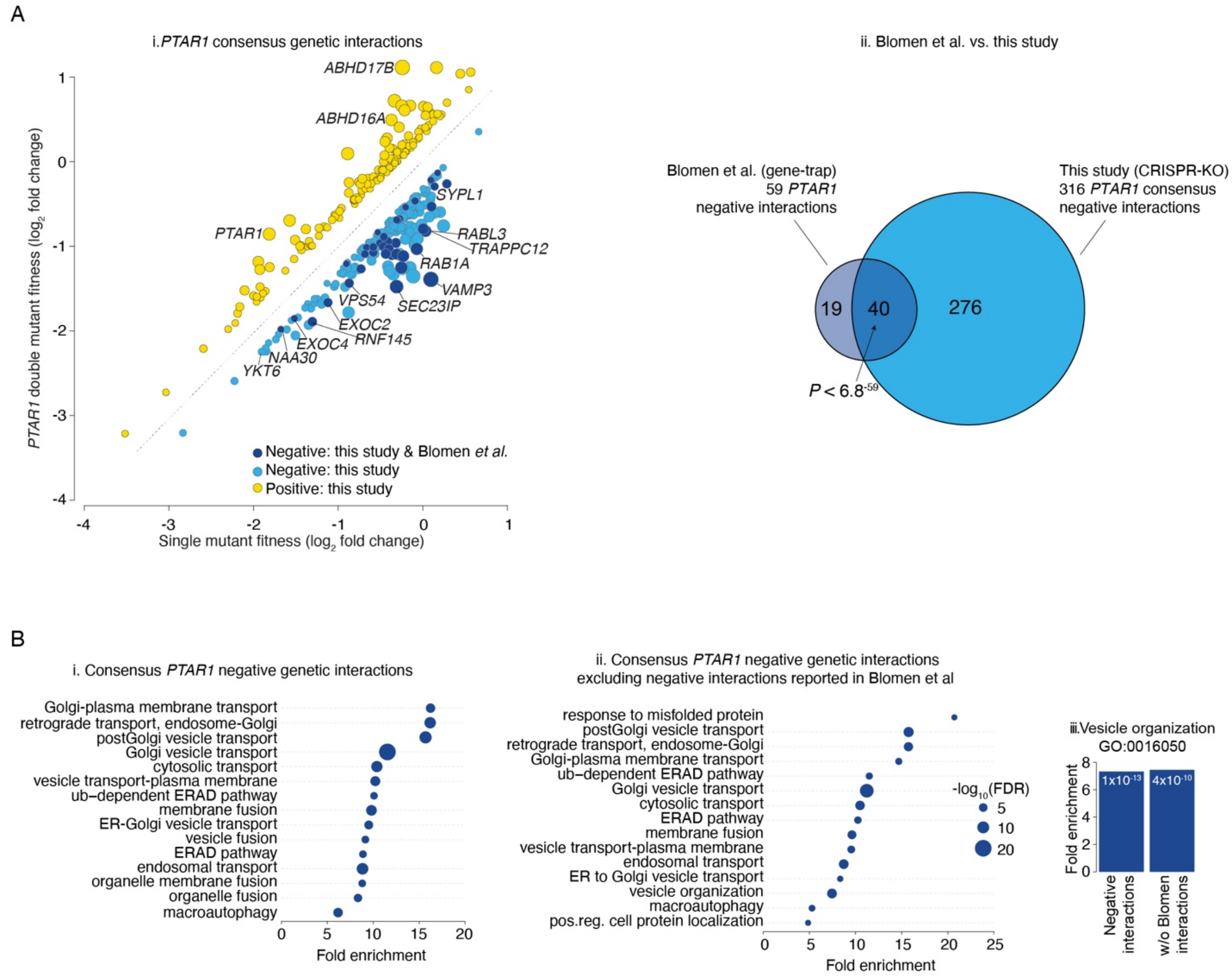

**Fig. S4. *PTAR1* genetic interactions identified in a CRISPR-KO versus a gene trap approach. (A)** (i) Scatterplot of mean consensus qGI scores derived from 4 independent *PTAR1* query screens with TKOv3 (GIN003, GIN043, GIN044, GIN109). Negative (blue) and positive (yellow) genetic interactions satifying a standard confidence threshold (|qGI| > 0.3 and FDR < 0.1) are shown. Selected negative (blue) and positive (yellow) interactions are indicated. *PTAR1* negative interactions identified by the gene-trap method {Blomen, 2015 #4298} are shown in dark blue. (ii) Venn diagram illustrating the overlap between *PTAR1* negative interactions identified in this study and the gene trap analysis (Blomen *et al*.){Blomen, 2015 #4298}. **(B)** (i) Functional enrichment (hypergeometric test, Benjamini-Hochberg-corrected FDR < 0.2) of genes that showed a negative interaction with *PTAR1* in this study or (ii) genes that showed a negative interaction with *PTAR1* after excluding genes that also showed a negative interaction with *PTAR1* in the Blomen *et al*. gene trap study {Blomen, 2015 #4298}. (iii) Bar chart indicating fold-enrichment for each of the indicated gene sets, specifically for vesicle organization.

figure S5

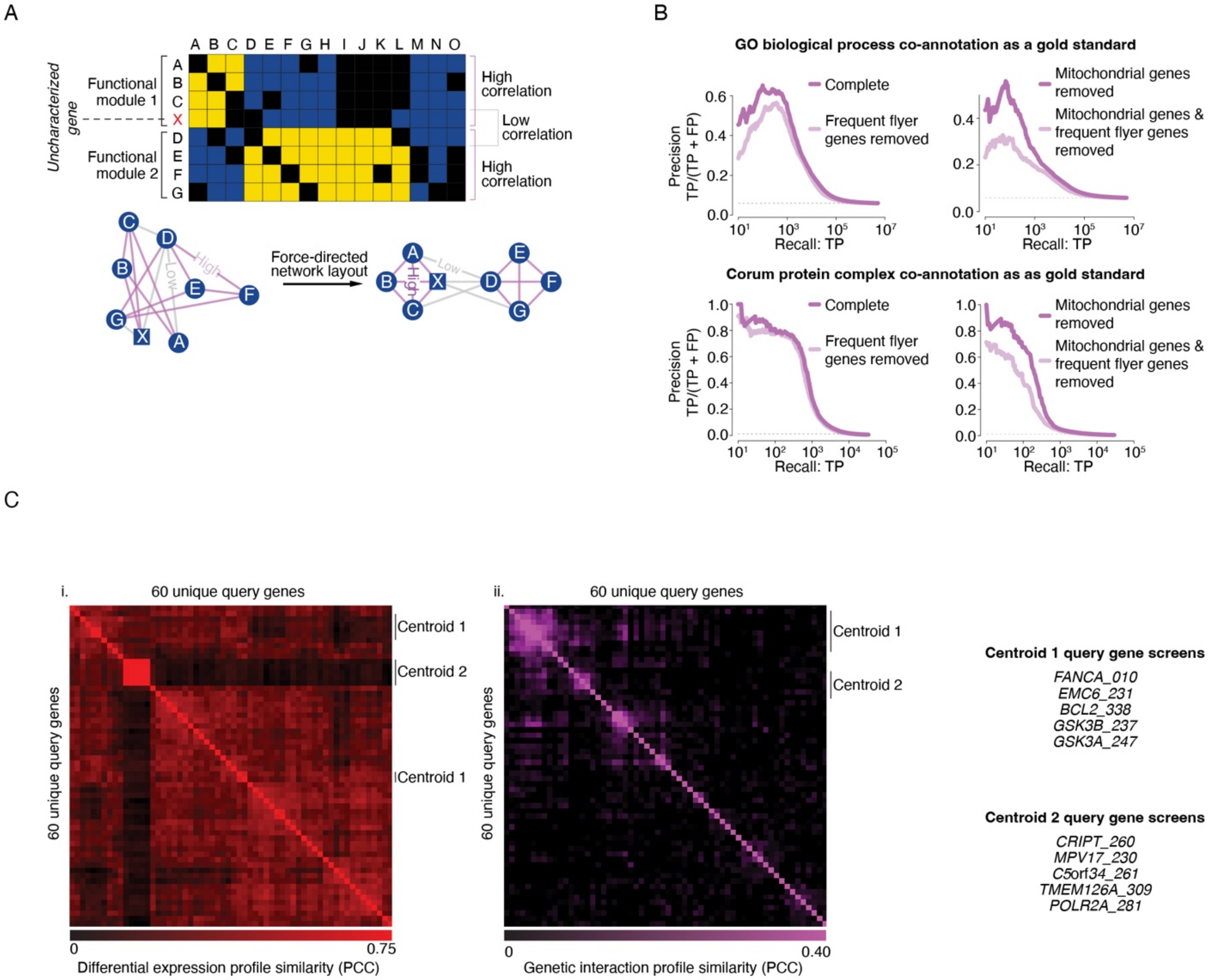

**Fig. S5. Functional evaluation of genetic interaction profiles. (A)** Schematic illustrating genetic interaction profile similarity. The set of negative (blue) and positive (yellow) interactions for a given mutant is referred to as a genetic interaction profile. Two-dimensional hierarchical clustering groups genes together based on genetic interaction profile similarity enabling identification of highly correlated groups of genes that correspond to functionally related gene modules. Genetic interactions can also be visualized as a correlation-based network connecting genes with similar genetic interaction profiles. Using a force-directed network layout, genes with highly similar genetic interaction profiles (purple lines) are placed close to each other in the network while genes with less similar interaction profiles (grey lines) are placed further apart from one another {Shannon, 2003 #1200}. **(B)** Genes with varying degrees of genetic interaction profile similarity were evaluated for overlap with either GO biological process co-annotations or CORUM Protein complex co-annotation using precision-recall analysis {material, #6370}. Gene pairs were sorted based on Pearson correlation coefficients, reflecting similarity in their genetic interaction profiles. Grey dashed lines show the background rate of co-annotation for the relevant set of gene pairs. Precision-recall analysis was completed separately for genetic interaction profiles derived from all library genes in the dataset as well as genetic interaction profiles excluding genes associated with highly variable single mutant fitness and/or mitochondrial genes, as indicated {material, #6370}. **(C)** (i) Matrix of 60 unique query gene mutant cell lines clustered based on pair-wise similarity of their differential gene expression profiles (red) or (ii) of their genetic interactions (purple). Two groups of genes that cluster together in both matrices and whose genetic interaction profiles were used to correct the complete genetic interaction matrix in order to construct a genetic interaction profile similarity network are indicated as Centroid 1 and Centroid 2 and described in detail in the methods section {material, #6370}. The query gene mutant cell lines that comprise Centroid 1 and Centroid 2 are listed.

figure S6

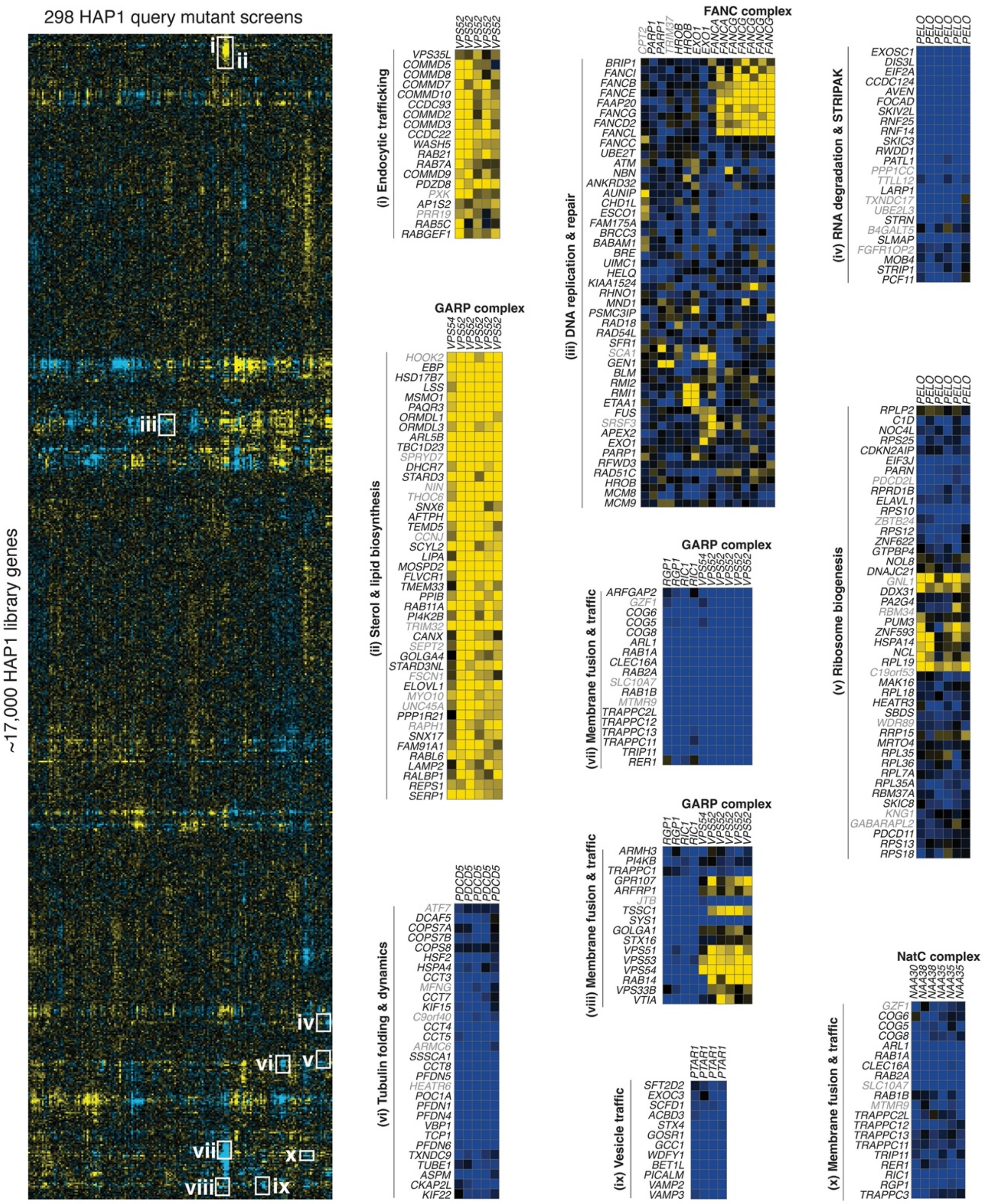

**Fig. S6. Two-dimensional hierarchical clustering of the HAP1 genetic interaction dataset.** Negative (blue) and positive (yellow) genetic interactions are shown. Rows in the matrix correspond to 17,297 genes in the TKOv3 library. Columns in the matrix represent 298 genome-scale genetic interaction screens corresponding to 222 unique query mutant cell lines. Sections (white outlines) are expanded to allow visualization of specific query-library gene-gene interactions.

figures S7

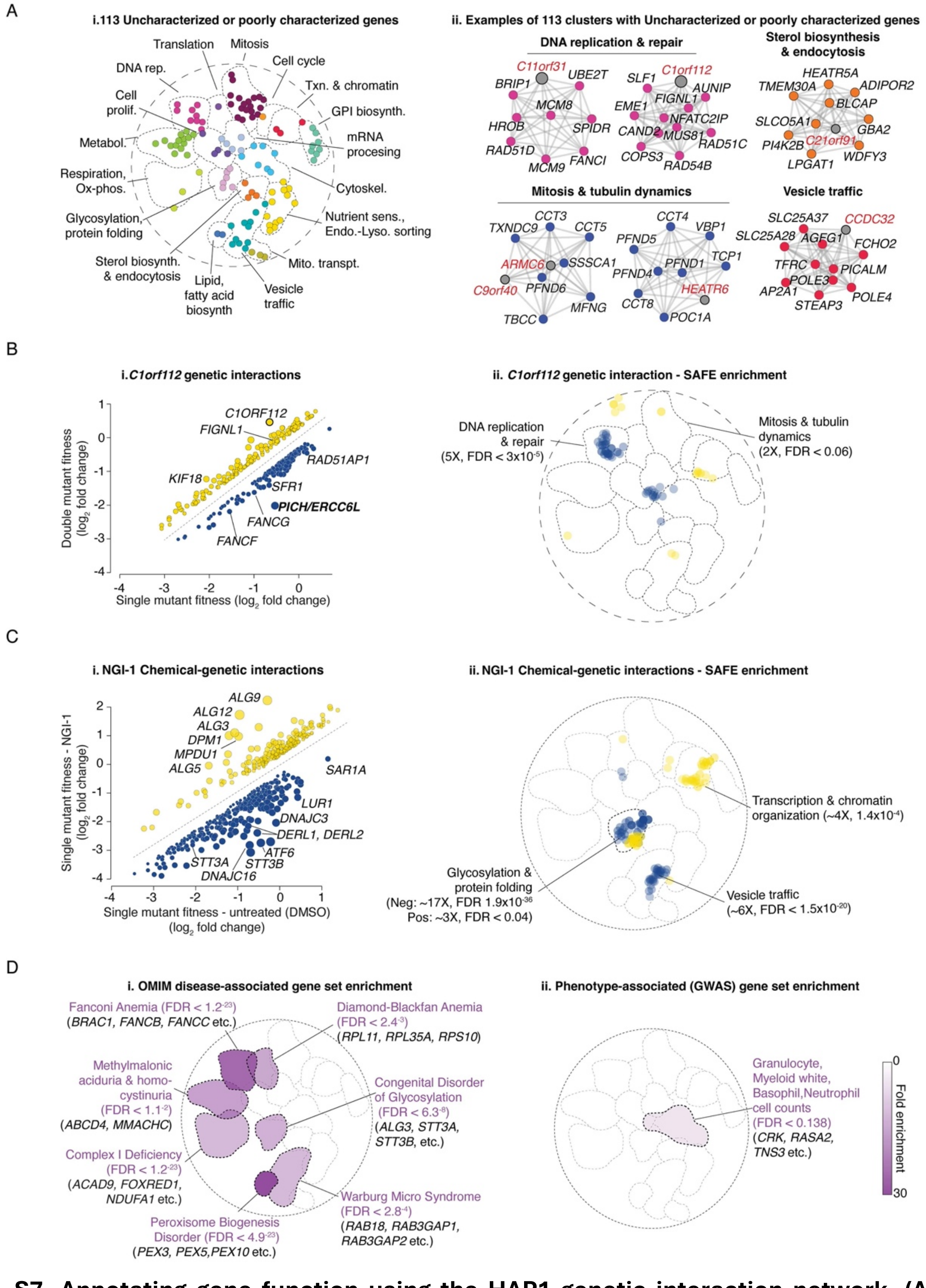

**Fig. S7. Annotating gene function using the HAP1 genetic interaction network. (A)** (i) Poorly characterized genes (i.e. GO bioprocess terms + PubMed citations < 15) that localize in a specific bioprocess–enriched cluster on the genetic interaction network shown in Fig. 2A-B. Nodes are colored according to bioprocesses shown in Fig. 2B. (ii) Selected examples of re-clustered gene modules from the complete HAP1 genetic interaction matrix, where genes with the same node color have a shared function and poorly characterized genes are indicated with grey nodes and red labels. **(B)** (i) Scatterplot of qGI scores for a HAP1 *C1*orf*112/FIRRMM* mutant query screen. Negative (blue) and positive (yellow) genetic interactions that satisfied a standard confidence threshold (|qGI| > 0.3 and FDR < 0.1) are shown, and specific negative and positive interactions are labeled, including the strongest negative interaction that confirmed a previously identified *PICH/ERCC6L-C1*orf*112/FIRRMM* interaction (bold font) {Stok, 2023 #5774}. (ii) Regions of the HAP1 genetic interaction network that are significantly enriched (hypergeometric test, Benjamini-Hochberg-corrected, FDR < 0.001) for genes exhibiting negative (blue) or positive (yellow) genetic interactions with a *C1*orf*112/FIRRMM* query mutant cell line. Enrichment was calculated using SAFE as described in the methods {material,  #6370}. **(C)** (i) Scatterplot of chemical-genetic interactions identified in the presence of 15uM NGI-1. Negative (blue) and positive (yellow) chemical-genetic interactions that satisfied a standard confidence threshold (|score| > 0.3, FDR < 0.1) are shown and select genes that exhibited negative and positive chemical-genetic interactions with NGI-1 are indicated. (ii) Regions of the HAP1 profile similarity network that are significantly enriched for genes exhibiting negative (blue) or positive (yellow) chemical-genetic interactions with NGI-1 (right). Enrichments for NGI-1 negative or positive chemical-genetic interactions within functional domains in the HAP1 genetic profile similarity network were determined using a hypergeometric test for over-enrichment was conducted for each domain, followed by a Benjamini–Hochberg correction. Fold enrichment and significance of negative and positive chemical-genetic interaction enrichment is indicated. **(D)** Regions of the HAP1 genetic interaction network that are significantly enriched for (i) genes associated with a Mendelian-inherited disease (OMIM disease) or (ii) phenotype-associated GWAS. For each disease or trait gene set, a hypergeometric test for enrichment was conducted against the 17 bioprocess domains defined in the HAP1 profile similarity network, followed by a Benjamini–Hochberg correction.

figure S8

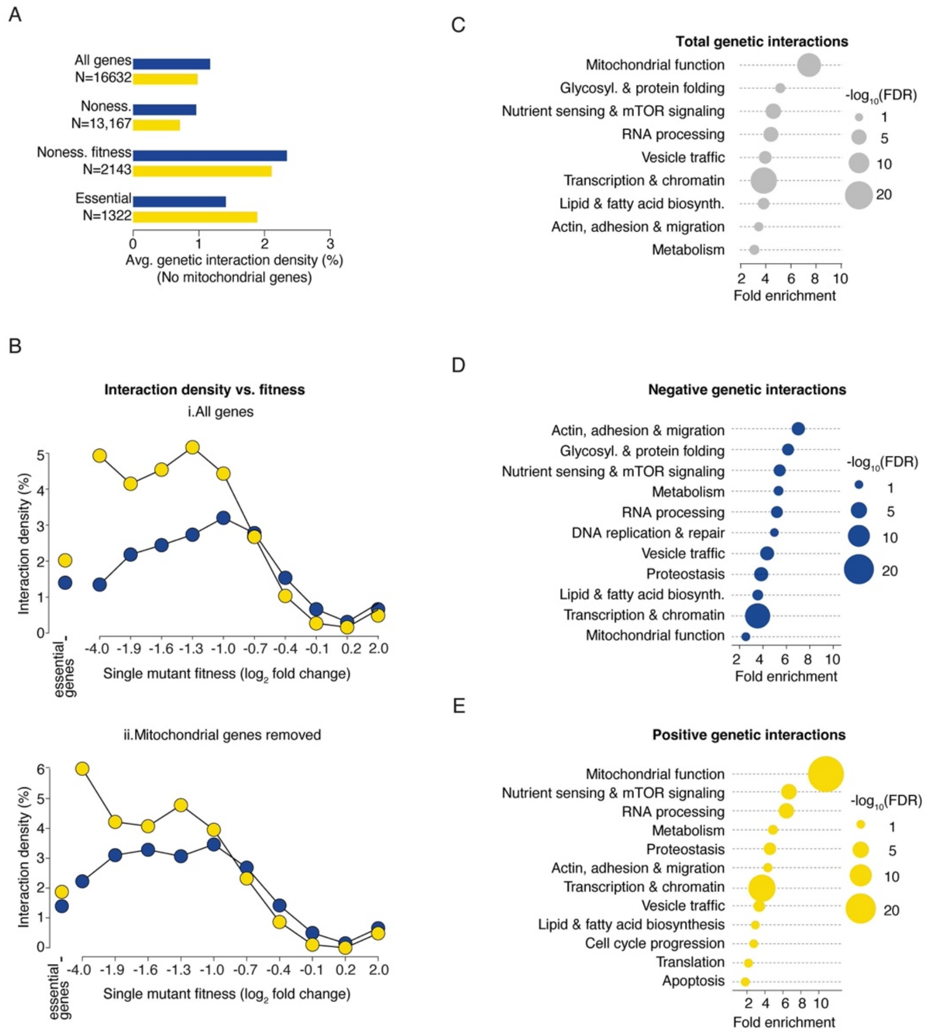

**Fig. S8. Genetic interaction density analysis. (A)** Bar chart showing genetic interaction density (observed interactions/total gene pairs screened) for library genes (excluding genes with roles in mitochondrial function) by category (all genes, nonessential (noness), nonessential with fitness phenotypes (noness fitness), essential) at |qGI| >0.3, FDR < 0.1. Negative (blue), positive (yellow) and total (grey) interaction densities, with the number of genes in each category indicated. **(B)** Line plots showing average density of negative (blue) and positive (yellow) genetic interactions for (i) all library genes and (ii) excluding genes with roles in mitochondrial function relative to library gene single mutant fitness. The average negative and positive interaction density for essential genes (ess.) is also shown on the left. **(C-E)** Dot plots showing functional enrichment of genetic interaction hubs. Reactome Pathway terms {Griss, 2020 #6281} statistically enriched (hypergeometric test, Benjamini-Hochberg-corrected FDR < 0.2) among highly connected genes representing total (grey), negative (blue) and positive (yellow) genetic interaction hub genes were summarized according to the functional descriptions shown.

figure S9

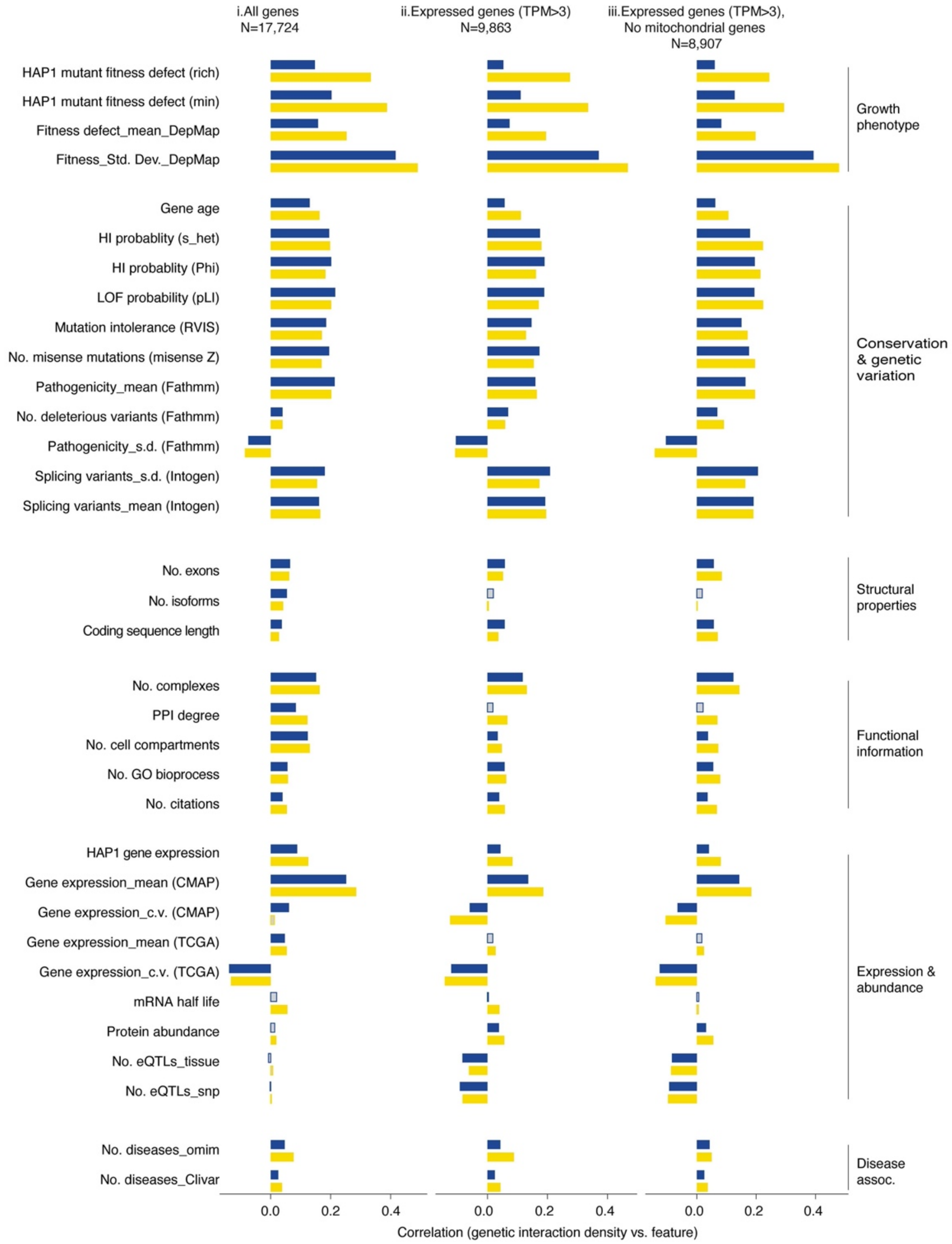

**Fig. S9. Correlation analysis of genetic interaction density.** Negative (blue) and positive (yellow) genetic interaction density was calculated for (i) all genes targeted by the TKOv3 CRISPR library, (ii) genes expressed in HAP1 cells and (iii)genes expressed in HAP1 cells without mitochondrial genes. The standard confidence threshold (|qGI| ≥ 0.3, FDR < 0.1) was applied and interaction density was computed as the percentage of observed interactions. Pearson's correlation coefficient was used to measure associations between genetic interaction density and features that are continuous or count based. Significant ($P$ < 0.05, using the cor.test function in R and Pearson's correlation) non-zero correlations relationships with negative (blue) and positive (yellow) interaction density. Open bars indicate relationships for which the 95% confidence interval on the correlation coefficient includes 0. Given that analysis of different features required different statistical tests, and some features are not expected to be independent of each other, multiple hypotheses correction procedures were not applied (See Methods and File S14 for details). HI=haploinsufficiency; LOF=loss of function.

figure S10

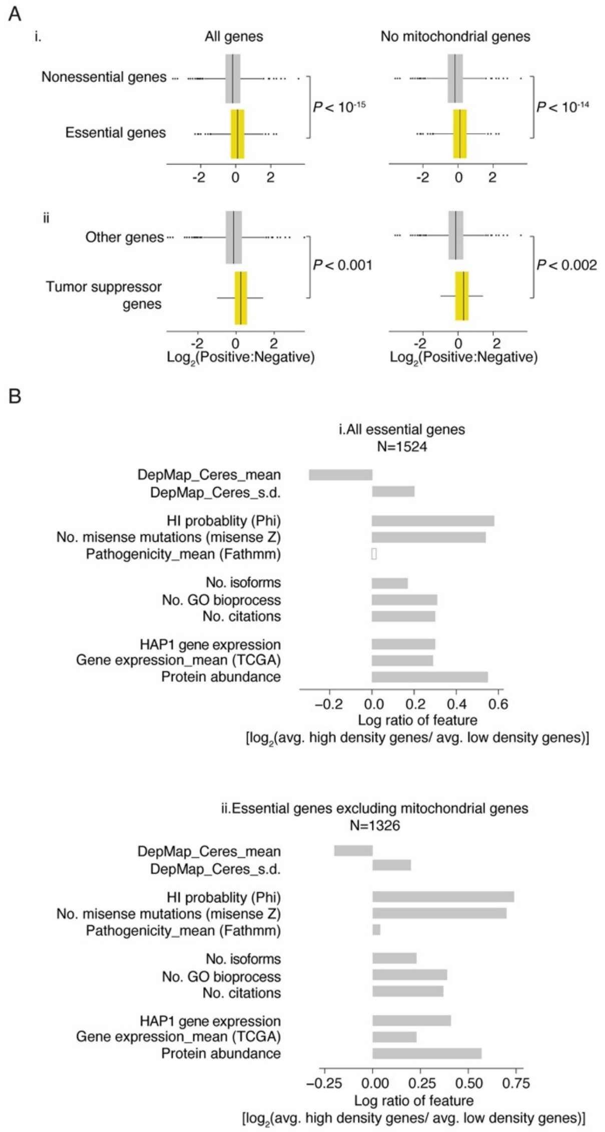

**Fig. S10. Properties of essential gene genetic interactions. (A)** (i) Box plots illustrating the distribution of the ratio of positive to negative genetic interactions for all non-essential and all essential library genes with (all genes) or without (no mitochondrial genes) mitochondrial genes. (ii) Box plots illustrating the distribution of the ratio of positive to negative genetic interactions for tumor suppressor genes and all other genes with (all genes) or without (no mitochondrial genes) mitochondrial genes. See File S19 for gene lists. **(B)** Sequence, functional and evolutionary properties that are significantly associated with essential library genes that exhibited high genetic interaction density relative to essential genes with few genetic interactions. The analysis was performed using (i) all genes or (ii) excluding mitochondrial genes. High interaction density was defined as the top 20% of HAP1 essential genes with the highest total genetic interaction density whereas low interaction density consisted of the 50% of HAP1 essential genes with the lowest genetic interaction density. For each feature, the mean of the high-density group and the low-density group was computed, and the $\log_2$ ratio of these means is plotted. Open bars indicate relationships for which the 95% confidence interval on the correlation coefficient includes 0.

figure S11

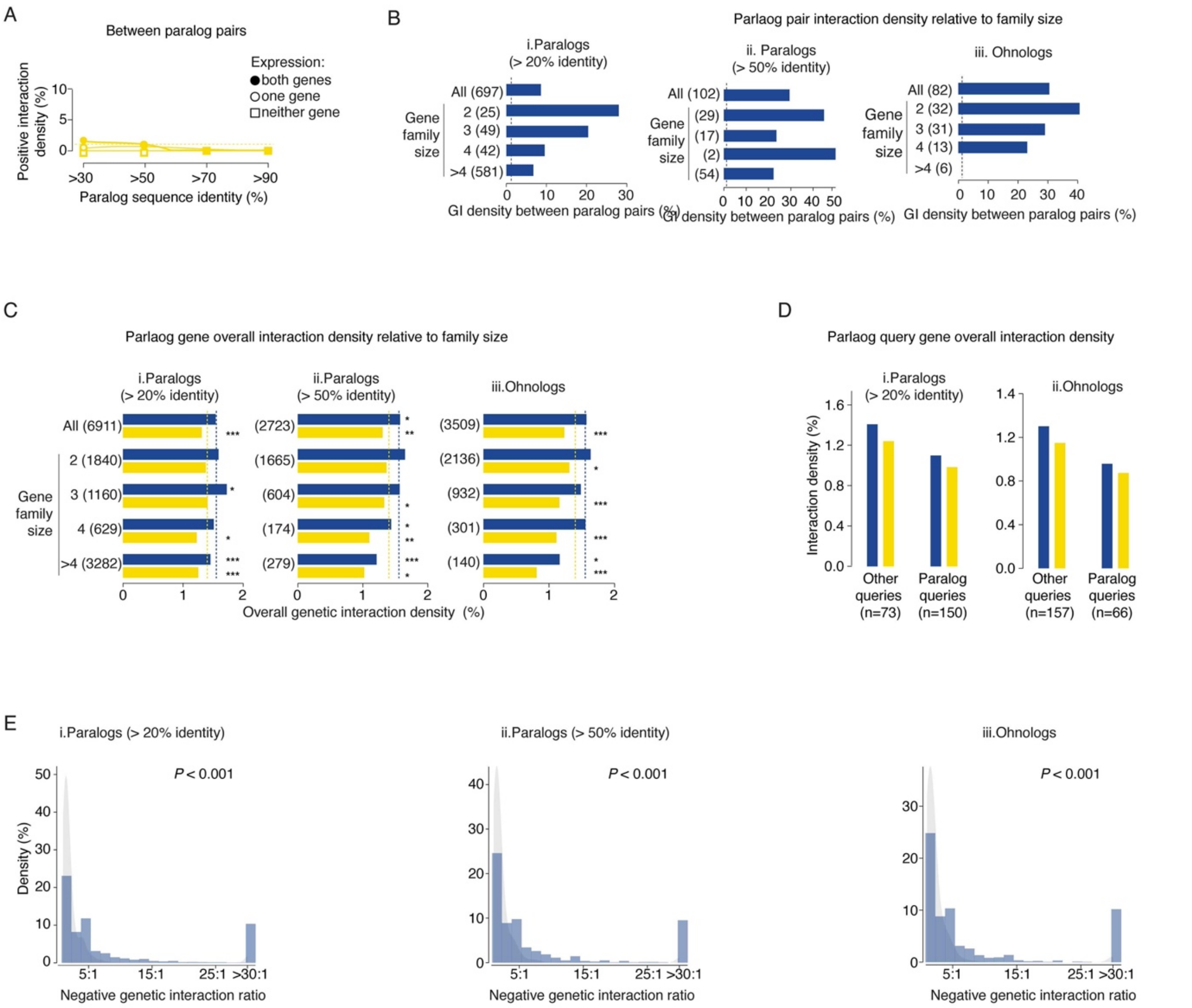

**Fig. S11. Genetic interaction density associated with paralogs.** **(A)** Positive genetic interaction density among pairs of duplicated genes with increasing sequence identity (i.e. paralogs). **(B)** Bar plots depicting paralog pair genetic interaction density (|qGI| >0.3, FDR < 0.1) relative to family size for paralogs sharing (i) >20% or (ii) >50% sequence identity, as well as (iii) ohnologs {Singh, 2020 #6282}. Interaction density was expressed as a percentage of all tested paralog gene pairs. The numbers of paralog pairs tested in each family size bin are indicated in brackets. **(C)** Bar plots depicting the average negative (blue) and positive (yellow) genetic interaction densities for individual paralog library genes with all tested query genes relative to gene family size for paralogs sharing (i) >20% or (ii) >50% sequence identity, as well as (iii) ohnologs. The numbers of interactions involving a paralog gene tested in each family size bin are indicated (* indicates level of statistical significance, * $P$ < 0.05, ** $P$ < 0.01, *** $P$ < 0.001, Wilcoxon rank-sum). **(D)** Bar plots indicating the average negative (blue) and positive (yellow) genetic interaction densities for individual paralog query genes versus non-duplicated query genes surveyed in this study. Query gene paralogs were defined as genes that share (i) at least 20% sequence identity, as well as (ii) ohnologs. **(E)** Histograms of negative interaction degree ratio as evidence for asymmetric functional divergence for paralog gene pairs sharing (i) >20% or (ii) >50% sequence identity, and (iii) ohnologs. The ratio is defined as the number of unique negative interactions (degree) identified for each gene of a duplicate pair with the higher degree in the numerator. Shown for comparison is another degree ratio histogram (symmetric null model) in which interactions for every duplicate pair are redistributed to either member with equal probability (grey).

figure S12

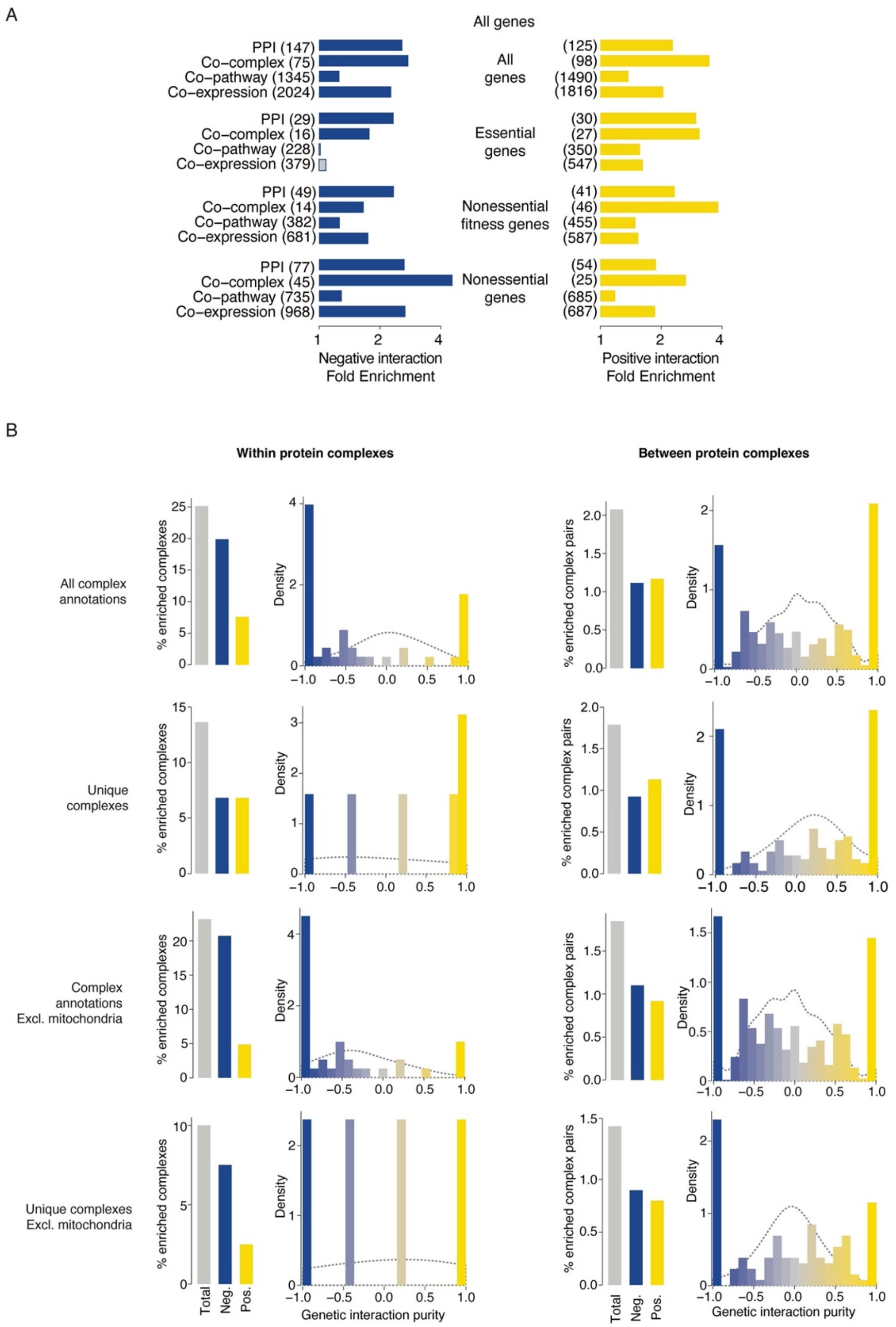

**Fig. S12. Relating genetic and physical interactions. (A)** Bar charts show significant fold-enrichment ($P$ < 0.05 hypergeometric test) for all gene pairs (including mitochondrial genes) encoding physically interacting proteins (PPI), co-complex proteins, co-pathway proteins, or co-expressed gene pairs among negative (blue) and positive (yellow) genetic interactions. Enrichment was measured for all gene pairs, essential gene pairs, nonessential genes with fitness phenotypes, and nonessential genes lacking a fitness phenotype. Grey bars indicate non-significant enrichment. The numbers of gene pairs tested for enrichment are indicated in brackets. **(B)** (left panels) Bar graphs summarize complexes enriched for genetic interactions among members of the same complex (Within protein complex) or enrichment of genetic interactions between pairs of protein complex (Between protein complex) as the percentage of protein complexes (CORUM database) with enriched genetic interactions, categorized as all interactions (grey), negative (blue), or positive (yellow) within/between protein complexes. (right panels) Histograms illustrate purity scores indicating the proportion of negative and positive interactions within a protein complex or between a pair of protein complexes. Purity scores range from -1 (purely negative interactions) to 1 (purely positive interactions) among complex members. The dotted grey line indicates the random expectation based on a binomial distribution. Analyses include all CORUM annotated complexes or a subset of complex annotations where redundant complexes with overlapping genes were removed. In both cases, analyses were based on complexes and complex pairs with >5 tested gene pairs. Analyses were repeated after excluding genes with mitochondrial-related functions.

figure S13

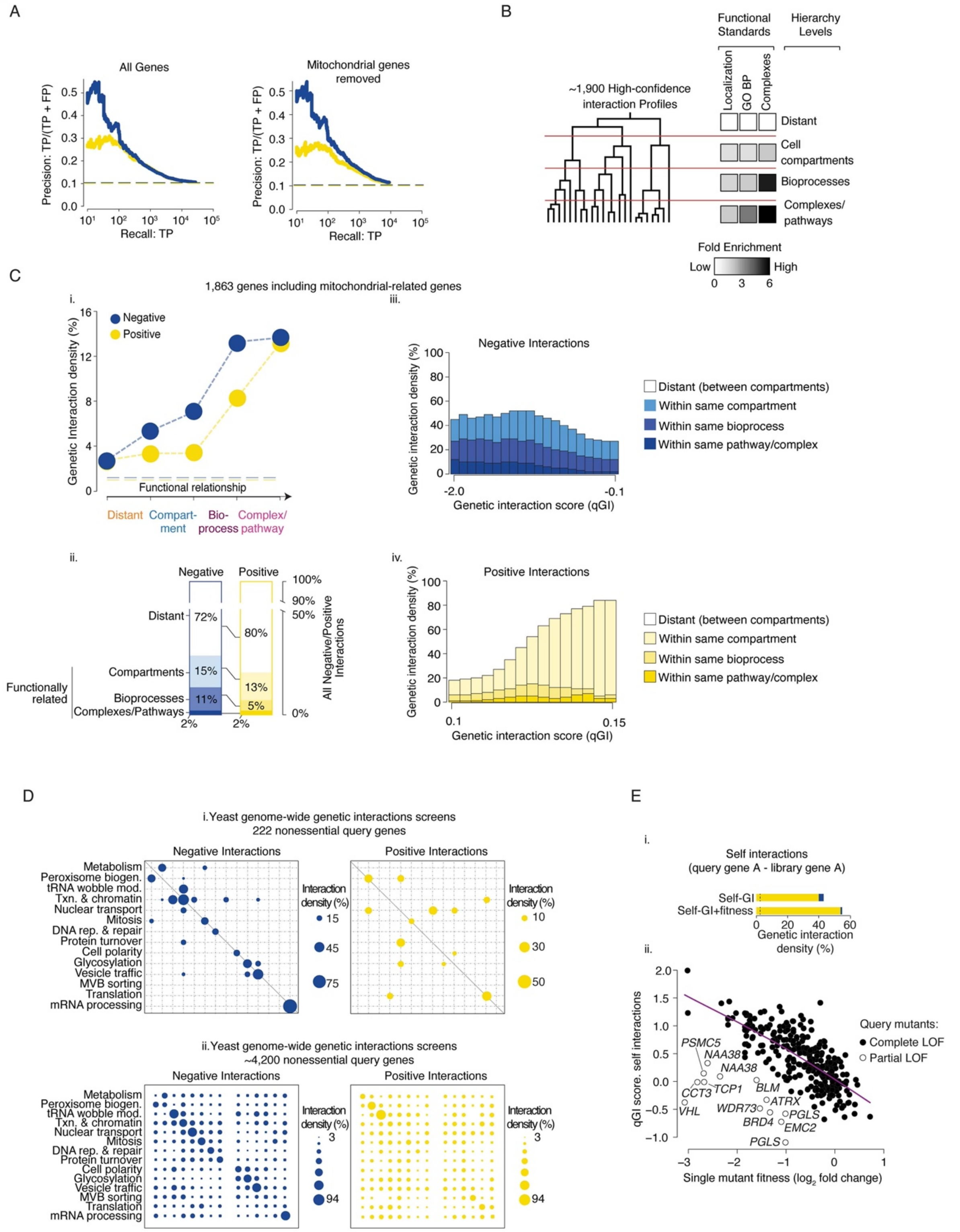

**Fig. S13. Functional distribution of genetic interactions. (A)** Precision-recall plots for negative (blue) and positive (yellow) genetic interactions (|qGI| >0.3, FDR < 0.1) based on co-annotation to GO biological process terms, including and excluding gene with mitochondrial-related functions. Dashed lines indicate background co-annotation rates. **(B)** Genetic interaction profile–derived hierarchy schematic. Genes with highly correlated genetic interaction profiles form small, densely connected clusters representing specific pathways or protein complexes. Intermediate correlation combines sibling clusters into larger biological process–enriched clusters, which further combine into larger clusters corresponding to cell compartments. Grey scale cale bar indicates sibling cluster enrichment for functional terms. Analysis includes ~1600 genes with high confidence profiles, excluding genes with mitochondrial-related functions. **(C)** (i) Line graph depicts density of negative (blue) and positive (yellow) genetic interactions (|qGI| > 0.3, FDR < 0.1) within a specific level of the genetic network hierarchy. Horizontal dashed lines represent background interaction density. (ii) Stacked bar chart shows functional distribution of all negative (blue) and all positive (yellow) interactions (|qGI| > 0.3, FDR< 0.1) among genes in the genetic network hierarchy, depicting percentages within clusters corresponding to a cellular compartments, bioprocesses, or complexes/pathways. The combined fraction of functionally related interactions is indicated (*). (iii-iv) Bar graphs show the fraction of negative (iii, blue) or positive (iv, yellow) interactions connecting genes within the same cluster at varying functional relatedness levels. **(D)** (i) Dot plots depicting yeast network density of negative (blue) and positive (yellow) nonessential gene interactions (|SGA score| >0.08, P < 0.05) within and across biological processes for 222 randomly sampled yeast query mutants. Node size reflects the fraction of interacting gene pairs observed for a given pair of biological processes; node color indicates significance above random expectation. (ii) Dot plot shows a similar analysis based on the complete set of ~4200 yeast nonessential gene query mutant strains. Nodes on the diagonal represent interactions within the same biological process; off-diagonal nodes represent interactions between processes. **(E)** (i) Bar plot shows the overall genetic density of negative (blue) and positive (yellow) interactions for all self-genetic interacting genes and the subset of self-genetic interacting genes associated with a fitness phenotypes. (ii) Scatter plot illustrating the correlation between HAP1 single mutant fitness and genetic interaction score (qGI) for genes with self-genetic interactions.

figure S14

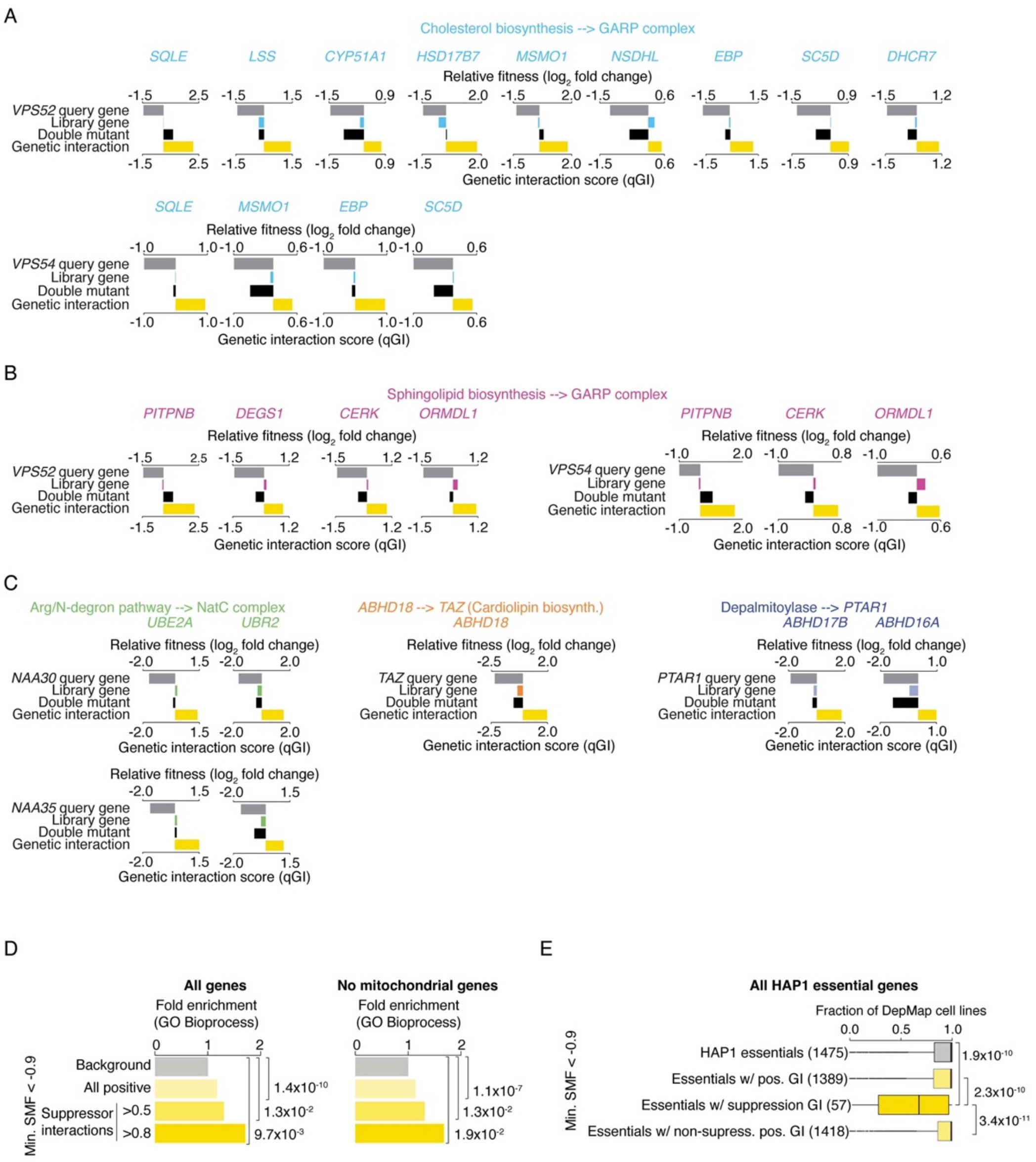

**Figure S14. Genetic suppression interactions. (A-C)** Bar graphs illustrating quantitative analysis of specific genetic suppression interactions. Query gene single mutant fitness (grey bars), library gene single mutant fitness (gene-specific colored bars), double mutant fitness (black bars) and positive genetic interaction score (qGI, yellow bars) are shown. **(D)** Bar graph showing the fold enrichment of GO biological process co-annotation among gene pairs that showed positive interactions and/or suppression interactions defined at two different suppression score thresholds as described in the materials and methods section {material, #6370}. Analysis with and without mitochondrial genes is shown. **(E)** Box plot showing the average fraction of DepMap cell lines that depend on the indicated groups of HAP1 essential genes for viability. Numbers of essential genes tested in each group are indicated. Analysis is based on all essential genes, including essential genes with mitochondrial-related functions. Text colors relate to node color for suppression interactions shown in Fig. 6.

figure S15

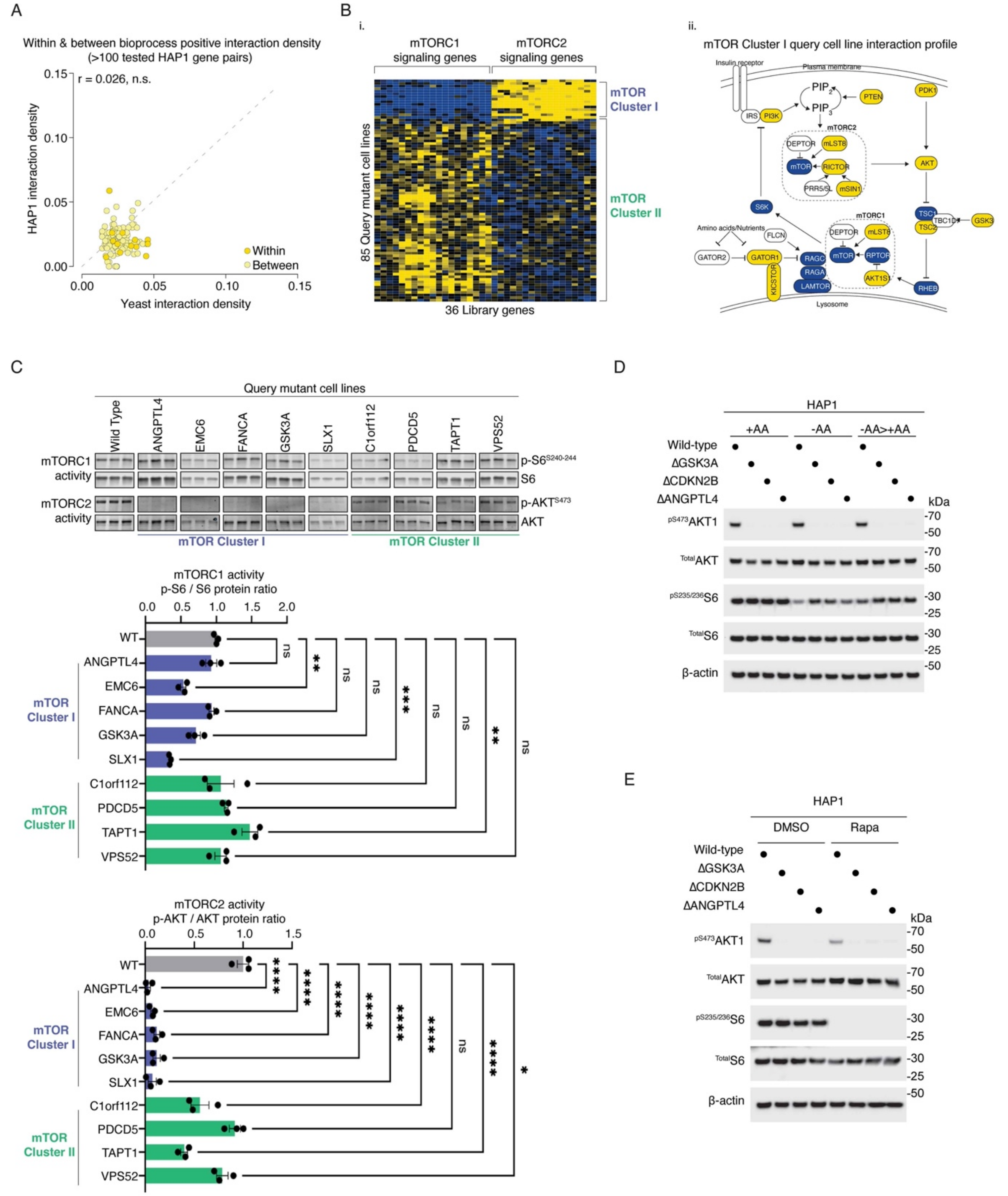

**Fig. S15. Query-driven mTOR signaling effects. (A)** Scatter plot of human (qGI > 0.3, FDR <0.1) and yeast (SGA score > 0.08, *P* <0.05) positive genetic interaction densities that occur within and between corresponding bioprocesses. **(B)** (i) Heatmap depicting hierarchically clustered genetic interaction matrix comprising 85 different query gene mutant cell lines that show coherent negative (blue) and positive (yellow) genetic interactions with library genes involved in mTORC1 and mTORC2 signaling. This analysis identified two inverse patterns of genetic interactions involving library genes with roles in mTORC1 and mTORC2 signaling. The first pattern, mTOR Cluster I, comprised 17 query mutant cell lines that showed strong negative interactions with mTORC1 signaling genes and many strong positive interactions with mTORC2 signaling genes. (ii) Schematic of mTORC1 and mTORC2 signaling pathways. Library genes are colored based on their genetic interactions with query genes belonging to mTOR Cluster I. **(C)** Immunoblots and related quantitation showing phospho-AKT1$^{S473}$ and phospho-AKT$^{S240/244}$ effects under baseline growth conditions in the indicated HAP1 queries representative of mTOR cluster 1 or mTOR cluster II signatures. Blue bars indicate mTOR cluster I and green bars indicate mTOR cluster II. * indicates level of statistical significance (* $P < 0.05$, ** $P < 0.01$, *** $P < 0.001$, **** $P < 0.0001$). **(D-E)** Immunoblots depicting phospho-AKT1$^{S473}$ and phospho-AKT$^{S6235/236}$ effects following amino acid starvation and starvation>stimulation (D) and rapamycin treatment (E) in the indicated HAP1 queries representative of mTOR cluster II signatures.

Query mutant cell lines belonging to mTOR Cluster I appeared to have normal mTORC1 activity because, in both standard growth conditions and in response to either amino acid starvation or rapamycin treatment, wild type levels of a phosphorylated mTORC1 substrate, p-S6, were observed for most mTOR Cluster I query mutants. Conversely, mTORC2 signaling was severely compromised in these same mutants, as indicated by reduced levels of a phosphorylated mTORC2 substrate protein, S473-phosphorylated AKT. Thus, mTOR cluster I positive interactions may indicate that additional perturbation of mTORC2 signaling pathway genes does not result in a more severe fitness defect in query mutant cell lines that already lack mTORC2 activity. However, a subset of mTOR Cluster I positive interactions could be classified as genetic suppression (~19% ,26/139, P $< 4.1 \times 10^{-10}$, hypergeometric)(File S17), identifying genes whose LOF may improve the fitness of cells with reduced mTORC2 activity. Nonetheless, these findings suggest that the fitness of mTOR Cluster I query mutants are more dependent on mTORC1 signaling relative to mTORC2 signaling. These results are discussed in more detail in the Supplementary text {material, #6370}.

figure S16

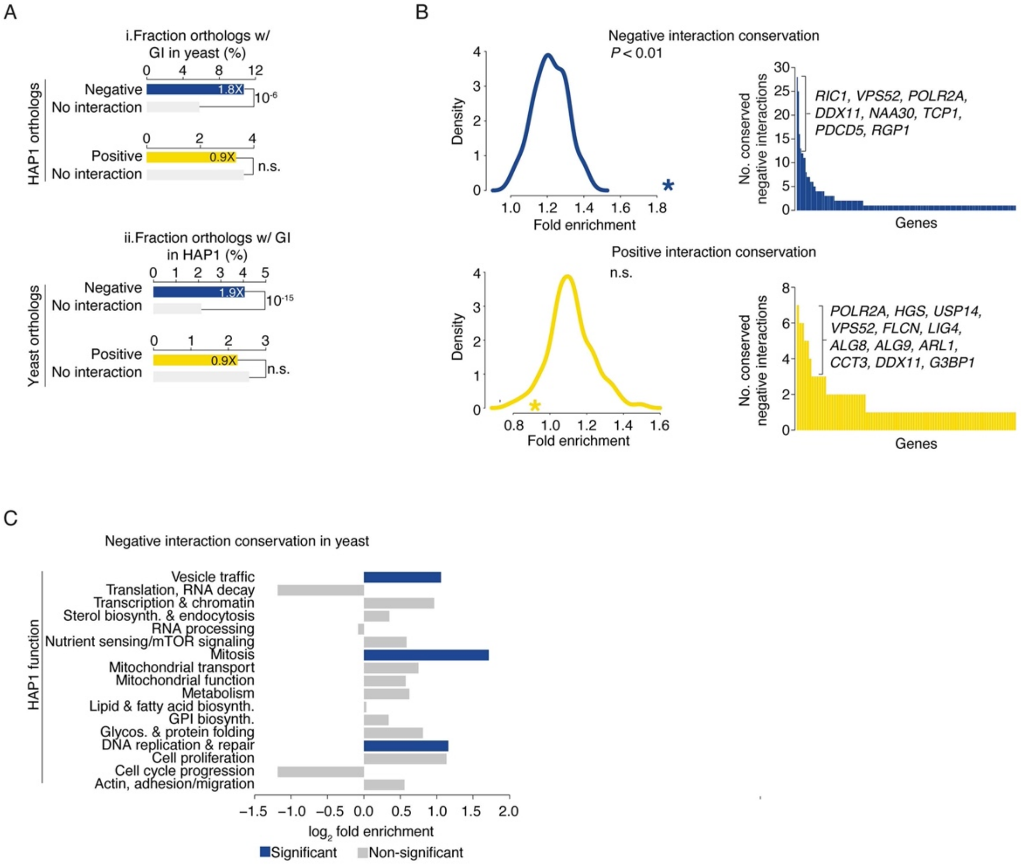

**Fig. S16. Genetic interaction conservation. (A)**(i) Bar graph showing enrichment for negative (blue) and positive (yellow) genetic interactions in yeast among conserved gene pairs scored as negative or positive interactions in HAP1 cells. Negative interactions in HAP1 were significantly enriched in yeast compared to other conserved gene pairs without negative interactions (grey). Positive interactions in HAP1 were not significantly enriched in yeast. (ii) Bar graph showing enrichment for negative (blue) and positive (yellow) interactions in HAP1 cells among conserved gene pairs with negative or positive genetic interaction in yeast. Negative interactions in yeast were significantly enriched in HAP1 compared to other conserved gene pairs (grey bar). Positive interactions in yeast were not significantly enriched in HAP1. **(B)** Density plots showing the fold enrichment in the conservation of negative (top) and positive (bottom) genetic interactions in *S. cerevisiae* for 100 randomized HAP1 genetic interaction networks. The observed fold enrichment of the real HAP1 genetic interaction network is represented by a star. Bar graphs show the number of negative (top) and positive (bottom) genetic interactions identified in both *S. cerevisiae* and HAP1 cells per human gene. Query genes that contribute many conserved negative and positive interactions are indicated. **(C)** Bar graph illustrating biological processes enriched for conserved negative genetic interactions among human-yeast orthologous gene pairs identified in HAP1 genetic interaction screens. Blue bars indicate significant enrichment.

figure S17

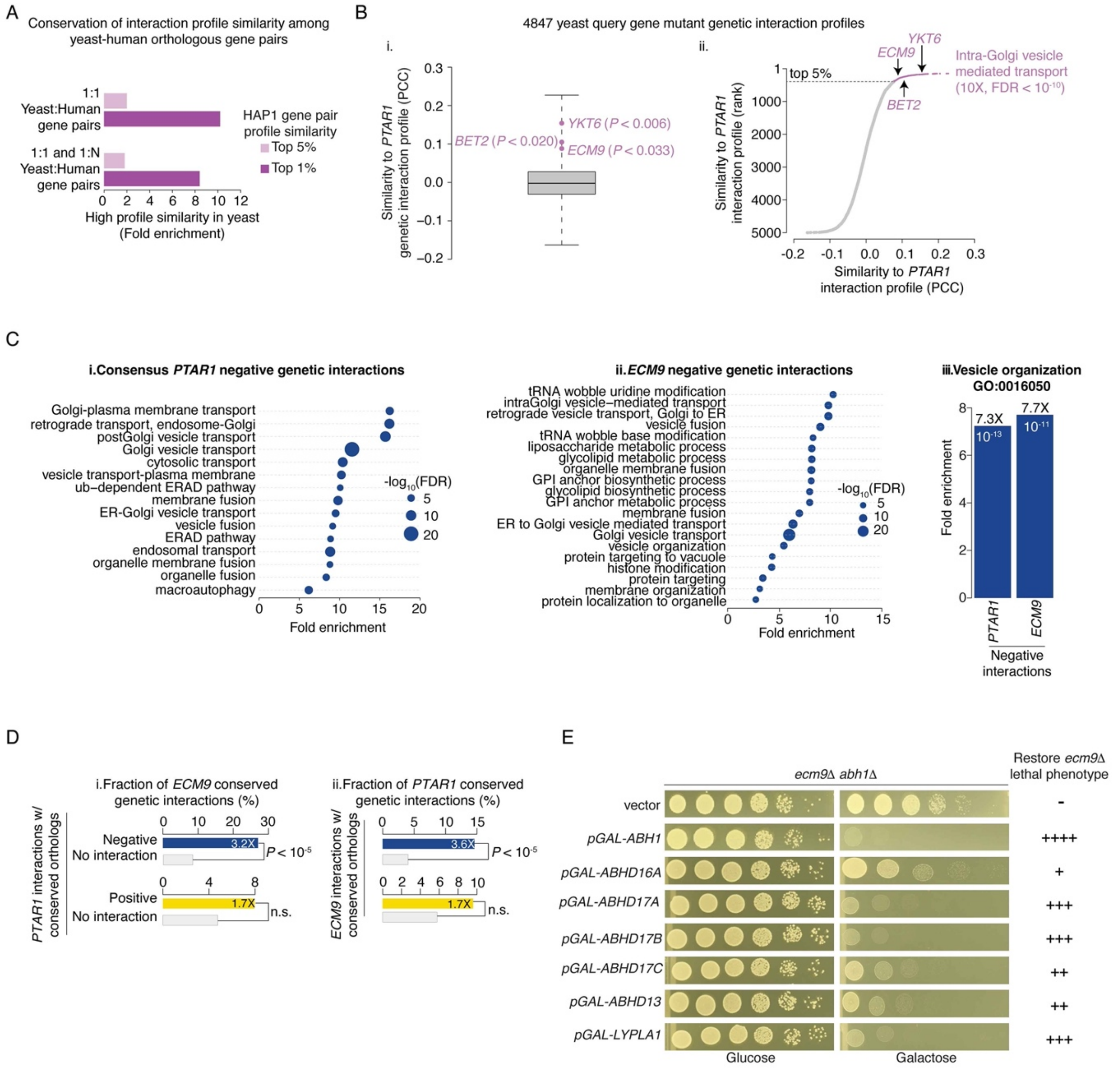

**Fig. S17. Conservation of *PTAR1/ECM9* genetic interactions. (A)** Bar graph illustrating that human gene pairs with conserved yeast orthologs (1:1 and 1 yeast: N human ortholog pairs) that share highly similar genetic interaction profile in the HAP1 network are enriched for gene pairs that also show high profile similarity in the global yeast genetic network. Gene pairs with high genetic interaction profile similarity were defined as those in the top 5% (light purple) or top 1% (dark purple) of gene pairs with the highest Pearson's Correlation coefficient similarity values in the corresponding HAP1 and yeast networks. **(B)** (i) Box plot illustrating the distribution of genetic interaction profile similarity (PCC) of 4847 yeast query genes with the HAP1 genetic interaction profile mapped for a *PTAR1* query cell line. The similarity of the human *PTAR1* genetic interaction profile with yeast *YKT6, BET2* and *ECM9* query gene genetic interaction profiles are indicated. (ii) Graph of 4847 yeast query genes plotted as a ranked order of the similarity (PCC) of their yeast genetic interaction profiles to that of human *PTAR1*. The top 5% of yeast query genes (~240) with the highest similarity to the *PTAR1* genetic interaction profile were enriched for the indicated GO bioprocess term (hypergeometric test,

Benjamini-Hochberg-corrected). **(C)** (i) GO biological process terms enriched among human genes that show a negative interaction with a *PTAR1* query gene and (ii) yeast genes that show a negative interaction with an *ECM9* query gene. (iii) *PTAR1* and *ECM9* negative interactions are both enriched for genes annotated to the GO bioprocess term, Vesicle Organization. **(D)** (i) Bar graph illustrating enrichment for negative (blue) and positive (yellow) interactions with yeast *ECM9* among conserved gene pairs that showed a negative or positive genetic interaction with human *PTAR1*. Negative interactions with human *PTAR1* were significantly enriched for genes with conserved orthologs that showed negative interactions with yeast *ECM9*, relative to all other conserved gene pairs that were tested for interactions did not show a negative interaction with *PTAR1* (grey bar). Positive interactions with *PTAR1* were not significantly enriched for positive interactions with yeast *ECM9* (grey bar). (ii) Bar graph showing enrichment for negative (blue) and positive (yellow) interactions with human *PTAR1* among conserved gene pairs that showed a negative or positive genetic interaction with yeast *ECM9*. Negative interactions with yeast *ECM9* were significantly enriched for negative interactions with human *PTAR1*, but positive interaction with yeast *ECM9* showed no significant enrichment with human *PTAR1*. **(E)** Serial dilution growth assays of a yeast *ecm9Δ abh1Δ* double mutant carrying the indicated galactose-inducible gene expression plasmid in glucose- (repressive condition) or galactose (inducible condition)-containing medium. The ability of each galactose-inducible gene to rescue the loss of *ABH1* and restore the lethal phenotype of an *ecm9Δ* deletion mutant is shown.

figure S18

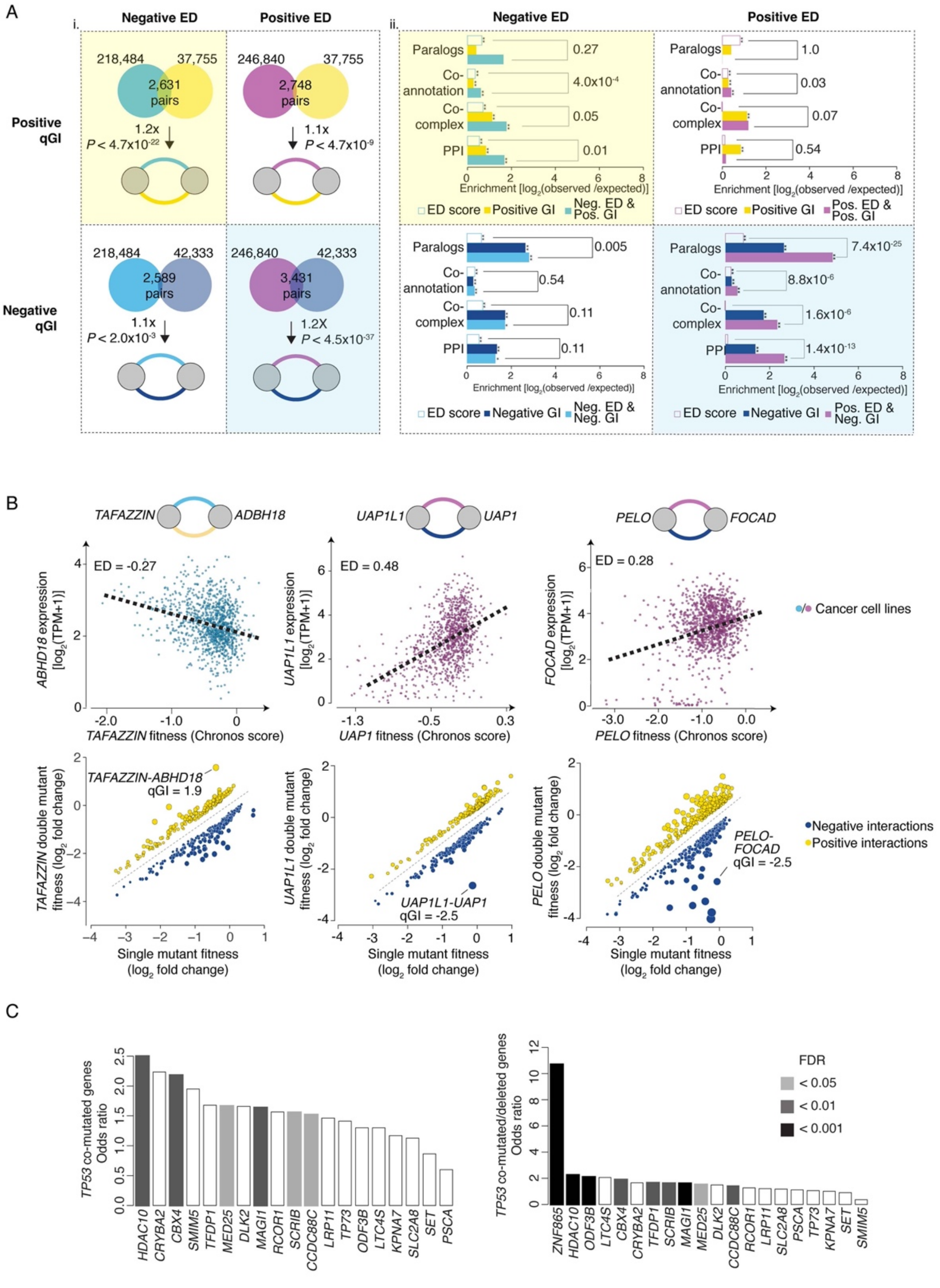

**Fig. S18. A relationship between expression dependency in cancer cell lines and HAP1 genetic interactions. (A)** (i) Overlap of gene pairs associated with all possible ED and qGI score combinations. (ii) Enrichment for indicated functional standards among gene pairs showing specific combinations of significant ED and qGI scores. Shaded regions indicate that gene pairs with a negative ED and positive qGI (yellow), or a positive ED and a negative qGI (blue), score combinations that share the most significant gene pair overlap. **(B)** Scatter plots illustrating the relationship between *TAFFAZIN* single mutant fitness and *ABHD18 expression*, *UAP1* single mutant fitness and *UAP1L1* expression, and *PELO* single mutant fitness and *FOCAD* expression, across a panel of DepMap cancer cell lines (22Q4). Regression lines (black dashed lines) indicate either a negative ED score & a positive genetic interaction score (qGI), or a positive ED score & a negative qGI score. Scatter plots illustrating significant negative and positive genetic interactions (|qGI| > 0.3, FDR < 0.1) for *TAFFAZIN, UAP1L1,* and *PELO* query genes. *TAFFAZIN-ABDH18, UAP1L-UAP1*, and *PELO-FOCAD* genetic interactions and qGI scores are indicated. **(C)** TCGA Pan-cancer analysis of co-occurring mutations with TP53. Genes with positive interactions with TP53 and significant ED-scores were evaluated for enrichment for co-occurring mutations across all tumor types in TCGA. The odds ratio reflects enrichment in co-occurrence. The left bar plot shows results based on predicted damaging mutations. The right bar plot shows results of both damaging mutations and deletion events. Shading indicates statistical significance (FDR) as tested by a Fisher exact test with Benjamini-Hochberg multiple hypothesis correction.

figure S19

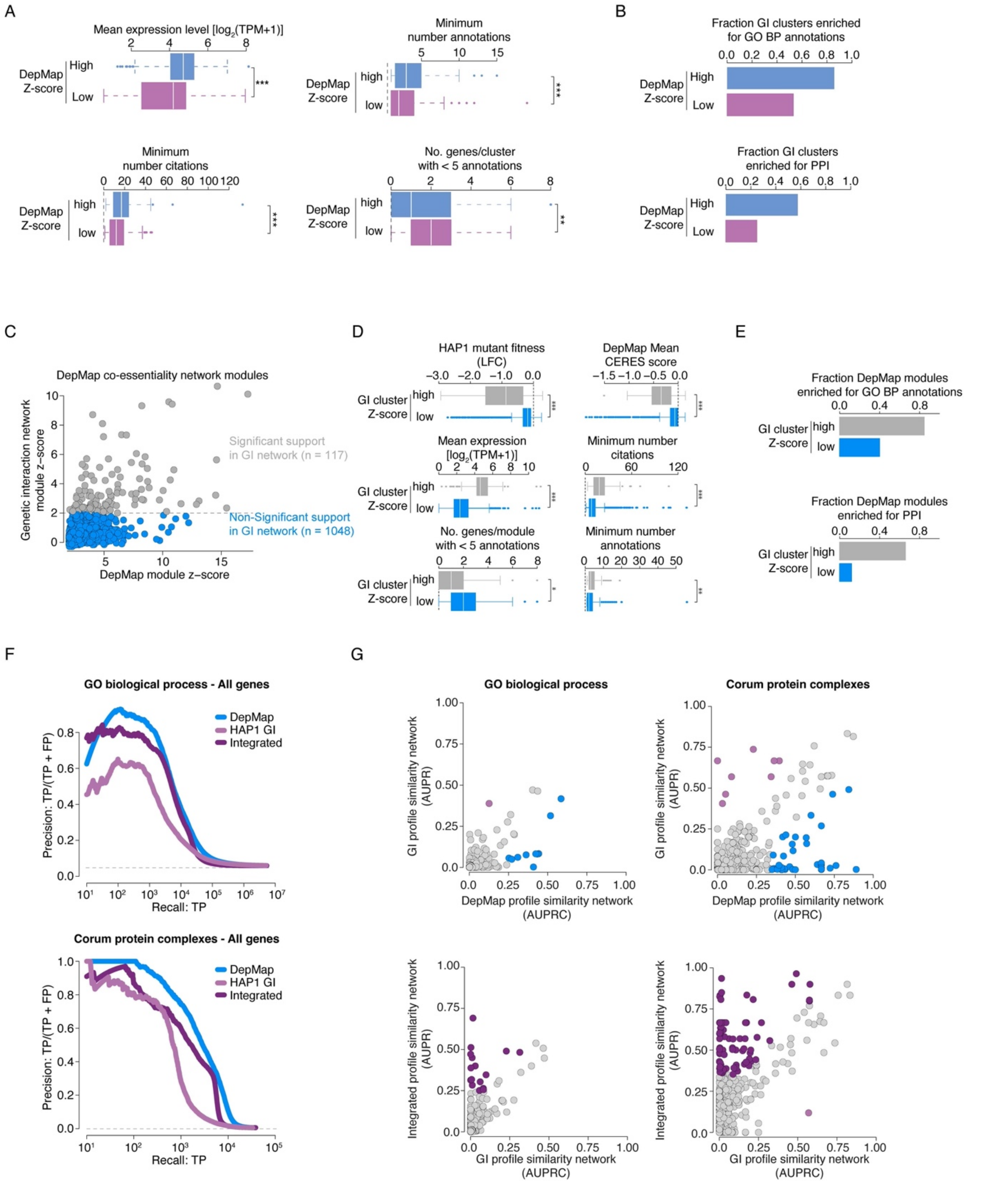

**Fig. S19. An integrated functional network based on genetic interaction and co-essentiality profiles. (A)** Box plots showing the relationship between features and genes in significant clusters or modules derived from the genetic interaction profile similarity network that either show highly correlated DepMap co-essentiality profiles (blue bars) or do not show highly correlated co-essentiality profiles (purple bars). (**B**) Enrichment GO biological process co-annotation and PPIs among genes that belong to the same genetic network-derived clusters that are also supported by modules derived from the DepMap co-essentiality network (blue bars), or among genes that cluster together to form modules based on similar genetic interaction profiles alone (purple bars). **(C)** Scatter plot of Z-scores associated with gene modules identified from the complete DepMap co-essentiality network. Modules with significant genetic interaction profile similarity in HAP1 are shown in grey, while those without significant similarity are blue. The grey dotted line indicates the Z-score threshold for significant genetic interaction profile similarity. **(D)** Box plots showing the relationship with indicated features belonging to significant modules derived from the DepMap co-essentiality network that share similar genetic interaction profiles (grey bars) or that do not have strongly correlated genetic interaction profiles (blue bars). **(E)** Bar plot illustrating the fraction of DepMap co-essentiality gene clusters whose members are enriched for the same GO-BP terms or PPIs. The fractions of enriched clusters uniquely identified in the DepMap co-essentiality network (blue bars) or DepMap-derived clusters comprising genes that also share highly similar genetic interaction profiles (grey bars) are shown. **(F)** Precision-recall plots for genes with similar DepMap co-essentiality profiles (blue), genetic interaction profiles (light purple) or integrated profiles (dark purple). TP involve gene pairs co-annotated to a gold standard set of GO-BP terms (top panel) or CORUM complexes (bottom panel). Grey dashed line represents background co-annotation rates. All genes, including those with mitochondrial-related functions, were included in the analyses. The performance of DepMap co-essentiality profiles (blue) is predominantly driven by profile similarity between mitochondrial genes {Rahman, 2021 #5666}. Exclusion of genes with mitochondrial-related functions, as shown in Fig. 9F and elsewhere {Rahman, 2021 #5666}, results in decreased precision-recall of DepMap co-essentiality profiles, indicating that these mitochondrial gene relationships remain a dominant signal in this analysis. **(G)** Comparison of individual GO-BPs or CORUM protein complexes captured by DepMAP (blue nodes), the genetic interaction profile similarity network (light purple nodes), or the integrated network (dark purple nodes). Nodes represent the genes annotated to a specific GO-BP term or a CORUM complexes. Axes show AUPRC values (see Methods). The diagonal indicates equivalent performance (grey nodes). Colored nodes above or below the diagonal represent GO-BP terms and CORUM complexes whose members show stronger profile similarity and thus, cluster together more tightly in the indicated network.

figure S20

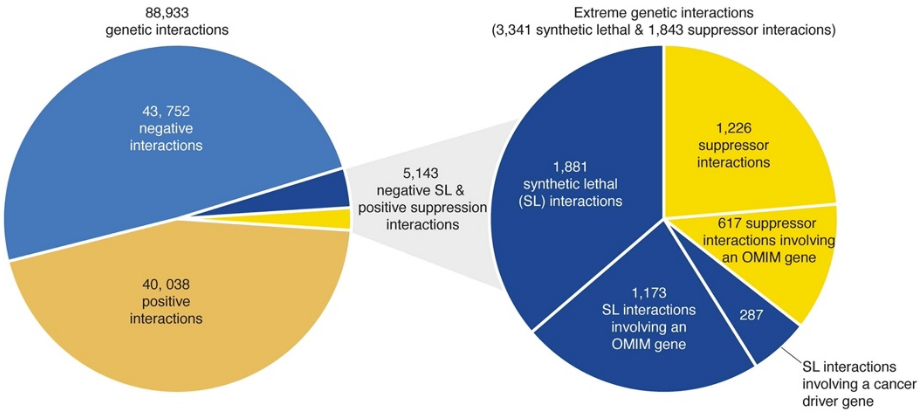

**Fig. S20. Extreme negative synthetic lethal and positive suppression interactions.** Summary of total genetic interactions identified in this study and the subset of extreme synthetic lethal or genetic suppression interactions. Negative interaction portions are labeled in blue, and positive interaction portions are labeled in yellow, with the subset of extreme negative synthetic lethal labeled in dark blue, and positive suppression interactions labeled in dark yellow. Extreme genetic interactions are further broken down (right pie chart) based on overlap with Mendelian disease genes (as identified in the OMIM database) or with characterized cancer driver genes (i.e. tumor suppressor genes){Sondka, 2023 #6285}. Extreme synthetic lethal interactions correspond to HAP1 expressed, nonessential gene pairs with a negative genetic interaction (qGI < 0, FDR < 0.1) where the single mutant fitness of the library gene (LFC) was > -0.5 and the corresponding double mutant fitness was < -1.0. Suppressor interactions were defined as gene pairs with a suppressor score > 0.5.
